# AGROtools: A comprehensive modular cloning toolset for *Agrobacterium* strain engineering

**DOI:** 10.64898/2026.09.29.755435

**Authors:** Chris Willig, Zachary Heinhold, Bethani M. Sutliff, Emily Brouwers, Colette Richter, Jay Adams, Tanner Whiting, James Warner, Phuoc Minh Duc Le, Grace Norman, Sarah Khan, Jeffrey C. Anderson, Greg S. Goralogia, Steven H. Strauss

## Abstract

*Agrobacterium*-mediated transformation is a cornerstone of plant genetic engineering, yet many crops remain recalcitrant to transformation and its efficiency is highly species- and genotype-dependent. The limited diversity of disarmed agrobacterial strains available for biotechnology, and the technically challenging workflows typically required for strain modification, constrain efforts to develop improved plant transformation platforms.

Here, we describe the development of AGROtools—a versatile, expandable, and user-friendly modular cloning plasmid collection comprising more than 75 individual components, that is designed to simplify genetic engineering of agrobacteria. Constructs assembled with this toolkit couple visual reporters with selectable and counterselectable markers to facilitate identification of recombinant clones, thereby reducing reliance on PCR-based screening. We constructed and tested reporter cassettes that are functional across diverse agrobacterial strains and genomic contexts. By linking a visual reporter combinatorially with various antibiotic resistance markers and orthogonal site-specific recombinase recognition sequences, AGROtools can enable marker removal and recycling for up to four successive genomic modifications.

We demonstrate the utility of AGROtools by disarming 10 novel wild-type strains, and accomplishing serial modifications within individual strain backgrounds. In addition to conventional *recA*-mediated homologous recombination, the collection incorporates components for applying contemporary strain engineering approaches, including CRISPR/dCas9 base editing, INTEGRATE-mediated targeted insertion, and phage-derived recombineering systems. Constructs can be assembled and deployed rapidly, enabling isolation of strains carrying an intended modification in less than two weeks. AGROtools provides a broadly applicable resource for accelerating strain development and genetic studies in agrobacteria, with potential to advance plant biotechnology by aiding improvement of disarmed strains.

## Introduction

Isolates of *Agrobacterium tumefaciens* and some closely related species within the Rhizobiaceae belonging to other genera (here collectively referred to as “agrobacteria,” following Weisberg *et al*., 2020), possess the capacity to transfer a segment of DNA into plant host genomes. They are a foundational resource in crop biotechnology. Though other methods of gene delivery into target plants are regularly applied in many species, including particle bombardment and PEG-mediated transfection, *Agrobacterium*-mediated transformation (AMT) remains the most widely used modality for producing transgenic and gene-edited crop lines (Hwang *et al*., 2017).

Shortly after the mechanism of *Agrobacterium* virulence was elucidated in the late 1970s to early 1980s, a few research groups began generating disarmed strains (Chilton, 2001). Strains are considered disarmed following the deletion of transferred DNA (“T-DNA”) segments residing on the tumor inducing (Ti) or root inducing (Ri) virulence plasmid, while retaining the set of 30 or so genes encoded elsewhere on the virulence plasmid (the *vir* regulon) which mechanistically facilitate DNA transfer (Lee and Gelvin, 2008). Such strains serve as useful tools for plant transformation by enabling the delivery of introduced recombinant sequences (flanked by T-DNA border repeats) into plant cell lines. Disarmed strains developed in the early years of plant transformation such as GV3101(pMP90), LBA4404, and EHA101 (along with its derivatives EHA105 and AGL1) remain by far the most popular vehicles for AMT even in newly developed plant transformation pipelines (De Saeger *et al*., 2021). These, along with only a few other strain backgrounds, are core parts of the conventional AMT toolbox.

Strains historically classified as *Agrobacterium* include pathogenic strains carrying Ti or Ri (“virulence”) plasmids, as well as closely related non-pathogenic strains. This group is highly diverse and polyphyletic, with several lineages reclassified into other genera including *Rhizobium* and *Allorhizobium* (Flores-Félix *et al*., 2020). Comparative genomic and phylogenetic analyses by (Weisberg *et al*., 2020) showed that plant transformation capacity evolved independently in multiple lineages within the broader agrobacteria-rhizobia complex (ARC), which also contains non-pathogenic and symbiotic lineages. Weisberg *et al*. (2020, 2021) also resolved relationships among diverse virulence plasmids, which have undergone extensive horizontal transfer and recombination, identifying eleven Ti plasmid types and three Ri plasmid types, each with additional subtypes in some cases.

At a group level, agrobacteria have a wide plant host-range, with over 600 plant species known to be infected under natural conditions (Guo *et al*., 2019). However, among different strain lineages diversity and specialization have been documented with respect to plant clades, species, or varieties (Anderson and Moore, 1979; Perry and Kado, 1982). Considering the naturally existing diversity among agrobacteria, the set of disarmed conventional AMT strains mentioned above is derived from a remarkably narrow slice of wild-type progenitors. GV3101(pMP90), EHA101, EHA105, and AGL1 are all derived from the C58 wild-type background (with the latter three all carrying a disarmed variant of the virulence plasmid from strain Bo542), while LBA4404 is descended from the Ach5 wild-type (Pennetti *et al*., 2025). Many plant species and varieties of economic and ecological significance remain recalcitrant to AMT, despite advancements in regeneration enhancing technology such as the use of morphogenic regulator genes and peptide hormones (Youngstrom *et al*., 2025). One potential way to make breakthroughs in transformation of some of these currently recalcitrant plants could be to leverage natural variation with respect to host range among agrobacteria strains by generating novel disarmed strains with diverse chromosomal backgrounds and virulence plasmid types.

The techniques available to early *Agrobacterium* researchers for manipulating strains relied on either transposon insertional mutagenesis or on the genome repair mechanism of homologous recombination (Klee *et al*., 1987). Since the former occurs randomly—requiring labor-intensive screening to obtain a desired modification—the latter method has been most commonly applied for disarmament. Homologous recombination (HR) is effectuated by a complex of proteins encoded by endogenous genes, of which *recA* is the key component in bacteria. Double crossover HR is a technique relying on incorporation of two discrete DNA fragments called “homology arms” (HA), which are often 1 kilobase or larger and derived from a strain genome at each end of a target sequence segment, into a recombinant plasmid construct. This construct can then undergo two subsequent recombination events following introduction into a strain background, resulting in replacement of the target sequence with a cargo positioned between the two HAs in the construct. In addition to its use for disarmament, this is also the primary method used to generate mutants for basic functional genetics studies in agrobacteria (Lai *et al*., 2025; Onyeziri *et al*., 2022; Williams *et al*., 2022; Zhao and Binns, 2011). However, modification through HR can be difficult to carry out in some strain backgrounds as it necessarily relies on relatively infrequent recombination events and may necessitate extensive PCR-based colony screening to identify mutant clones.

The development of CRISPR/Cas and related systems has expanded available tools for agrobacterial strain engineering to include several *recA*-independent approaches. Rodrigues *et al*. (2021) used a CRISPR/dCas9 cytidine base editor to generate gene knockouts through early stop codons, a strategy further refined by Pennetti *et al*. (2025) with variable antibiotic markers and chromoprotein reporters to facilitate colony screening. Aliu *et al*. (2022) developed the CRISPR-associated transposase system INTEGRATE for targeted genomic insertions, while Bian *et al*. (2022) developed novel recombineering systems in agrobacteria to incorporate linear DNA fragments with 80-bp homology arms for gene knockouts. Gao *et al*. (2025) iterated on the latter tools, coupling recombineering with CRISPR/Cas9-mediated double-strand cleavage to achieve a higher frequency of targeted genomic modifications. However, available plasmid resources for each of these systems vary in design and often offer limited customizability, restricting their broader application.

With the aim of empowering a wide range of sequence alterations across diverse agrobacterial strains, we designed a cloning toolkit in the vein of previously developed resources such as MoClo (Weber *et al*., 2011). Our toolkit incorporates both novel and classical strain engineering methods while prioritizing versatility, expandability, and user-friendliness. Here we describe and demonstrate the use of this suite of molecular cloning components, called “AGROtools,” that can facilitate modular assembly of constructs enabling nearly any desired genomic modification to organisms within the agrobacteria/rhizobia complex. A flow diagram representing the overall progression and elements of this project is displayed in Figure 1.

**Figure 1:**
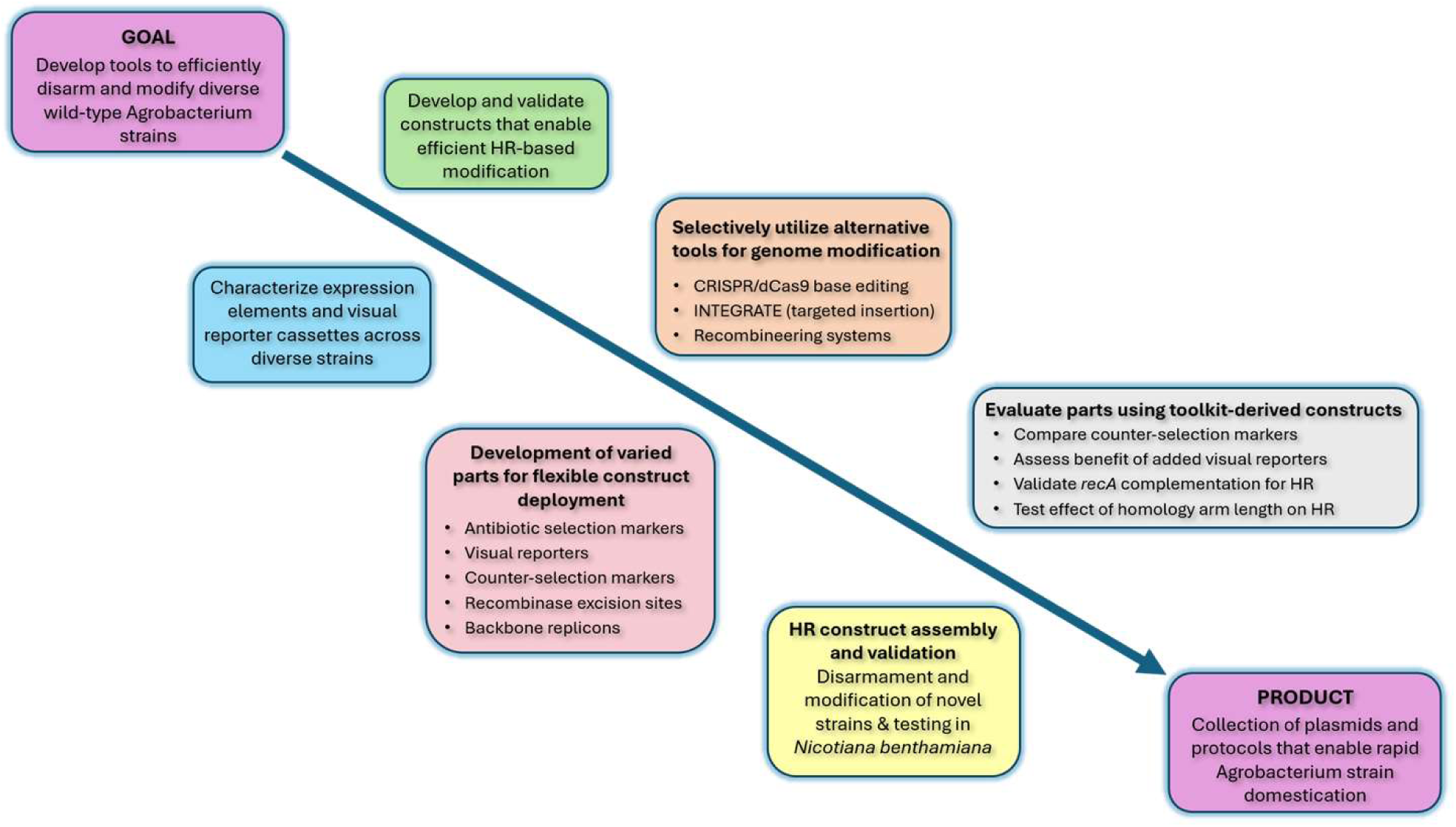
Roadmap of experimental progression in this study.

## Results and Discussion

### AGROtools design criteria

We initially sought to establish generalizable methods for efficient disarmament of diverse wild-type agrobacteria from the strain collection that was sequenced and characterized in Weisberg *et al*. (2020, 2021). However, we observed wide variation among strains with respect to their amenability to plasmid introduction, their susceptibility to various antibiotics, and other growth characteristics. Pennetti *et al*. (2025) described the use of a construct incorporating a dual paired marker design to aid in disarmament of the difficult wild-type strain background Chry5 via HR. This strategy involves coupling two different visual reporters to antibiotic selection and counterselection markers, respectively, enabling visual tracking of recombination status. Taking inspiration from this approach, we generated and deployed a similar construct for disarmament of a novel strain (*Allorhizobium vitis* str. T60/94) and observed that it greatly streamlined colony selection at each stage of the process (detailed in Figures 2 and 3). Assembly of this initial construct required a sophisticated cloning scheme using parts from many disparate nucleic acid templates, making the process difficult to repeat for custom projects. Thus, we sought to design and build a cloning toolkit to simplify the construction of these complex designs for custom engineering applications by generating a set of plasmid components designed for “off-the-shelf” use.

**Figure 2:**
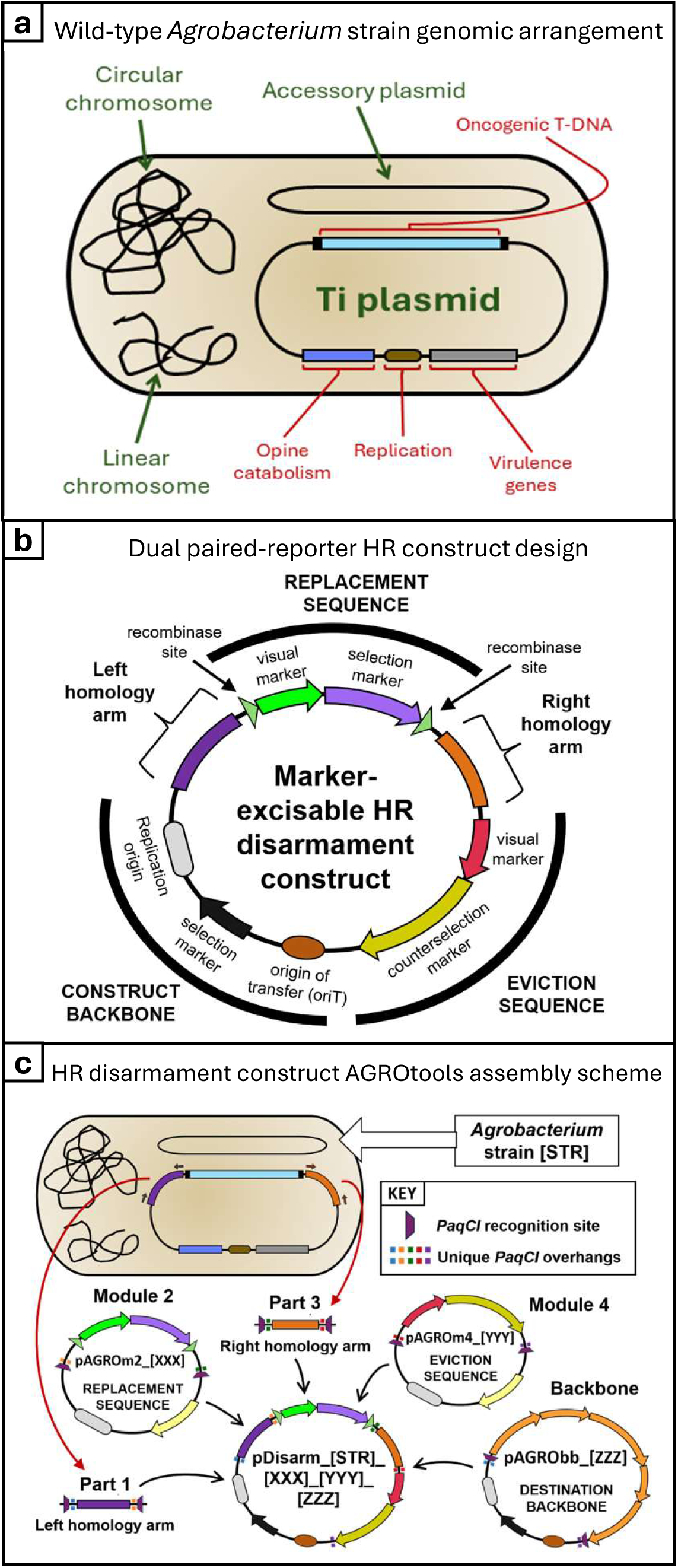
AGROtools components enable simplified construct assembly for disarmament and other genetic modifications. **a)** Cell diagram depicting a generalized genomic configuration of virulent wild-type *Agrobacterium* strains (based on strain C58) with four discrete replicons labeled in green and important features on the virulence (Ti) plasmid labeled in red. **b)** Generalized construct diagram for HR constructs employed in this study with both modular cargo segments and individual molecular parts labeled. **c)** For deletion of the native T-DNA segment (disarmament) left and right homology arms flanking the target T-DNA (purple and orange segments, respectively) are amplified by PCR (designated by curved red arrows) from the target strain genomic DNA template using two primer pairs (small block arrows). The two resulting linear fragments (labeled as Parts 1 and 3) can then serve as direct inputs, along with AGROtools components of choice from each of the module 2, module 4, and backbone classes, for construct assembly (designated by curved black arrows). Suffixes in names for individual AGROtools component plasmids denote specific molecular parts contained in each of their cargos. Generalized suffixes, shown in brackets, demonstrate how an assembled construct is named based on the inputs used to generate it.

**Figure 3:**
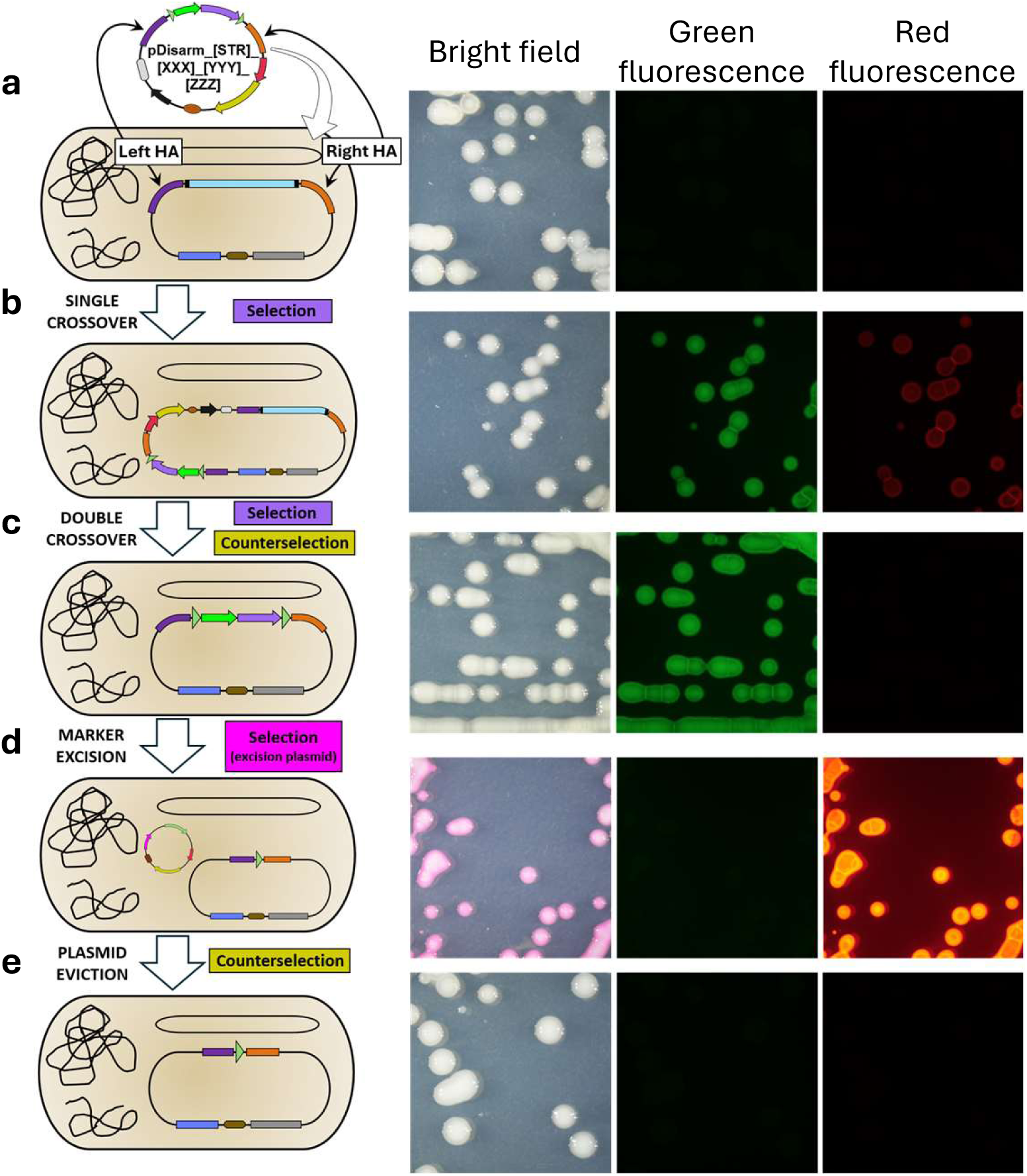
Steps in the process of HR-based modification enabled by AGROtools. An assembled disarmament construct is introduced into the target strain background **(a)**, which is placed on media selection for the corresponding marker in the replacement sequence. Single-crossover colonies **(b)** are identified by their expression of both visual reporters and are re-plated on media amended with both selection and counterselection agents to filter secondary recombinants. Double-crossover, disarmed colonies **(c)** are identified by expression of replacement sequence marker and loss of eviction sequence marker. An excision plasmid expressing a recombinase enzyme specific to the pair of recognition sites flanking the replacement sequence is then introduced into the disarmed strain background and plated on media to select only for the introduced plasmid, resulting in marker-excised colonies negative for the replacement sequence marker, but expressing the red fluorescent signal conferred by the plasmid **(d)**. Finally, colonies are re-plated on sucrose counterselection media to evict the excision plasmid, resulting in a disarmed, marker-excised strain derivative carrying only a small scar sequence with a single recombinase site in place of the original target sequence **(e)**.

### Toolkit design scheme, component descriptions, and usage

The AGROtools cloning kit was designed to facilitate highly customizable one-pot construct assembly from five discrete inputs. For a generalized construct designed to facilitate HR-based strain disarmament as depicted in Figure 2A and B, sequences of molecular parts conferring specific functions were divided into modular segments and assigned to one of five plasmid component classes (designated as modules 1 through 4 and “backbone”). These modular plasmids enable positionally ordered construct assembly directed through ligation of unique overhangs generated upon digestion by the type IIS restriction enzyme *PaqCI*. Modules 1 and 3 components can serve as container plasmids for left and right HA fragments, respectively; plasmids from module 2 and 4 classes contain “replacement sequence” and “eviction sequence” cargos, respectively; and backbone plasmid components provide the construct replicon.

Replacement sequence refers to the DNA segment positioned between the left and right HA sequences in a disarmament construct which remains at the genomic target site in place of a deleted sequence following the secondary recombination event. In our construct design, it consists of a visual reporter cassette tethered to an antibiotic selection marker, both of which are flanked by a pair of recombinase recognition sites in direct repeat orientation to enable downstream excision of the marker genes. Eviction sequence refers to the segment positioned adjacent to a construct backbone, just outside the right HA, consisting of a different visual reporter than that used in the replacement sequence, paired to a counterselection marker. This sequence is integrated along with the construct backbone into the genomic target site upon primary recombination and assists with filtering for secondary recombinant events as the parts are employed to select against cell populations still carrying them.

To generate a custom disarmament construct, a user first designs tailored left and right HA fragments from a target strain genome, which may either be inserted into module 1 and 3 entry plasmids or utilized directly as PCR amplicons. Modular plasmids from each of the remaining classes are then selected for combination with the two HA inputs in a golden gate assembly reaction (Figure 2C).

To provide a high degree of customizability in construct design, we generated many variant plasmids for each component class carrying unique combinations of functional molecular parts within their cargo sequences. Replacement sequence cargos in module 2 variants carry one of a few different visual reporters (primarily sfGFP) joined combinatorially with a selection marker conferring resistance to the antibiotics kanamycin, spectinomycin, gentamicin, or apramycin, along with one pair from set of four orthogonal recognition sites for site-specific recombinases. The varied selection markers were included in AGROtools to assist in circumnavigating potential basal or cryptic antibiotic resistance phenotypes, which have been previously documented in various wild-type strain backgrounds (Alzohairy *et al*., 2013; Lopez-Agudelo *et al*., 2026; Luo and Farrand, 1999; Pennetti *et al*., 2025). Eviction sequence cargos in module 4 variants also include a visual reporter (typically the chromoprotein eforRed is used) as well as one of three counterselection markers, and in some cases an additional cassette for ectopic expression of the recA protein (discussed below). Cargos in modules 2 and 4 were designed so that individual molecular parts are interspersed with an ordered set of unique type IIP restriction enzyme sites, allowing simple part replacement and toolkit enhancement (Figure S1). Backbone plasmid variants offer diverse replicon types belonging to different incompatibility groups (both replicative and non-replicative) in factorial combination with a set of antibiotic selection markers. Table 1 details the naming conventions used for AGROtools components, listing the specific molecular parts used for each class and how they are encoded in plasmid name suffixes.

**Table 1.**
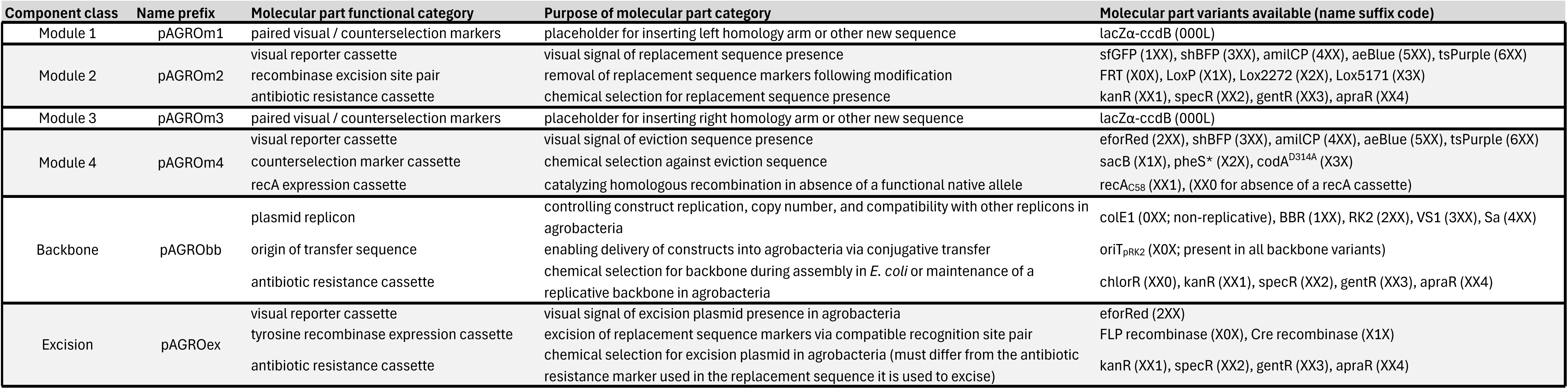
Summary of AGROtools component layout, molecular parts, and naming conventions for homologous recombination-based disarmament.

The paired visual reporter HR constructs assembled using AGROtools enable clear visual marking of agrobacterial strain colonies at each individual recombination step (Figure 3). Assuming the default design utilizing sfGFP and eforRed markers in the replacement and eviction sequences respectively, a primary recombination event resulting in whole-construct integration into the target sequence confers expression of both reporters in colonies formed 2-3 days post-introduction. A secondary recombination event causing deletion of the target sequence (typically obtained by applying active counterselection against the eviction sequence) results in loss of red fluorescence, but retention of green fluorescence. Next, an AGROtools plasmid of a sixth class denoted as “excision” can be introduced into the strain to remove the replacement sequence markers for recycling and potential subsequent modifications. Excision plasmids contain an expression cassette for either Cre or FLP recombinase paired with an antibiotic resistance marker. The plasmids, which are replicative in agrobacteria, also carry eforRed and *sacB* (counterselection) cassettes. Colonies which are successfully marker-excised, but which carry the excision plasmid are thus positive for red fluorescence, but negative for green fluorescence. Finally, colonies formed after plating on media amended with sucrose, thereby evicting the excision plasmid, are negative for all visual reporters.

Though AGROtools was primarily designed to expedite the process of making HR-based modifications, we sought to create a maximally versatile resource to simplify construct design for all published methods for agrobacterial strain modification. We thus incorporated and modularized molecular parts for CRISPR/dCas9 cytidine base editing, INTEGRATE targeted sequence insertion, and recombineering systems into AGROtools component plasmids (Aliu *et al*., 2022; Bian *et al*., 2022; Rodrigues *et al*., 2021). Key constituents of these systems were segmented and inserted into container plasmids of module 1, 2, and 3 classes to conserve the order of these parts upon construct assembly. Further plasmid variants were made based on these components to enhance customizability or to simplify the cloning process for custom sgRNA/crRNA spacer and INTEGRATE cargo insertion. Plasmids containing the building blocks for each of these tools may be combined with any existing AGROtools module 4 and backbone components (Figure S2) to enable custom marking, eviction, or copy number tuning for these tools, offering flexibility far beyond what is currently available among plasmid resources utilizing these tools. Table S1 details the naming conventions and specific molecular parts used for AGROtools plasmids containing these alternative modification tools.

The AGROtools cloning kit, consisting of 85 individual components, will be made available either as a complete set in 96-well plate format or individually from Addgene. The experiments described below were performed throughout the course of developing the toolkit to optimize the design of molecular parts incorporated into AGROtools, to evaluate components of various functional classes, and to demonstrate the versatility, efficiency, and ease of use for employing these tools in genetic modification projects.

### Evaluation of reporter cassettes and expression elements for agrobacteria

When we initially attempted to generate an HR-based disarmament construct employing the dual paired visual-selectable marker design, we observed that neither of the sfGFP or eforRed visual reporter expression cassettes which had been PCR-amplified as assembly parts from publicly available plasmid sources (Addgene #196250 and #117837, respectively) were functional when introduced into agrobacteria (Figure 4B and C, left panels). Thus, we sought to identify operative reporter cassettes and to characterize the functionality across diverse agrobacteria of four different promoters which had been employed previously in agrobacterial construct designs (P_J23119_, P_J23100_, P_int_ [Toh *et al*., 2023], and P_Atu8024_) by pairing them combinatorially with coding sequences for these two visual reporters each with corresponding terminator sequences (T3 and T7) which were also previously validated in agrobacteria.

**Figure 4:**
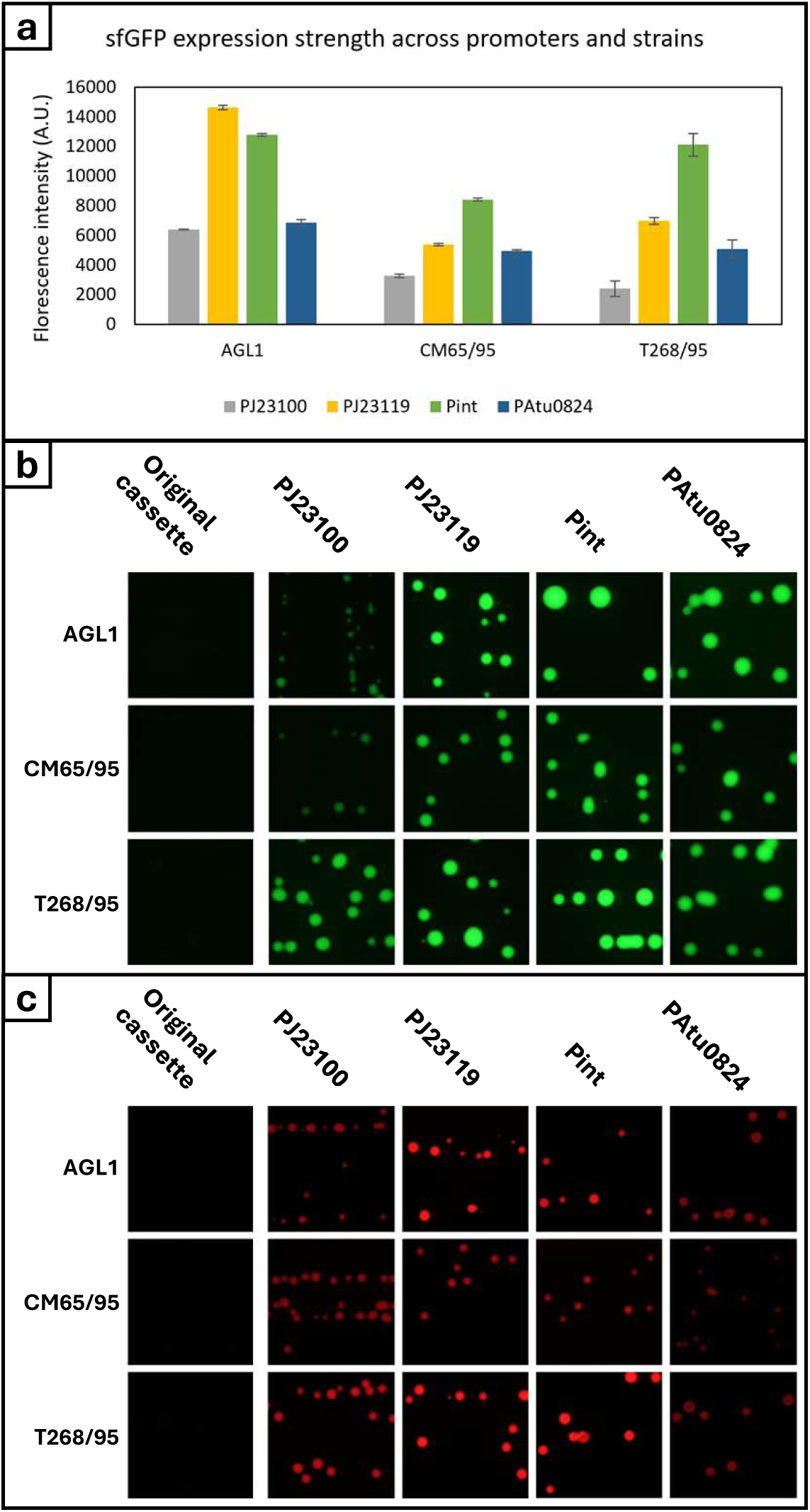
Evaluation expression elements and visual reporter cassettes across diverse agrobacteria. **a)** Quantitative comparison of fluorescence intensity conferred by sfGFP cassettes constructed using four different promoter elements, versus the expression cassette contained in plasmid originally acquired from Addgene. Each of these cassettes was screened across three agrobacterial strain backgrounds representing each of the classical biovars (BV) in this group, BV1, BV2, and BV3. Error bars represent the standard deviation across three sample wells. **b)** Images of each sfGFP expression cassette by strain background variant taken using a confocal microscope in the green fluorescence channel. **c)** Parallel cassettes to those described for sfGFP were constructed with the eforRed chromoprotein and then introduced factorially into the same strain backgrounds, to evaluate eforRed’s utility as a visual reporter. Bright field images of plates streaked with each variant are displayed.

Constructs carrying each of these cassette variants were assembled and introduced by electroporation into each of three phylogenetically diverse agrobacterial strains: AGL1 (disarmed; biovar 1 =*Agrobacterium* sp.), CM65/95 (biovar 2 =*Rhizobium rhizogenes*), and T268/95 (biovar 3 =*Allorhizobium vitis*). Quantitative fluorescence intensity data was collected for the complete set of substrains carrying sfGFP reporter cassette variants using a 96-well fluorescent plate reader. This data is displayed in Figure 4A. Streaked plates imaged with a fluorescence microscope using the corresponding filter set are shown in Figure 4B and C for the complete substrain sets carrying sfGFP and eforRed, respectively.

The P_int_ promoter conferred the highest expression out of the set in two out of the three strain backgrounds (excluding AGL1, where it was slightly outperformed by P_J23119_). P_J23100_, being a weaker variant of its synthetic series counterpart P_J23119_, conferred the lowest levels of expression in each strain, while the native *A. tumefaciens* promoter P_Atu8024_ provided moderate expression levels slightly above those of P_J23100_. We also observed overall that the fluorescent signal of sfGFP was consistently stronger under our detection system than that of eforRed across all strains. Thus, we chose to utilize the stronger on average P_int_ promoter to drive eforRed and P_J23119_ to drive sfGFP in the visual reporter cassettes incorporated as standard parts in eviction and replacement sequence cargos, respectively, for our toolkit design going forward. We also evaluated additional chromoprotein visual reporters in these three strains (not shown), which generally displayed more variable expression resulting in higher ambiguity for colony detection than when screening for sfGFP and eforRed fluorescence. Nevertheless, we generated both module 2 and module 4 variants containing expression cassettes for each of these alternative visual reporters as they may be useful for researchers without easy access to a means of visualizing the fluorescent reporters. P_J23100_ and P_Atu8024_ were used to drive expression of other molecular parts used in AGROtools discussed below.

### Comparing the utility of three counterselection marker gene cassettes

The *sacB* gene (Gay *et al*., 1983), which confers sensitivity to sucrose is, to our knowledge, the only counterselection marker that has been deployed previously in agrobacteria. To enhance the versatility of our toolkit, we developed cassettes for two additional counterselection markers demonstrated to function in other bacterial species and evaluated their utility versus that of *sacB* in filtering for secondary recombinant clones. The candidate markers we selected were *pheS*\*, a mutant phenylalanine-tRNA synthetase enabling the lethal incorporation of the Phe analog 4-chlorophenylalanline (4-CP) into proteins (Gao *et al*., 2022; Miyazaki, 2015; Wang *et al*., 2018), and *codA*^D314A^, a mutant form of a cytosine deaminase catalyzing conversion of 5-fluorocytosine (5-FC) into the toxic 5-fluorouracil (5-FU) (Zeng *et al*., 2015). Cassettes were designed using open reading frames codon-optimized for rhizobia, which were placed under the control of previously validated expression elements (P_J23100_ promoter and the rrnB T1 terminator). Each cassette was incorporated into eviction sequence cargos within module 4 variant plasmids, each of which was then utilized in the assembly of three parallel disarmament constructs targeting strain C58.

These constructs were introduced in parallel into wild-type C58 cells to obtain a series of primary recombinant events carrying each of the three counterselection markers. Independent clones of each (along with secondary recombinant clones lacking any counterselection marker) were grown in liquid cultures, adjusted to equivalent densities, and serially diluted in a 96-well plate setup by 10-fold increments down to 10^-7^. A 96-pin replicator tool (VCP Scientific, Inc.; San Diego, CA) was then used to transfer this culture array onto a series of media plates amended with each of the corresponding counterselection agents at varied concentrations. Control plates lacking any counterselection agent were also used (Figure 5).

**Figure 5:**
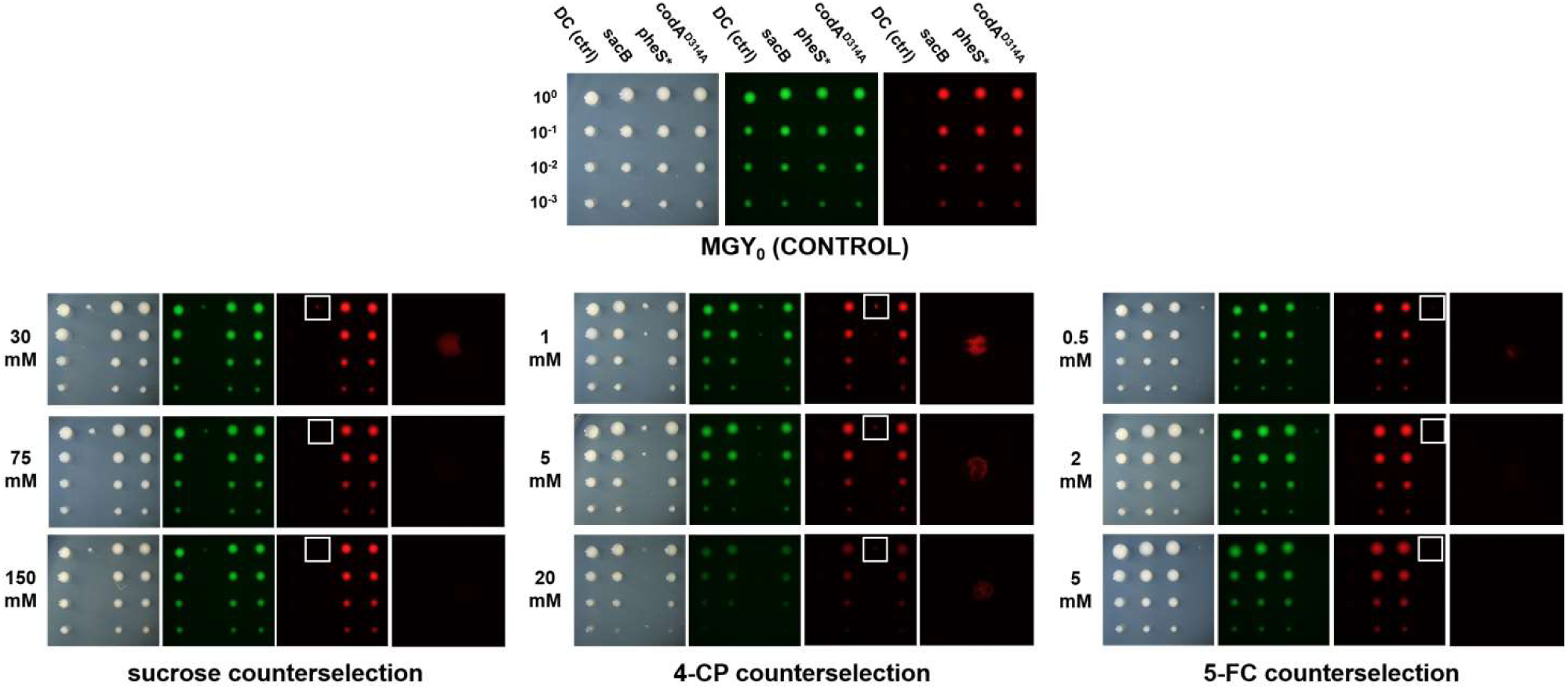
Comparison of counterselection marker efficiency. A series of single-crossover recombinant derivatives were generated from strain C58 by introducing parallel disarmament constructs carrying cassettes for each of the three counterselection markers *sacB*, *pheS*\*, and *codA*^D314A^, along with a double-crossover derivative (DC) used as a control, were serially diluted in 10-fold increments and then transferred onto a set of plates containing media amended with each of the three corresponding counterselection agents at three levels. Images display representative replicate blocks of each derivative type at four inoculum densities on each media formulation after 2 days of growth, photographed under (from left to right) bright field, green fluorescent, and red fluorescent filter sets. Images on the far right of each panel show the magnified spots (as shown by the white square inset in the image to the left) for clones under counterselection at the highest inoculum density in the red fluorescence channel. Relative efficiency of counterselection was inferred from the degree of loss of red fluorescent signal in this plate sector.

Counterselection efficiency was inferred based on the degree to which sectors lacking red fluorescence but expressing green fluorescence (indicating secondary recombination) were present two days after plating at the pin impression site with the highest inoculum density on media containing the corresponding counterselection agent. Though red fluorescence was detected in *sacB*-containing colonies at the 30 mM concentration, sucrose appeared to confer stringent counterselection at the higher levels of 75 and 150 mM. Meanwhile, reduced but detectable red fluorescent signal remained in nearly all of the evaluated pin impression sites for cultures containing *pheS*\* at all concentrations of 4-CP tested. For clones carrying *codA*^D314A^, red fluorescence was detected at the lowest concentration of 5-FC, while a very low level was present on the medium concentration. No signal was seen at the highest level tested.

Though neither of the alternative counterselection marker cassettes appeared to outperform *sacB* in this experiment, *codA*^D314A^ was comparable in its efficiency at the higher levels tested. We observed from including *sacB* in the eviction sequence in all of the constructs used to disarm wild-type strains (discussed below) that it gave variable, apparently strain-dependent results with respect to the proportion of recovered colonies lacking red fluorescent signal following plating of primary recombinants on media amended with selective antibiotic and 5% sucrose. For strain C58, over 90% of colonies had lost red fluorescence while for other strains such as Ach5 (as discussed below), less than 2% of colonies were apparent secondary recombinants under the same conditions. It is known that one potential source of failure for *sacB* is its tendency to serve as an insertion site for the transposable element IS426 (Morton and Fuqua, 2012). Though they may not be necessary for most applications, the inclusion of *pheS*\* and *codA*^D314A^ in AGROtools enhances the options available to the user in overcoming potential obstacles to double crossover in contexts for which *sacB* performs poorly.

### Disarming novel wild-type strains

Having developed a library of component plasmids for HR constructs, we were able to rapidly generate and deploy constructs for disarmament of diverse wild-type strains (Table 2). Once assembled and verified by sequencing, constructs were introduced into their respective target strains, followed by plating on media containing the antibiotic corresponding to the selectable marker in the construct replacement sequence. Colonies expressing both green and red fluorescence were pooled and re-plated to media amended with 5% sucrose and the appropriate antibiotic. Colonies formed on this plate expressing green, but not red fluorescent signal were selected and validated as successfully disarmed by colony PCR using “flanking” primers specific to each strain. These primers anneal to sites on the target Ti/Ri plasmid lying just outboard of the left and right homology arm sequences and thus will generate a band typically between 4 and 5 kbp in length only when colonies in which the native intervening sequence has been removed are used as templates, but not in wild-type or single-crossover events since the annealing sites are positioned at too great a distance from each other (Figure 6A).

**Figure 6:**
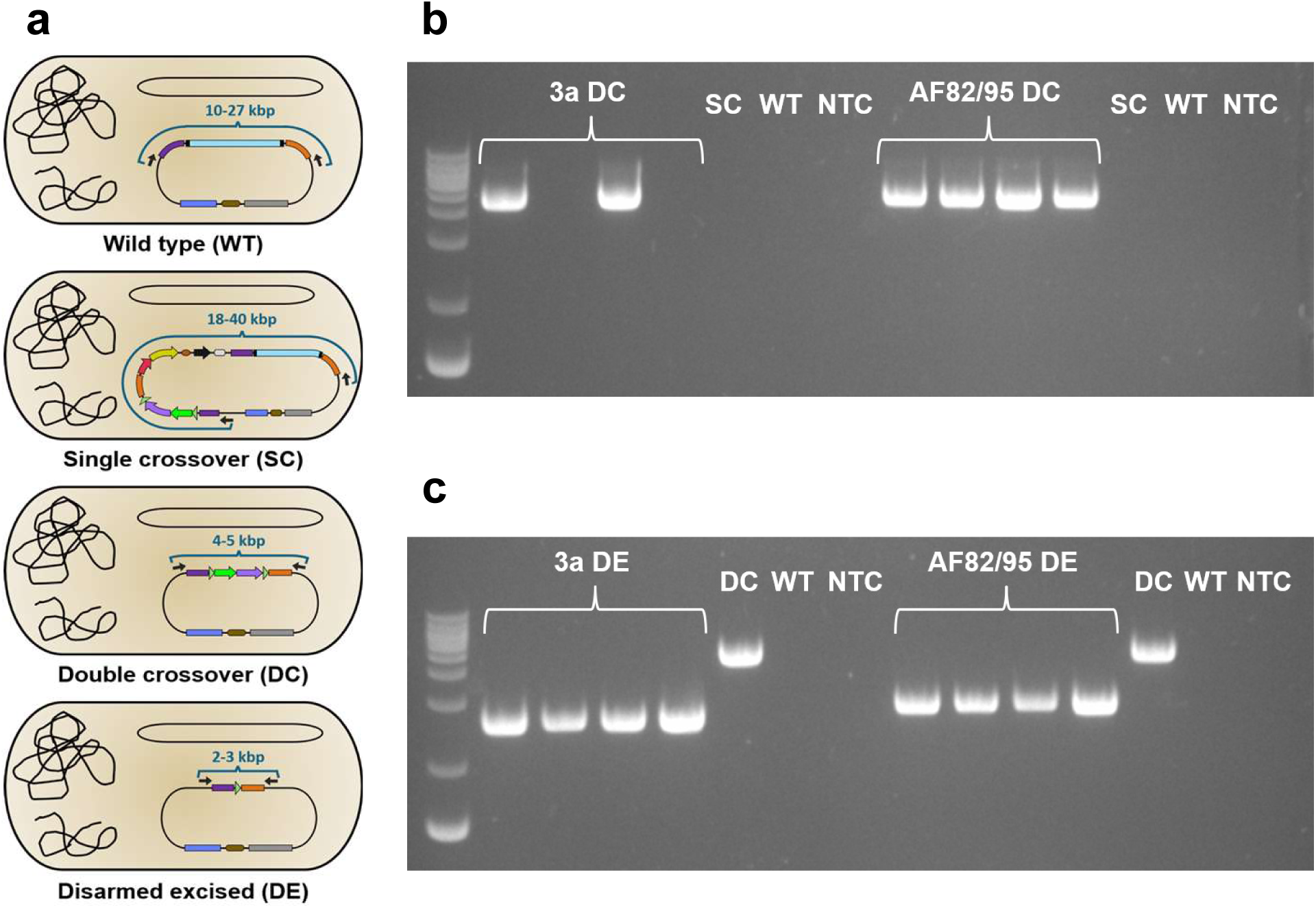
PCR validation of double crossover and marker excision. **a)** Position of annealing sites for a pair of flanking primers (charcoal arrows) in each cellular configuration. The distance between the two annealing sites is annotated in blue. **b)** Colony PCR to confirm double crossover disarmed (DC) derivatives of the strains ‘3a’ and ‘AF82/95’ with four test samples each. Single crossover (SC) and wild-type (WT) templates from each strain were used as negative controls. **c)** Colony PCR to confirm disarmed and marker-excised (DE) derivatives for ‘3a’ and ‘AF82/95’. DC templates for each were used as positive controls, while WT variants were used as negative controls. NTC = no-template control.

**Table 2.**
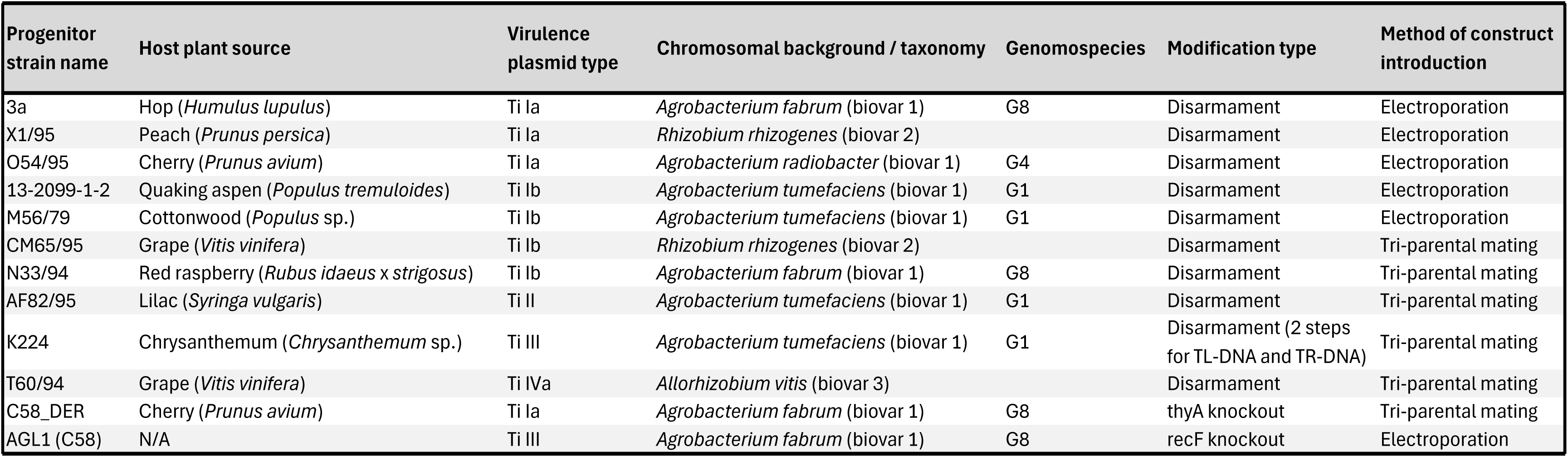
Disarmed and modified strains generated in this study.

One clone of a validated disarmed strain derivative was then transformed with an AGROtools excision plasmid to remove the residual antibiotic and visual marker pair. After selecting apparent excised colonies and re-plating them on sucrose-containing media to evict the plasmid, marker excision status was confirmed by colony PCR, again using the same flanking primer pair, which now resulted in a reduced band size. Agarose gel images of representative colony PCR runs for derivatives of both wild-type strains 3a and AF82/95 confirming double crossover recombinant and marker-excised status are displayed in Figure 6B and C, respectively. Figure S3 shows gel images demonstrating colony PCR validation of double-crossover and marker-excised status for all targeted strains.

Final disarmed and marker excised strains (designated with “_DE” following their wild-type derivative name) were whole-genome sequenced using the Oxford Nanopore platform. Analysis confirmed that all strains were disarmed and that only ∼100 bp residual sequence containing a single recombinase site remained in place of the deleted sequence. No evidence of off-target sequence insertions resulting from the modification was observed. Annotated genomic sequences for each disarmed strain discussed in this study will be made available for download via Github.

To ensure that they retained T-DNA transfer capacity, a binary plasmid carrying T-DNA conferring GFP expression in plants was introduced into a subset of the disarmed, marker excised strain backgrounds. These isolates were then utilized for agroinfiltration into *Nicotiana benthamiana* leaves, which were assessed for green fluorescence after 3 days (Figure S4). Though two of the strains gave comparatively weak signal, most conferred high levels of reporter expression apparently similar to the AGL1 comparator strain. All disarmed strains outperformed a mock control, indicating that T-DNA delivery was occurring in all cases.

A partially overlapping set of strains was used in an experiment comparing the intensity of GFP transgene expression in infiltrated *N. benthamiana* leaves conferred by these disarmed derivatives. A series of infiltrated leaf disks were measured on a 96-well plate reader. Relative to the AGL1 comparator, strain AF82/95_DE showed a significant increase in fluorescent intensity, while three strains had no detectable difference (K224_1DE_2DE, N33/94_DE, and C58_DE), and two strains significantly lower intensity (O54/95_DE and T60/94_DE) (Figure S5). Though their utility in stable transformation remains to be evaluated, these experiments suggest that some of the newly disarmed strains can confer higher levels of transgene expression than strains routinely used in plant transformation processes.

### Serial modification enabled by orthogonal recombinase sites

Our strain modification efforts benefitted from the inclusion of the orthogonal recombinase excision sites we designed across AGROtools module 2 variants. The wild-type strain K224 carries a type III Ti plasmid, which possesses two discontiguous T-DNA sequences. To avoid deleting genes in the sequence intervening them, we removed each T-DNA segment in serial operations using assembled constructs in which a pair of either *LoxP* or *Lox2272* sites were carried in the replacement sequence. Because these two different sites cannot efficiently recombine with each other, this strategy enabled a secondary marker removal step facilitated by Cre excision without catalyzing an interaction with the residual site from a previous modification cycle, which would result in the unwanted deletion taking place.

A similar strategy also enabled us to develop a *thyA* knockout mutant using HR in a previously disarmed background derived from strain C58. In addition to having had the native *recA* allele disabled through a strategy similar to that used in the development of AGL1 (Lazo *et al*., 1991), resulting in a non-excisable ampicillin resistance marker at that locus, this strain carried a remnant *LoxP* site from the T-DNA deletion. We utilized the alternative recombinase site *Lox5171* (in a construct temporarily supplying a functional *recA* allele as discussed below) to excise markers during the *thyA* deletion procedure, again to avoid unwanted recombination. Precise removal of marker genes without unwanted recombination was confirmed by whole genome sequencing of the resulting strain.

The availability of pairs of the flanking recombinase recognition sites *FRT*, *LoxP*, *Lox2272*, and *Lox5171* in AGROtools components allows for up to four discrete HR-based modifications to be made in the same background since the latter two are orthogonal to *LoxP* (Lee and Saito, 1998) and FRT is recognized by a FLP recombinase rather than Cre. Expansion of the current toolset by incorporating additional orthogonal recombinase sites that have been previously characterized (Cautereels *et al*., 2024; Langer *et al*., 2002; Missirlis *et al*., 2006; Schlake and Bode, 1994), or the utilization of base editing and INTEGRATE constructs could enable a practically unlimited number of iterative modification steps.

### Assessing the advantage of a dual paired reporter construct design

We next sought to evaluate the degree to which pairing of visual reporters with selection and counterselection markers in the AGROtools HR construct design scheme improves colony selection during each step. To investigate this, we assembled pairs of HR disarmament constructs targeting both C58 and Ach5 wild-type strains which differed in whether they contained visual reporter cassettes (and the pair of recombinase excision sites), but which were otherwise equivalent (Figure 7A). Each of these constructs were introduced into their respective target strains in parallel and in equimolar quantities by electroporation and transformed cell cultures were each plated on media amended with spectinomycin.

**Figure 7:**
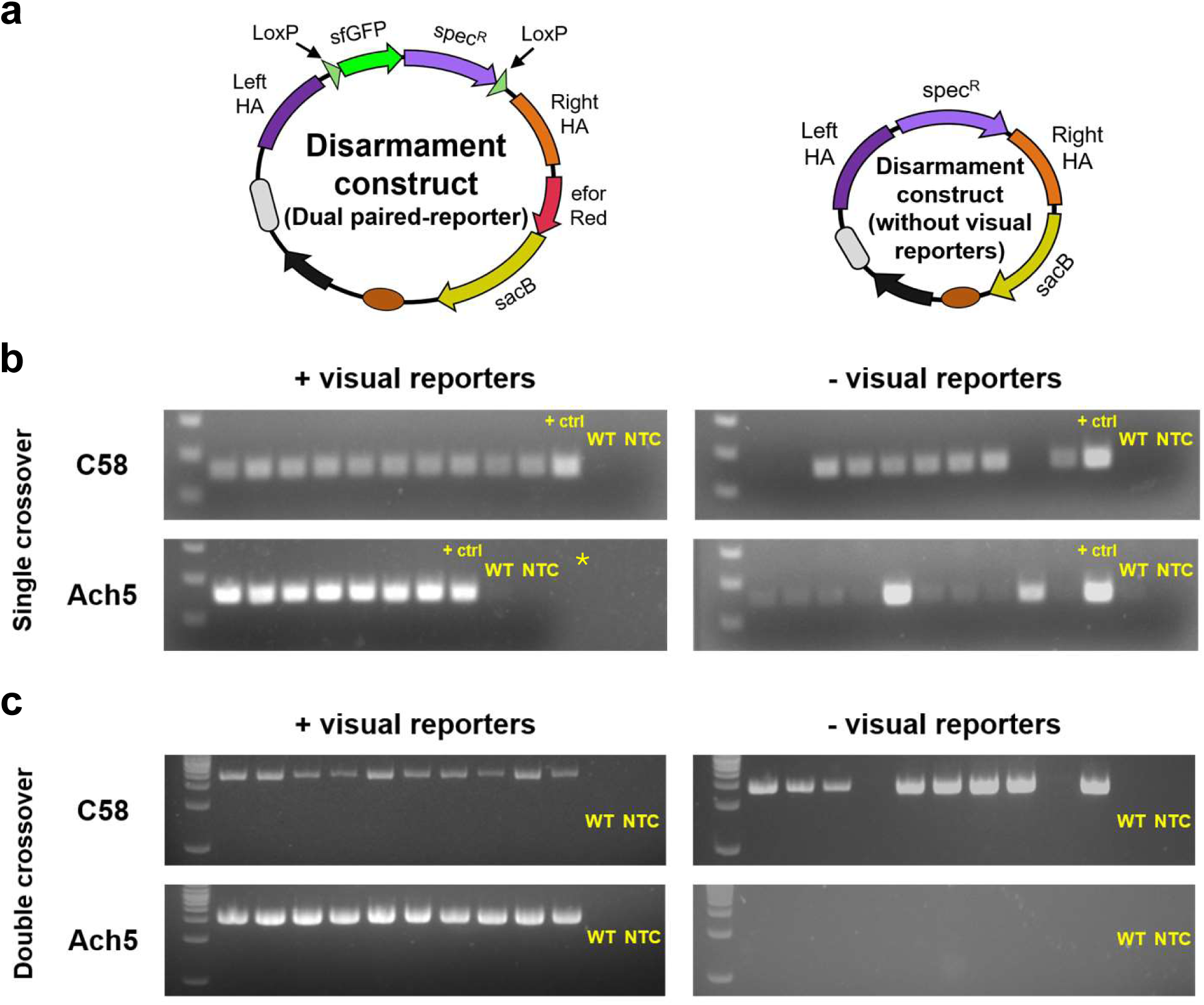
Evaluating visual reporter-assisted colony selection. **a)** Diagrams showing design of the parallel constructs assembled with or without visual reporter cassettes used for disarmament of C58 and Ach5. **b)** PCR series for single crossover recombinants performed on 10 colonies re-streaked from those formed following introduction of the above constructs. Colony selection was either guided by visual reporter expression or random. Positive colonies produced a 256 bp band from primers annealing to the spectinomycin resistance gene. Construct naked DNA (+ ctrl) and a wild-type colony (WT) were used as controls for each run. **c)** PCR series for double crossover recombinants performed on 10 colonies formed following re-plating on counterselective media of PCR-positive clones from above series. Colony selection was again either guided by visual reporter expression or random. Flanking primers specific to each strain were used, resulting in 4.8 kbp or 3.8 kbp bands for the C58 series with and without visual reporters, respectively, and 4.3 kbp or 3.3 kbp bands for the Ach5 series. *Only 7 colonies expression both visual reporters were seen following introduction of the construct containing them into Ach5. Thus, only 7 samples were screened in that PCR series for single crossover.

After colonies formed, up to 10 from each plate were used to evaluate successful recombination by colony PCR with primers probing for the spectinomycin antibiotic resistance marker (Figure 7B). Colonies transformed with the construct containing the visual reporters were selected by screening for expression of both fluorescent signals, while colonies transformed with the control construct were randomly selected. PCR positive single-crossover colonies for each construct were then pooled and replated to sucrose-containing media to select for secondary recombinants. After colonies appeared, 10 from each original transformant population were again selected, either based on them being positive for green fluorescence signal and negative for red fluorescence, in the cases of the construct containing reporters, or they were selected randomly in the case of the control construct. Colony PCR was performed using flanking primer pairs specific to each strain for assessment of double crossover/disarmament status (Figure 7C). Images of each set of original plates from which colonies were selected for this experiment and displayed in Figure S6.

In the absence of visual reporters to aid in colony selection, identification of primary recombinant colonies was still high in C58, with 7 out of 10 colonies being positive for the replacement sequence marker. However, for Ach5, only 2 out of 10 colonies were positive. In the presence of visual reporters, all selected colonies for both strains were positive by PCR, though for Ach5 only 7 total colonies expressing both fluorescent signals were formed following construct introduction. Among the colonies recovered after applying counterselection to the PCR-confirmed primary recombinant clones, in C58, 8 out of 10 colonies in the treatment without visual reporters had undergone double crossover, while for Ach5, no double crossover colonies were identified without reporters. Meanwhile, in both strain background sets with the visual reporters, the fluorescent signature assisted in correctly identifying double crossover events in all 10 colonies screened.

This experiment demonstrates a phenomenon we observed through the course of disarming the novel wild-type strain set described above—that while selection and counterselection can provide stringent filtering for colonies with the intended recombination state in some strains and contexts, (as exemplified by disarmament of C58) with our selected HA sequences, in other situations, recombinants can be relatively infrequent among the colonies formed on plates following each step in applying chemical agents. Though the frequency of correct recombinant colonies can potentially be improved by optimizing the concentration of the antibiotic and counterselective agents applied to a particular strain, visual reporter-assisted selection can mitigate the need for this, allowing quick recovery of rare recombinants without extensive and laborious PCR-based screening.

### The effect of homology arm length on recombination

Most literature sources describing modification of agrobacterial strains using HR utilize homology arms with lengths falling roughly in a window between 800 and 1,200 base pairs (Aly *et al*., 2008; Budnick *et al*., 2020; Collier *et al*., 2018; Zhong *et al*., 2025). We examined the impact of using homology arms with reduced lengths on the efficiency of recombination. In addition to the original disarmament construct we had generated for targeting the C58 strain, which contains left and right homology arms each near 1,200 bp in length, two additional constructs were assembled for which shorter homology arms about 600 or 300 bp in length (respectively) amplified from the same regions of pTiC58 were used. These three constructs were each introduced into C58 cells by electroporation in parallel and serially plated on media amended with spectinomycin. The rates of marker-positive colony formation for each construct are represented in the images in Figure S7.

This experiment showed that there is a substantial reduction in recombination frequency using shorter homology arm sequences, with very few single-crossover colonies for pTiC58 disarmament being recovered when homology arms were as small as 300 base pairs. Thus, we recommend using longer homology arms for HR construct designs whenever possible to ensure maximal targeting efficiency.

### Achieving HR in a recA-strain background

The native gene *recA*, which is widespread across bacteria, encodes a key component in what is thought to be the primary complex that facilitates homologous recombination in agrobacteria (Farrand *et al*., 1989). Modified strains in which the *recA* locus is disabled have been developed to eliminate unwanted recombination giving rise to deletions in introduced binary vector constructs (Aliu *et al*., 2022; Pennetti *et al*., 2025; Rodrigues *et al*., 2021). Further gene modification in such *recA*-strain backgrounds using HR-based methods are thus extremely difficult or impossible to achieve (Aliu *et al*., 2022).

The commonly used plant transformation background AGL1 (Lazo *et al*., 1991) is one such mutant strain. However, construct instability apparently resulting from HR has still been observed in AGL1 (Wang *et al*., 2026). One possibility for how this might occur, is via a *recA*-independent pathway involving the gene *recF*, *addA*, and *aadB*, which has been documented to function in rhizobia (Zuniga-Castillo *et al*., 2004). We evaluated a strategy for accomplishing HR-enabled gene replacement in a *recA* mutant background by targeting the *recF* for knockout in AGL1. A cloned cassette for expressing a *recA* coding sequence derived from the C58 circular chromosome (under promoter P_Atu0824_ and the rrnB T1 terminator) was incorporated just adjacent to the *sacB* counterselection marker (in the eviction sequence, donated by a modified module 4 plasmid variant) into an assembled *recF* deletion construct. A construct lacking the *recA* cassette, but which was otherwise equivalent was also assembled for deployment as a control.

By positioning the *recA* cassette in the eviction sequence, transient expression of the gene product may be enough to compensate for lack of the native sequence and facilitate single crossover resulting in integration of the construct into the chromosome. Then, being now expressed from the chromosome, could facilitate double-crossover resulting in its removal from the genome along with the counterselection marker. The *recA*-containing and control constructs were introduced into AGL1 cells in parallel by electroporation. Apparent single-crossover colonies comparable to the levels seen for C58 disarmament appeared for the culture transformed with the *recA*-containing construct, while as expected, no colonies appeared for cells transformed with the control construct (Figure S8). Marker-excised *recF*-derivatives of AGL1 were obtained from these following the established pipeline. This strategy should enable additional modifications to be made using HR in other strain backgrounds for which the *recA* gene has already been disabled. Module 4 variant plasmids pairing a *recA* cassette with the alternative counterselection markers *pheS*\* and *codA*^D314A^ are also available in the AGROtools set.

### Utilizing AGROtools for efficient gene alteration in Pseudomonas

To test whether the AGROtools kit could be applied to genetic modification of bacterial strains outside the ARC, we used the plasmid components to assemble an HR-based construct for deleting a gene in *Pseudomonas syringae* pv. *tomato* (locus tag: PSPTO_5060). Primary and secondary recombinants were obtained, identifiable by fluorescent signatures at each step. Successful deletion of the target gene was also confirmed by colony PCR (Figure S9). This experiment suggests that the utility of the AGROtools suite is not limited to the Rhizobiaceae and that these components may potentially serve as a valuable resource for gene functional studies or strain improvement in a wider variety of microbes than those for which they were designed.

### Conclusion

The diverse set of modular plasmid components generated as part of the AGROtools kit, together with the dual paired-reporter HR construct design scheme, enables rapid construct assembly and recovery of recombinant clones. For the simplest strain disarmament applications, we were able to progress from project conception and primer design to validation of disarmed clones in less than two weeks. Constructs designed using this toolkit offer substantial advantages over existing tools for this purpose such as the widely used pK18mobsacB vector (Schäfer *et al*., 1994), which requires either two rounds of restriction cloning to incorporate homology arm segments, or a bespoke assembly strategy tailored to the specific application. In contrast, AGROtools enables highly customizable modular construct assembly in a single cloning step, with the homology arm sequences themselves being the only user-provided input required for most applications. The visual reporters employed in our design further accelerate and simplify colony selection at each stage, from construct assembly through successive recombination events and (optional) subsequent marker removal for recycling. A complete list of AGROtools component plasmids, descriptions of their uses, and sequences is provided in Dataset S1, while Figure S10 presents a visual catalogue of AGROtools plasmids emphasizing combinations of molecular parts used in them.

The disarmed strains produced in this work vary in the chromosomal lineages and virulence plasmid types they are derived from. This collection thus offers an expansion of the genetic diversity of agrobacteria deployable as plant transformation tools beyond that of the relatively narrow conventional transformation strain toolbox. Future experimentation will be focused on the question of whether some of these strains can assist in overcoming barriers to T-DNA delivery in plant species and genotypes for which transformation is prohibitively inefficient or not currently achievable.

More broadly, the AGROtools kit provides a flexible platform for continued genetic improvement of agrobacterial resources for plant transformation. Its modular design offers multiple routes for generating strains with known potentially useful phenotypes, including auxotrophic and recombination-deficient mutants, while also facilitating systematic evaluation of other promising genetic targets that have not yet been widely explored. In addition to its applications in strain improvement, the toolkit could serve as a resource supporting basic genetic research in agrobacteria (and potentially in bacteria beyond this group), simplifying and accelerating the technical processes required for conducting experiments that provide insight into the biology of these important organisms.

## Materials & Methods

### Strains, media, and growth conditions

Chemically competent TOP10 *E. coli* cells (Thermo Fisher; Waltham, MA) were utilized in cloning of all AGROtools component plasmids and assembly of final constructs following methods described in Inoue *et al*. (1990), with the exception of the entry plasmid variants for modules 1-4 containing a lacZα-ccdB fragment. In these cases, DB3.1 ccdB survival cells (Thermo Fisher) were used for electroporation following Tu *et al*. (2016). *E. coli* cultures were maintained on LB solid media or liquid broth following Miller (1972) amended with the appropriate antibiotics. Antibiotic selection was applied in *E. coli* at the following concentrations: carbenicillin at 100 µg/mL, kanamycin at 50 µg/mL, spectinomycin at 100 µg/mL, gentamicin at 10 µg/mL, apramycin at 50 µg/mL, and chloramphenicol at 25 µg/mL.

A large set of wild-type agrobacterial strains originating from collections initiated by Dr. Larry Moore and isolated from samples sent to the Plant Clinic (both Oregon State University) was graciously provided to us by Dr. Jeff Chang. Wild-type strain 3a, derived from a gall found growing on hop (*Humulus lupulus*), was isolated and provided to us by Dr. Michele Wiseman.

All strains were grown and maintained on MGY media using the following recipe obtained from the Oregon State University Plant Clinic: D-Mannitol (10 g/L), L-glutamic acid, monosodium salt (2 g/L), KH_2_PO_4_ (0.5 g/L), NaCl (0.2 g/L), MgSO_4_·7H_2_O (0.2 g/L), and yeast extract (1 g/L), adjusted to pH=7.0, with 15 g/L of BD Bacto™ agar added for solid formulations. Antibiotics and other amendments were added to the base media as appropriate to select for recombinant colonies (see Dataset S7 for details). Cultures plated onto solid media were maintained at 28°C. Cultures initiated in liquid broth were grown at 28°C with shaking at 200 rpm.

### Cloning and plasmid assembly

AGROtools plasmid components were generated using sequences which were PCR-amplified as segments from plasmid templates obtained from Addgene, plasmids sent to us directly by colleagues (listed in Table S2), or from genomic DNA isolated from agrobacteria, as well as synthetic gBlock fragments (Dataset S2) and annealed oligonucleotides (IDT; Coralville, IA). Cloning of the components was accomplished using golden gate, restriction/ligation, and Gibson assembly approaches. Assembly of the disarmament and gene deletion constructs used in this study from AGROtools components was performed using golden gate assembly with the *PaqCI* restriction enzyme and T4 DNA ligase (New England Biolabs; Ipswich, MA). More detailed cloning procedures used in development of molecular parts used in AGROtools and modification construct assembly are given in Methods S1, S2, S3, and S4.

### Construct introduction into agrobacterial cells

Constructs were introduced into agrobacterial strain backgrounds via either electroporation at room temperature or by triparental mating, the procedures for which are described in detail in Methods S5 and S6, respectively.

### Fluorescent reporter detection and image acquisition

Following introduction of constructs expressing sfGFP and/or eforRed fluorescent visual reporters, *E. coli* or agrobacterial colonies appearing on plates were quickly screened using the Royal Blue (for green fluorescence) and Green (for red fluorescence) versions of the Xite Portable Fluorescence Flashlight System (NIGHTSEA; Lexington, MA). Images of colonies on plates were acquired using a Nikon SMZ25 fluorescence stereomicroscope equipped with GFP and TRITC filter sets.

### Quantitative analysis of reporter expression

A Spark® multimode microplate reader (Tecan; Männedorf, Switzerland) was used to collect fluorescence intensity data both for the experiment evaluating visual reporter cassettes across agrobacterial strains and for the comparison of fluorescent transgene transient expression in leaves infiltrated with disarmed strain derivatives. Samples were deposited into a 96-well plate (Greiner Bio One GmbH; Kremsmünster, Austria) in triplicate wells for bacterial suspension cultures and in sets of 12 replicates for leaf samples. Detailed procedures describing the staging and analysis of the visual reporter/promoter characterization assay and the transient transgene expression assay are given in Method S7 and Method S10, respectively.

### Generation of disarmed strains

A detailed process for handling of target strains following introduction of a modification construct is described in Method S8.

### Bacterial genome sequencing

Total genomic DNA was extracted from each of the final disarmed or modified and marker-excised strain derivatives using the Wizard® DNA Purification Kit (Promega; Madison, WI) on 2-mL saturated cultures of each, following the protocol for gram-negative bacteria. Concentration and purity of each sample was assessed using a Qubit 4 Fluorometer and a Nanodrop 2000 (both from Thermo Fisher Scientific; Waltham, MA), respectively. Samples were diluted to 50 ng/µL concentrations and 20 µL of each was sent to Plasmidsaurus Inc. for bacterial whole genome sequencing. A custom assembly pipeline using reads generated by Plasmidsaurus, followed by downstream analysis of modified strain genomes for target specificity is discussed in Method S9.

### Agroinfiltration assay

Disarmed and marker-excised novel strains, along with the conventional transformation strain AGL1 were transformed with the same binary plasmid (pT200-GFP-HygR) for the purpose of assessing their transient T-DNA delivery. Each strain derivative was grown from an overnight primary culture in 10 mL of MGY broth amended with kanamycin at 50 mg/L. Cells were then prepared for agroinfiltration according to methods described in Leuzinger *et al*. (2013). Each strain was flooded into young leaves of two separate 4 to 6 week-old plants. The AGL1 derivative was injected into a leaf on each plant included in the experiment as a positive control, while leaves flooded with only infiltration buffer served as a negative control. For the qualitative assay, at 3 days post-inoculation whole leaves were detached, cut with a razor blade to lay the flooded sectors as flat as possible, and imaged under the fluorescence microscope in the bright field and green fluorescence channels. Method S10 details the procedures used for collection and quantitative analysis of fluorescence intensity data in an additional agroinfiltration experiment.

### Additional methods

Detailed procedures used for evaluating counterselection markers, testing the utility of dual paired reporters in disarmament, assessing disarmament using homology arms of varying lengths, knockout of *recF* in the AGL1 background, and gene deletion in *P. syringae* are given in Methods S11, S12, S13, S14, and S15.

## Acknowledgements

We thank several colleagues for helpful discussions and ideas on design and use of strain engineering tools and toolkit parts, particularly Ray Collier (Wisconsin Crop Innovation Center), Bill Gordon-Kamm (Corteva Agriscience), Keunsub Lee (Department of Agronomy, Iowa State University), Vincent Pennetti (Department of Crop & Soil Sciences, University of Georgia), Savio Rodrigues (Department of Biosystems, KU Leuven), along with Jim Thomson and Roger Thilmony (USDA-ARS). Roger also suggested the use of native *Agrobacterium* promoter (P_Atu0824_) and gifted us with plasmid pCGT4464A. We thank Jeff Chang, Alex Weisberg (Department of Botany and Plant Pathology, Oregon State University) and lab members for their grace and patience in helping us procure large parts of their agrobacterial strain collections which we utilized in this project. Alex was particularly helpful with technical suggestions on genome extraction/sequencing analysis. We are also grateful to Michele Wiseman for her efforts in isolating strain ‘3a’ from gall tissue (Department of Botany and Plant Pathology, Oregon State University). Finally, we express our gratitude to the following researcher for sharing constructs with us via Addgene, which we utilized in construction of various toolkit components: Kan Wang, Daniel Voytas, Bing Yang, Wayne Parrott, Jun Fu, Sylvestre Marillonnet, Anthony Forster, Philip Poole, and Christopher Reisch.

## Author contributions

C.W. conceived the project, oversaw experimental work, and prepared the manuscript. B.M.S., E.B., J.A., T.W., J.W., P.M.D.L., and C.W. developed plasmid toolkit components. Z.H., C.W., C.R., T.W., J.W., and P.M.D.L. disarmed and modified strains. Z.H. performed whole genome sequencing analysis and *N. benthamiana* agroinfiltration assays. B.M.S. constructed and tested activity of reporter expression cassettes. E.B. performed homology arm length assessment and *recA* complementation studies. G.N., S.K., and J.C.A. performed the *Pseudomonas* gene deletion work. G.S.G. gave critical advice and assisted with manuscript preparation. S.H.S. provided project direction.

## Data and Materials Availability

AGROtools plasmid components are being deposited in Addgene for distribution. Whole genome sequence assemblies will be made available via a Github page linked to our lab’s website. Disarmed strains described in this manuscript will be provided to academic researchers upon request following issuance of applicable USDA APHIS movement permit and subject to the terms of a signed MTA. A fee will be charged to recover costs associated with strain distribution.

