## Supplemental Materials for "AGROtools: A comprehensive modular cloning toolset for *Agrobacterium* strain engineering"

**Supplementary material contents**

**Figure S1:** Detailed design of generalized module 2, module 4, and backbone plasmid components.

**Figure S2:** Construct assembly and usage for alternative (*recA*-independent) modification methods.

**Figure S3:** PCR validation of disarmament, genomic modification, and marker excisions by HR.

**Figure S4:** Agroinfiltration of disarmed strains carrying a binary vector into *Nicotiana benthamiana* leaves.

**Figure S5:** Fluorescence of agroinfiltrated *Nicotiana benthamiana* leaves with multiple disarmed agrobacteria strains.

**Figure S6:** Plate images for strains disarmed using constructs with and without visual markers.

**Figure S7:** Homology arm length comparison for disarmament in C58.

**Figure S8:** *RecA* complementation in a construct restores HR in a *recA*- strain background.

**Figure S9:** AGROtools can be used to facilitate gene knockout in *Pseudomonas*.

**Figure S10:** Graphic catalogue of AGROtools plasmid components.

**Table S1:** Summary of AGROtools components layout, molecular parts, and naming conventions for utilizing *recA*-independent modification systems.

**Table S2:** Original plasmid sources for parts used in constructing AGROtools.

**Method S1:** Primer design, PCR fragment amplification and handling, origin of molecular fragments, and general cloning methodology.

**Method S2:** Cloning and production of agrobacterial strain variants for evaluating visual reporters.

**Method S3:** Assembly of all AGROtools plasmid components.

**Method S4:** Assembly of disarmament and modification constructs.

**Method S5:** Construct introduction into agrobacteria cells by electroporation.

**Method S6:** Construct introduction into agrobacteria cells by triparental mating.

**Method S7:** Detailed plate reader assay.

**Method S8:** Disarmament of wild-type strains and marker excision.

**Method S9:** Genome assembly and analysis of modified strains.

**Method S10:** Quantitative assay of reporter expression from agroinfiltrated disarmed strains.

**Method S11:** Counterselection agent/marker testing assay.

**Method S12:** Assessment of dual paired reporter constructs for disarmament.

**Method S13:** Homology arm length evaluation.

**Method S14:** Knockout of *recF* in the AGL1 background.

**Method S15:** Gene deletion in *Pseudomonas*.

**Supplemental References:** (Applies to supplemental tables and methods)

**Dataset S1:** Complete list of AGROtools component plasmids.

**Dataset S2:** Synthesized gene fragments used in toolkit construction.

**Dataset S3:** Primers used for visual reporter parts construction.

**Dataset S4:** Primers used for restriction sequence purging by site-directed mutagenesis.

**Dataset S5:** Primers used in AGROtools module construction.

**Dataset S6:** Primers used in homology arm amplification for disarmament and other modification construct assembly.

**Dataset S7:** Additional information on disarmed and otherwise modified strains.

**Dataset S8:** Raw output from fluorescence plate reader assay.

**Dataset S9:** Flanking primer sequences used to validate secondary recombination and marker excision.

**Dataset S10:** Sequence of binary plasmid pT200-GFP-HygR.

**Dataset S11:** Primers used in accessory plasmid construction and for validating primary recombination.

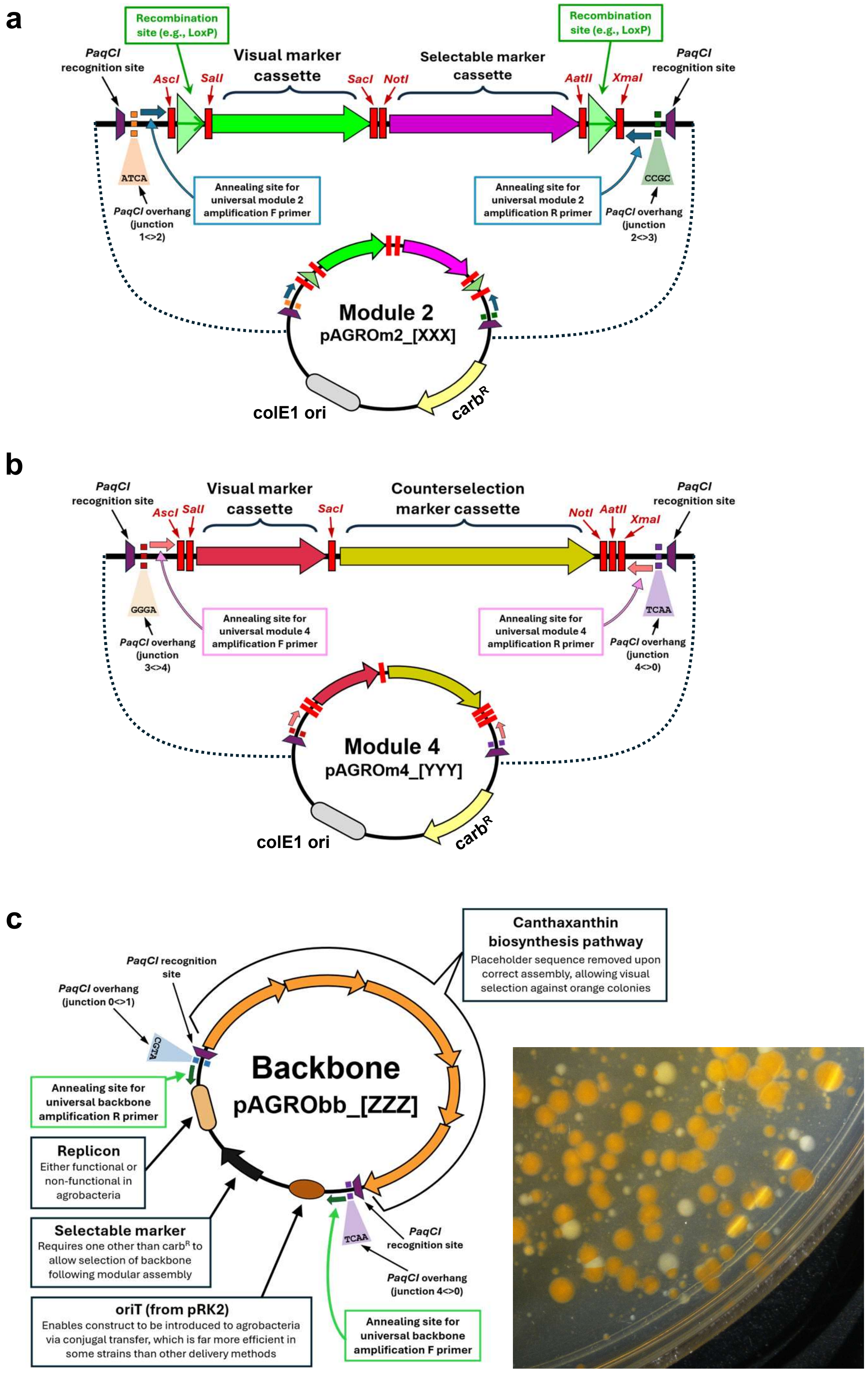


**Figure S1: Detailed design of generalized module 2 (a), module 4 (b), and backbone plasmid (c) components**

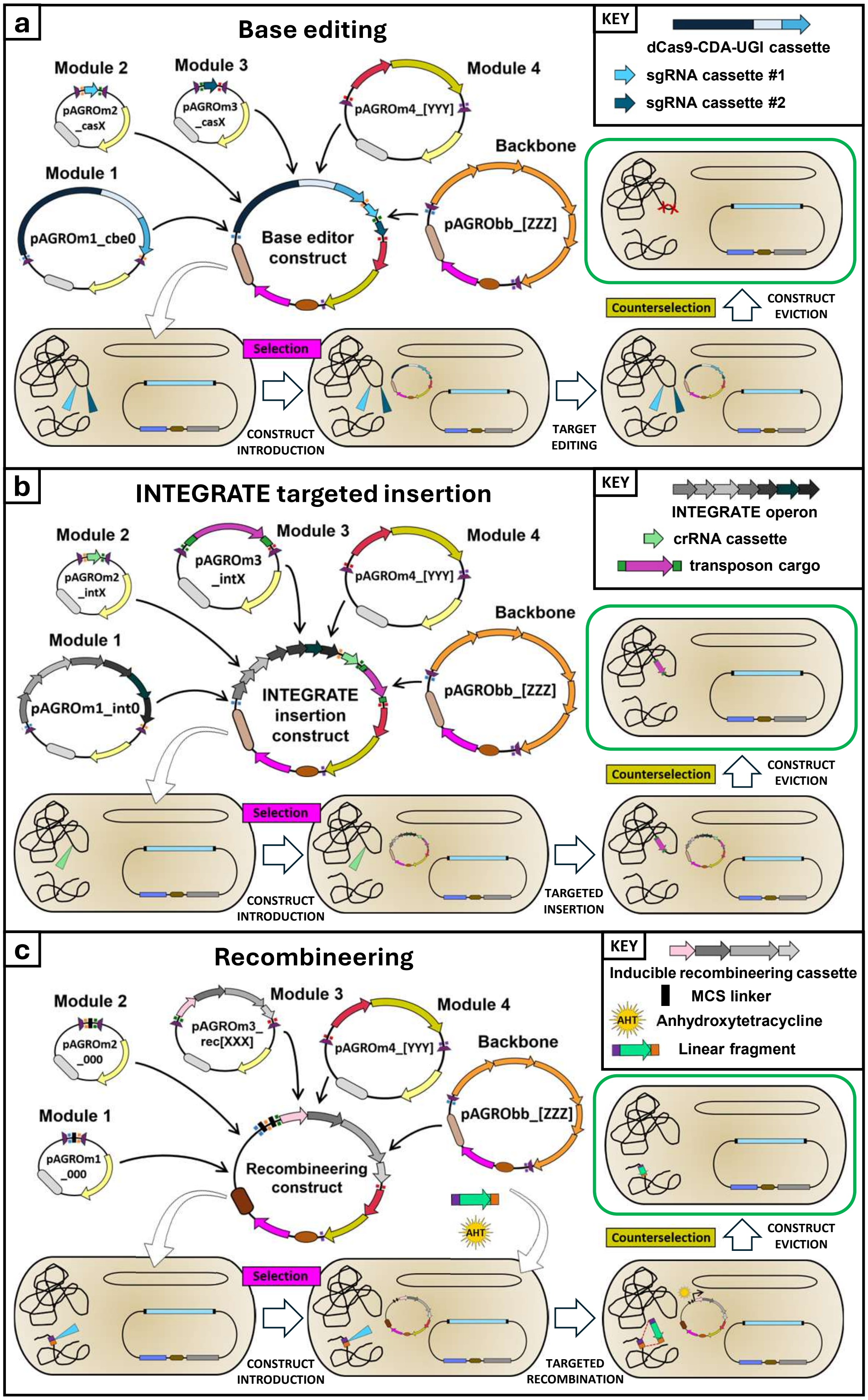


**Figure S2: Construct assembly and usage for alternative (*recA*-independent) modification methods. a)** Using cytidine base editing to mutate chromosomal gene targets (light and dark blue triangles). The user clones custom spacer sequences into module 2 and/or 3 entry plasmids prior to assembly. **b)** Using the INTEGRATE system to make a sequence insertion at a chromosomal target site (light green triangle). The user clones custom spacer sequence(s) and custom insertion cargo segment into module 2 and module 3 entry plasmids, respectively, prior to assembly. **c)** Using a recombineering system for targeted introduction of a linear fragment at a chromosomal site (light blue triangle). The user supplies an inducer compound (anhydrotetracycline) and a linear fragment following construct introduction into the cell. Recombination can occur based on two short (~80-bp) homology arm sequences at each end of the fragment (purple and orange rectangles).

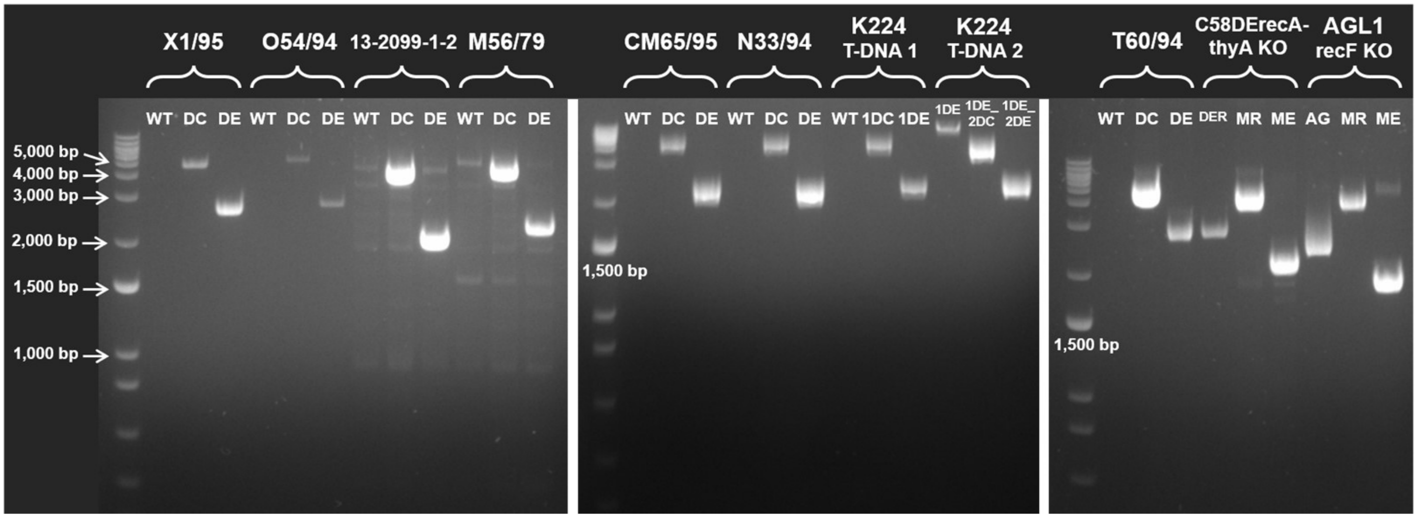


**Figure S3: PCR validation of disarmament, genomic modification, and marker excisions by HR.** Colony PCR samples showing double crossover replacement and marker excision for each of the disarmed and otherwise modified agrobacterial strains backgrounds not shown in Figure 6. For disarmed strains, WT = wild-type, DC = double crossover disarmed, DE = disarmed-excised. 1DC and 1DE indicate double crossover and disarmed-excised isolates (respectively) for T-DNA 1 in K224, while 1DE_2DC and 1DE_2DE indicate double crossover and disarmed-excised isolates (respectively) for T-DNA 2 in the T-DNA 1 deleted background. For strains in which chromosomal gene modifications were carried out, DER indicates the disarmed-excised and *recA-* C58 strain background in which the *thyA* knockout was performed. AG = AGL1 base strain, MR = marker retained (double crossover), and ME = marker excised. “flanking” primers used in validation PCR for each modified strain are given in Dataset S8.

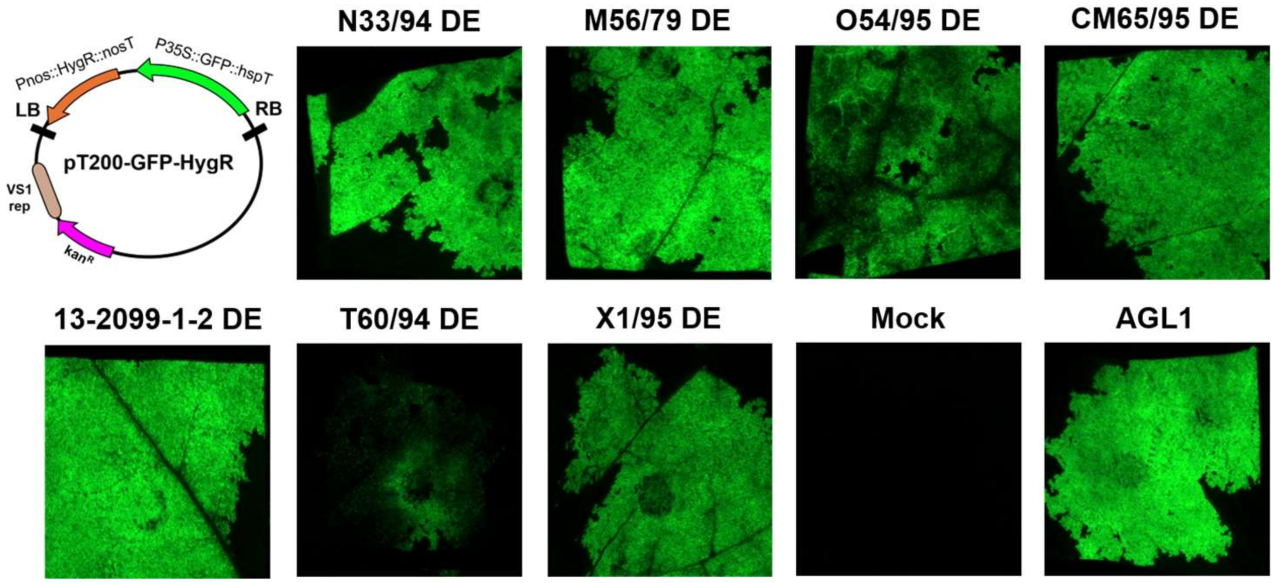


**Figure S4: Agroinfiltration of disarmed strains carrying a binary vector into *Nicotiana benthamiana* leaves.** The construct diagram in the upper left depicts the binary plasmid introduced into each of the disarmed strain backgrounds for this experiment. Images show GFP expression representative of two leaves on different plants, infiltrated with each strain derivative after 3 days. Mock treatment (infiltration buffer only) and standard plant transformation strain AGL1 served as negative and positive controls, respectively.

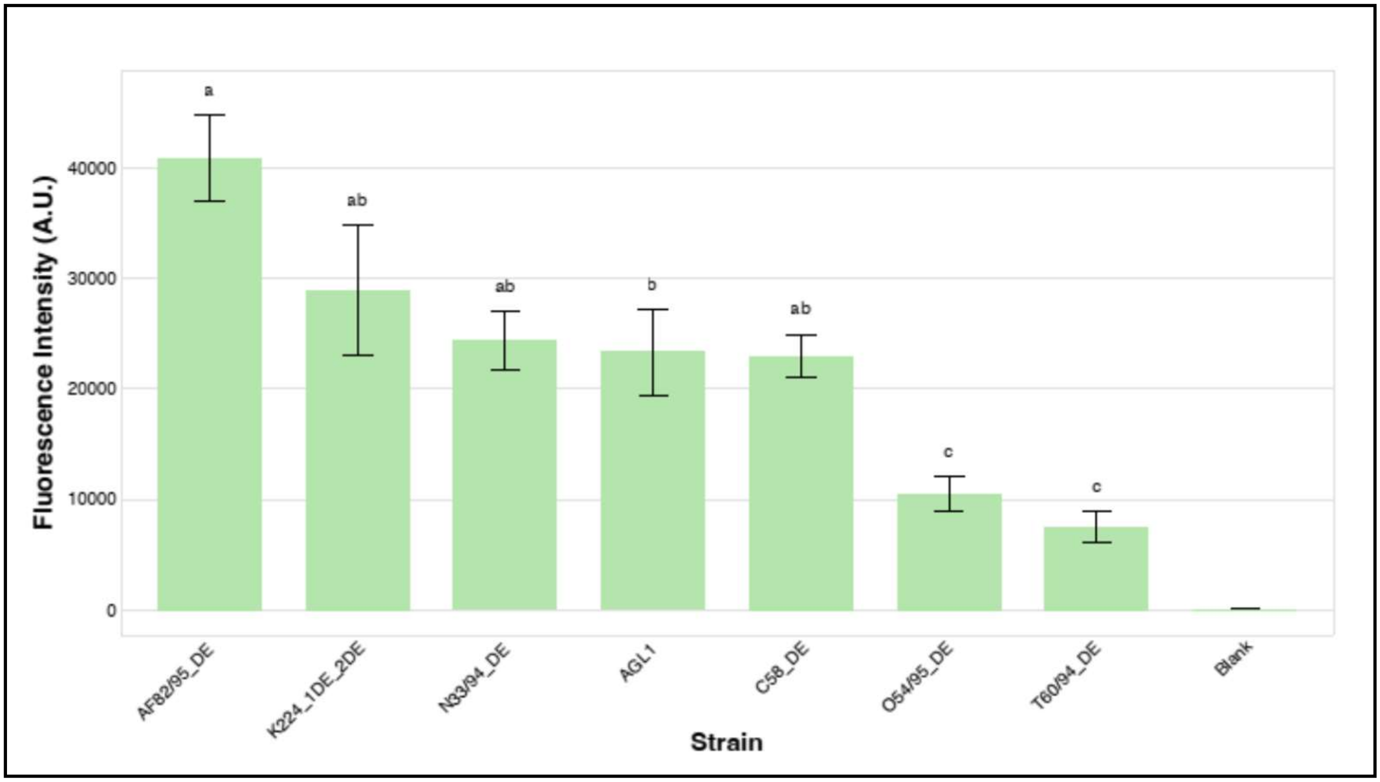


**Figure S5: Fluorescence of agroinfiltrated *Nicotiana benthamiana* leaves with multiple disarmed agrobacteria strains.** Bars show mean fluorescence ± standard error, ordered by decreasing mean. Blank controls included for reference. Compact letter display indicates Dunn's post hoc groupings with Benjamini-Hochberg correction, statistical analysis and experimental procedure described in Methods S10. Strains sharing a letter are not significantly different (adjusted *p*-value > 0.05).

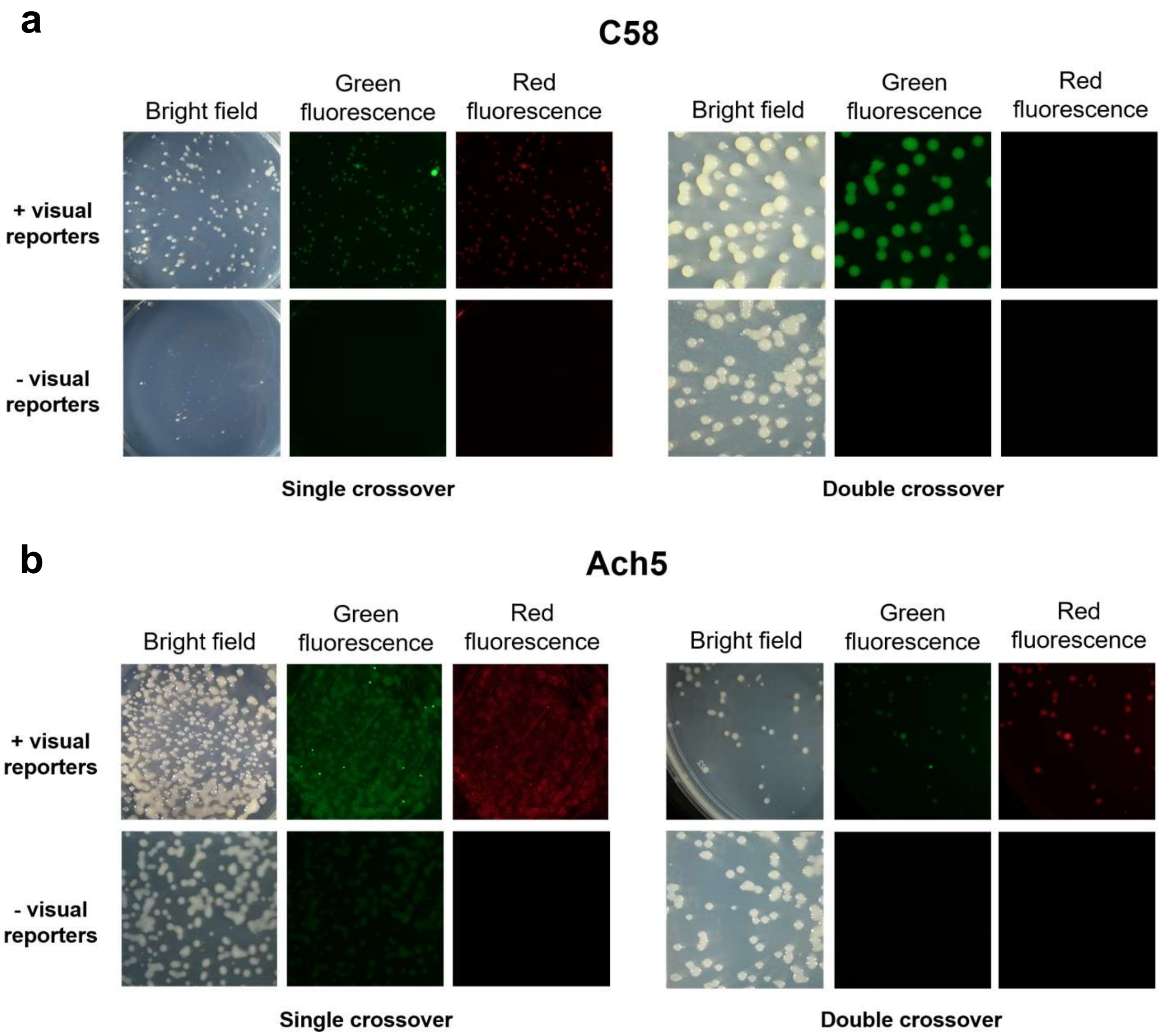


**Figure S6: Plate images for strains disarmed using constructs with and without visual markers.** Plates corresponding to the colony PCR gene images shown in Figure 7 for C58 **(a)** and Ach5 **(b)** disarm recombinant series. For the single crossover series in the left panels, images show colony growth on plates 3 days following electroporation, while the right panels show images of colonies formed following plating of PCR-confirmed single crossover clones onto counterselection media. Images for Ach5 single crossover plates were acquired using 3X the exposure of the other images owing to the high background fluorescence and relatively weak signal of recombinant colonies in the "+ visual reporters" treatment, thus a background signal can be seen in the colonies for the "- visual reporters" treatment.

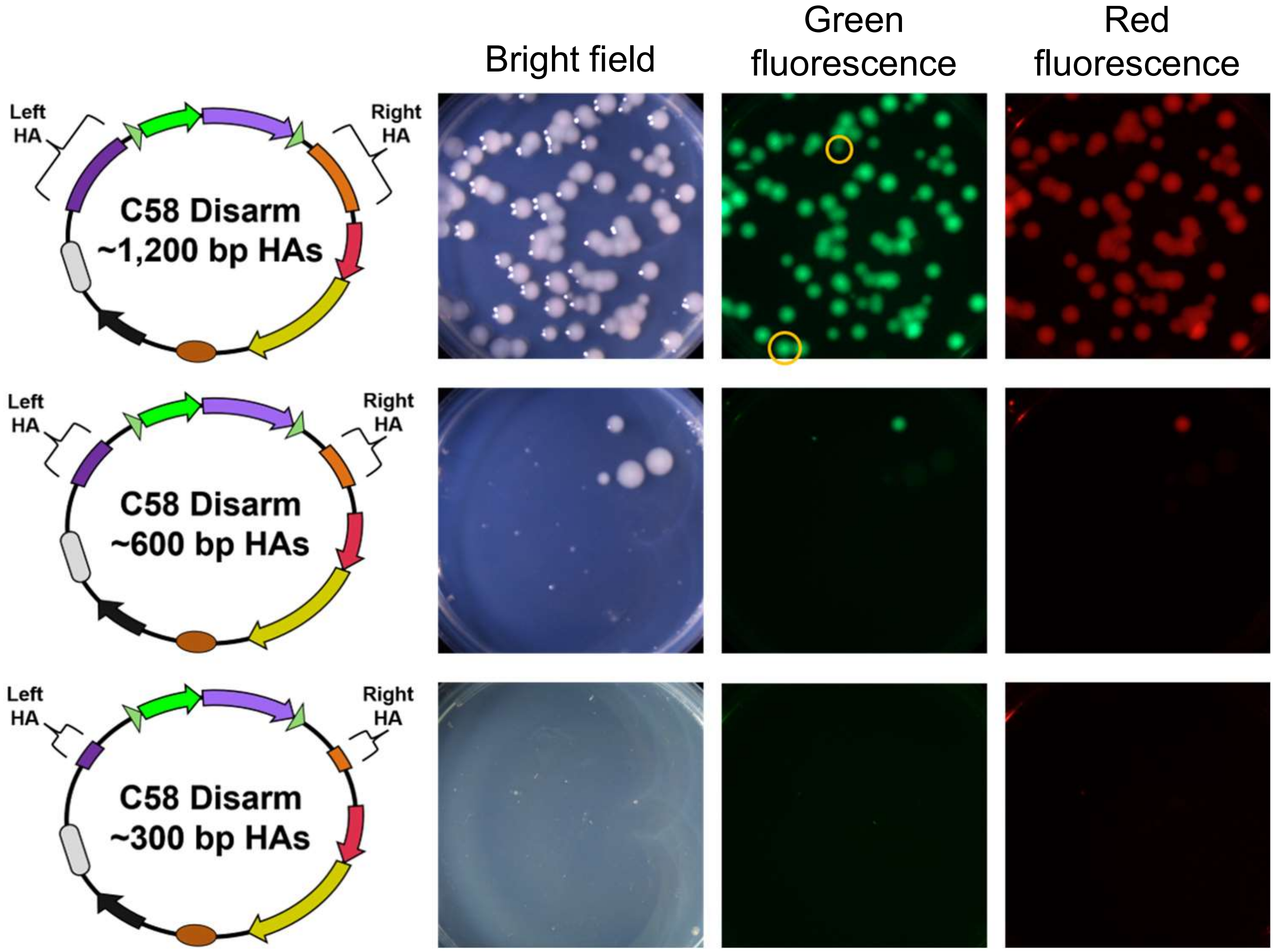


**Figure S7: Homology arm length comparison for disarmament in C58.** Diagrams on the left show the design of constructs differing only in the length of the homology arm segments, which were introduced into the C58 background. Plate images show colony formation after introduction of each construct and placement on selection after 3 days. Yellow circles on the green fluorescence image in the top row show apparent double-crossover events in the first culture plating following electroporation without the application of counterselection, as evidence by a lack of red fluorescent signal for these colonies in the top right image.

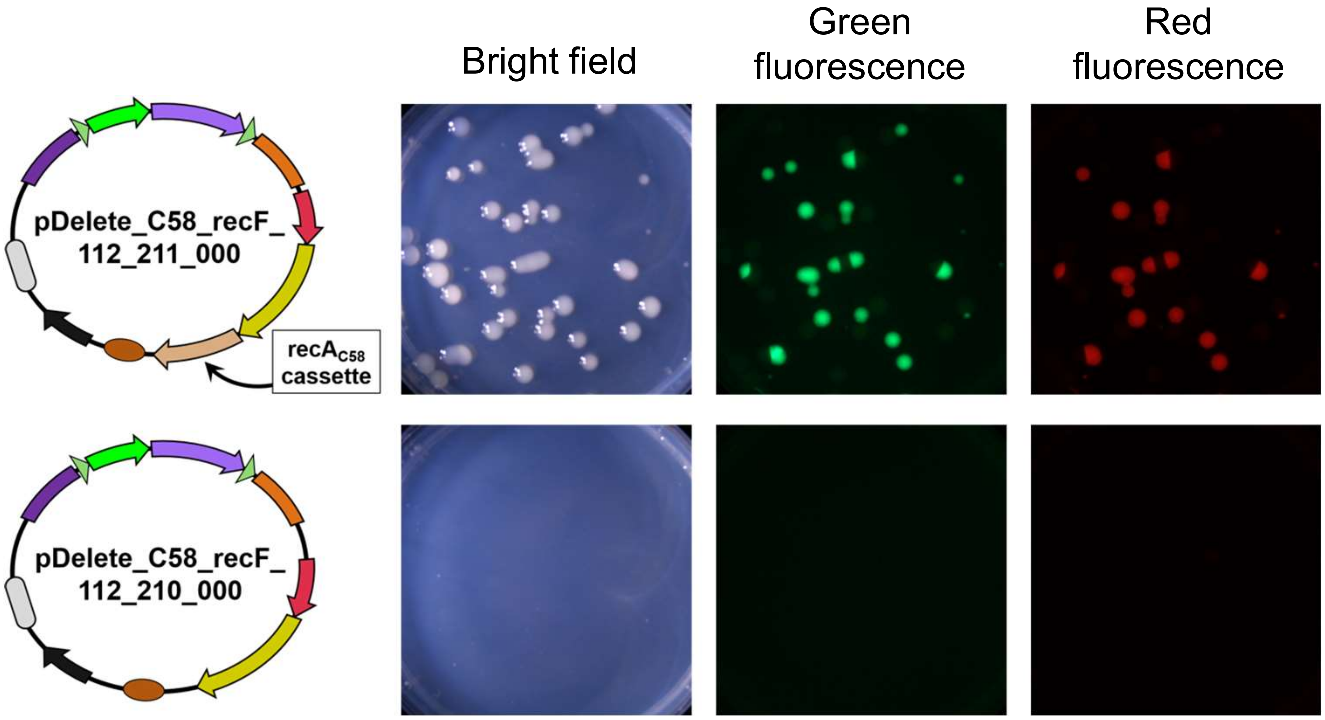


**Figure S8: *RecA* complementation in a construct restores HR in a *recA-* strain background.** Diagrams on the left show the design of HR constructs targeting the *recF* gene for deletion, differing only by whether they carry in the eviction sequence, an expression cassette for the *recA* gene cloned from C58. These constructs were introduced in parallel into the AGL1 background (a *recA*- strain) by electroporation. Plate images show colony formation after introduction of each construct and placement on selection after 3 days. The *recA*-containing construct enabled successful generation of *recF* knockouts, while no recombinant colonies were recovered when attempted modification without supplemental *recA* expression.

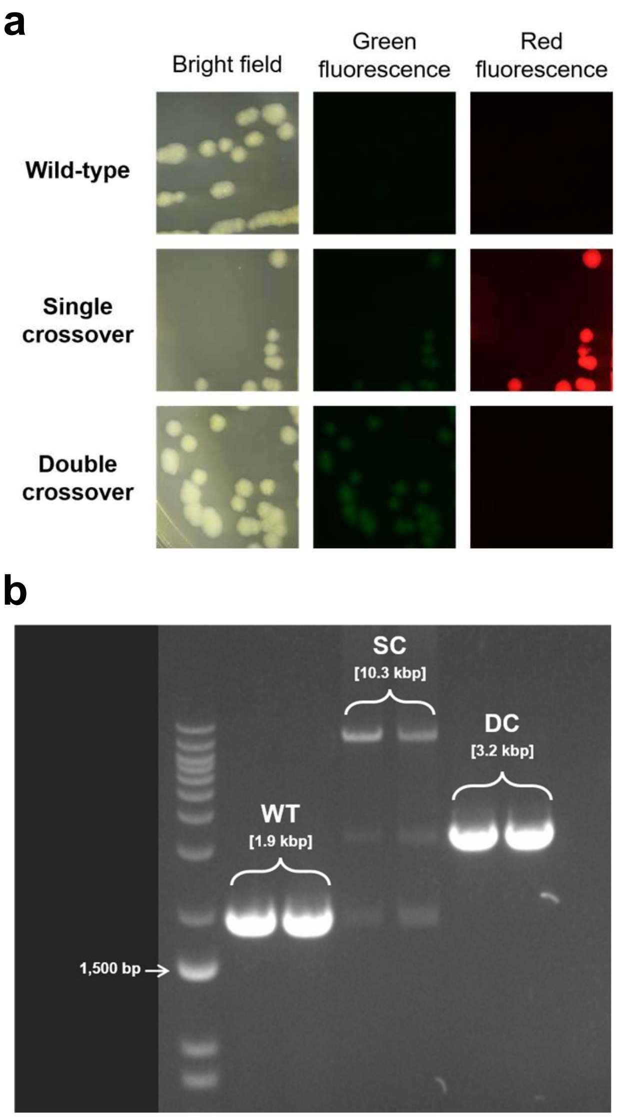


**Figure S9: AGROtools can be used to facilitate gene knockout in *Pseudomonas* a)** Images of streaked *Pseudomonas* clones representing each recombination state. **b)** A gel showing colony PCR samples using flanking primers to validate each recombination state. WT = wild-type *P. syringae* pv. *tomato* DC3000, SC = single-crossover, DC = double-crossover

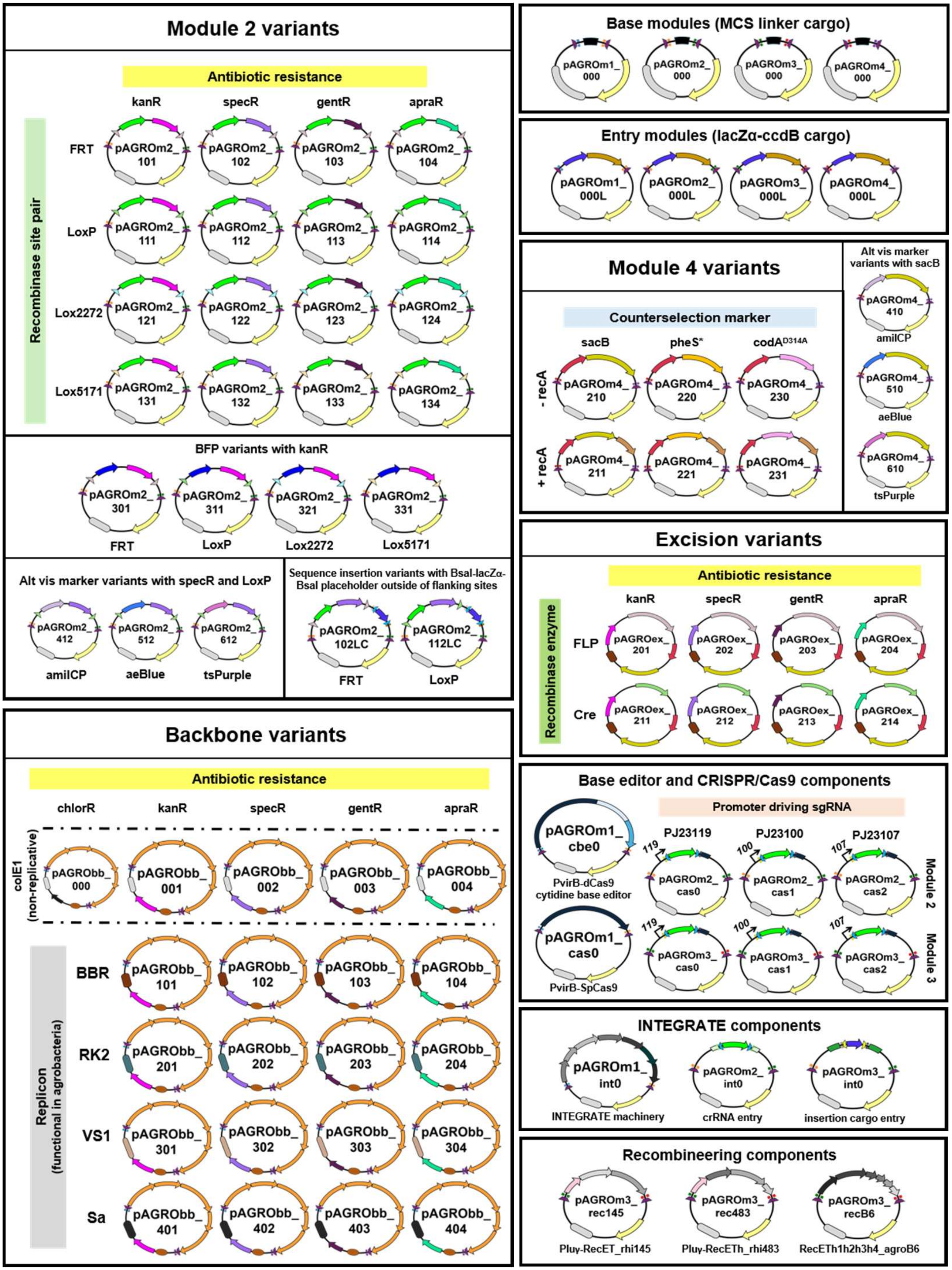


**Figure S10: Graphic catalogue of AGROtools plasmid components**

| Component class | Plasmid name | Genome engineering method | Description of molecular part/cargo | Molecular part expression elements/variants (name suffix code) |
|---|---|---|---|---|
| Module 1 | pAGROm1_cbe0 | CRISPR/dCas9 cytidine base editor | Chimeric protein consisting of the catalytically inactive dCas9 fused to a cytidine deaminase, a uracil glycosylase inhibitor, and an LVA degradation tag enabling targeted C to T base transitions | Expressed under the virB promoter from pTiBo542 and the T3 terminator (cbe0) |
| | pAGROm1_cas0 | CRISPR/Cas9 (active nuclease) | Catalytically active SpCas9 enabling targeted double-stranded break generation; could be combined with a recombineering system for CRISPR-assisted recombineering approach | Expressed under the virB promoter from pTiBo542 and the T3 terminator (cas0) |
| | pAGROm1_int0 | INTEGRATE (targeted insertion system) | Gene cluster constituting the protein complex of the type I-F CRISPR-Cas and associated transposase system derived from *Vibrio cholerae* Tn6677 enabling targeted insertion of a sequence cargo | Expressed under the PJ23119 promoter without a terminator (int0) |
| Module 2 | pAGROm2_cas[X] | CRISPR/dCas9 cytidine base editor; CRISPR/Cas9 (active nuclease) | Entry plasmid for cloning a single sgRNA spacer sequence for cytidine base editing or double stranded break targeting. sfGFP expression cassette flanked by divergent BsaI sites serves as a placeholder sequence for spacer insertion upstream of an RNA scaffold sequence. | Expressed under PJ23119 (cas0), PJ23100 (cas1), or PJ23107 (cas2) promoters with sgRNA scaffold followed by polyT |
| | pAGROm2_int0 | INTEGRATE (targeted insertion system) | Entry plasmid for cloning one or more crRNA spacer sequences for INTEGRATE targeted insertion. sfGFP expression cassette flanked by divergent BsaI sites serves as a placeholder sequence (with green fluorescence as a visual cloning aid) for spacer insertion between two crRNA repeat sequences. | Expressed under the PJ23119 promoter continuous with operon in pAGROm1_int0 cargo, followed by the T7 terminator (int0) |
| Module 3 | pAGROm3_cas[X] | CRISPR/dCas9 cytidine base editor; CRISPR/Cas9 (active nuclease) | Entry plasmid for cloning a single sgRNA spacer sequence for cytidine base editing or double stranded break targeting. sfGFP expression cassette flanked by divergent BsaI sites serves as a placeholder sequence for spacer insertion upstream of an RNA scaffold sequence. | Expressed under PJ23119 (cas0), PJ23100 (cas1), or PJ23107 (cas2) promoters with sgRNA scaffold followed by polyT |
| | pAGROm3_int0 | INTEGRATE (targeted insertion system) | Entry plasmid for cloning insertion sequence cargo for INTEGRATE targeted insertion. A lacZα expression cassette flanked by divergent BbsI sites serves as a placeholder sequence (enabling blue-white cloning as an aid) for inserting custom sequences between the required Left and Right transposon ends. | Cargo insertion site flanked by Left and Right end arms from Tn6677 transposon |
| | pAGROm3_ rec[XXX] | Recombineering systems | Inducible expression cassette for a RecET-like operon (of phage origin) with or without the "γ" ssDNA binding protein from *Photorhabdus luminescens* (Pluγ) | Pluγ and/or RecET cluster from *Rhizobium* sp. LC145 (rec145), sp. Root483D2 (rec483), or *A. tumefaciens* B6 (recB6) each expressed under a Tet operator sequence and terminated with the rrnB T1 and T2 sequences. iIn each case, the operon is positioned downstream from a divergent TetR cassette terminating in the bidirectional tonB terminator |

**Table S1** Summary of AGROtools components layout, molecular parts, and naming conventions for utilizing *recA*-independent modification systems

| Plasmid name | Parts obtained from | Where obtained from | Accession number | Depositer/ Backbone manufacturer | Reference |
|---|---|---|---|---|---|
| pEA186 | specR; INTEGRATE machinery; crRNA cassette; INTEGRATE cargo sequence | Addgene | 187874 | Kan Wang/ Keunsub Lee | Aliu *et al*. (2022) |
| pKL2315 | Cre cassette | Addgene | 187876 | Kan Wang/ Keunsub Lee | Aliu *et al*. (2022) |
| pKL2299A | RK2 replicon; gentR | Addgene | 222105 | Kan Wang/ Keunsub Lee | Aliu *et al*. (2024) |
| pMOD_B2303 | colE1 ori; colE1 ori/carbR fragment | Addgene | 91068 | Daniel Voytas | Čermák *et al*. (2017) |
| pTRANS_220 | ccdB | Addgene | 91113 | Daniel Voytas | Čermák *et al*. (2017) |
| pMIN-VS1 | VS1 replicon; kanR | Addgene | 197783 | Daniel Voytas | Chamness *et al*. (2023) |
| pFlp-Km | BBR replicon; FLP cassette; sacB | Addgene | 222334 | Philip Poole | Jorrin *et al*. (2024) |
| pEN-L1-PJ23119-PplpT-sfGFP-TrrfB-BsaI-Scaf-L2 | sfGFP; sgRNA cassette | Addgene | 196250 | Bing Yang | Li *et al*. (2023) |
| pAGM4673 | canthaxanthin operon | Addgene | 48014 | Sylvestre Marillonnet | Weber *et al*. (2011) |
| eforRed chromoprotein | eforRed | Addgene | 117837 | Anthony Forster | Liljeruhm *et al*. (2018) |
| aeBlue chromoprotein | aeBlue | Addgene | 117846 | Anthony Forster | Liljeruhm *et al*. (2018) |
| pWLlR5 | Sa replicon | Addgene | 149492 | Christopher Reisch | Schuster and Reisch (2021) |
| pVP143 | CRISPR/dCas9-CDA-UGI | Addgene | 222080 | Wayne Parrott | Pennetti *et al*. (2025) |
| pBBR1-kan-Ptet-plu-gamma-ET_rhi145 | Ptet-plu-gamma-ET_rhi145 recombineering system | Addgene | 180427 | Jun Fu | Bian *et al*. (2022) |
| pBBR1-kan-Ptet-RecETh1h2h3h4_agrob6 | Ptet-RecETh1h2h3h4_agrob6 recombineering system | Addgene | 180428 | Jun Fu | Bian *et al*. (2022) |
| pBBR1-kan-Ptet-plu-gamma-ETh_rhi483 | Ptet-plu-gamma-ETh_rhi483 recombineering system | Addgene | 180429 | Jun Fu | Bian *et al*. (2022) |
| pCGT4464A | $oriT_{pRK2}$; chlorR | Gift from Roger Thilmony | N/A | Roger Thilmony/Ray Collier | Collier *et al*. (2018) |

**Table S2**. Original plasmid sources for parts used in constructing AGROtools

**Method S1**: Primer design, PCR fragment amplification and handling, origin of molecular fragments, and general cloning methodology

All primers were designed manually based on the template sequence context with the NEB Tm calculator (https://tmcalculator.neb.com) being used to polish the template-annealing portion of primer candidate sequences to bring primer pairs within a <2°C difference using the Q5 high-fidelity DNA polymerase default setting. For golden gate assemblies, the NEBridge GetSet® Tool (https://ligasefidelity.neb.com/getset/run.cgi) was used to identify compatible 4-base overhangs predicted to yield >95% ligation fidelity according to the specific type IIS restriction enzyme used. For Gibson assembly schemes, fragment overlaps were designed manually to be 20-28 bp in length with a Tm value above 48°C.

All PCR amplification in this study was performed with Q5® high-fidelity DNA polymerase (New England Biolabs; Ipswich, MA) using a modification of the standard protocol (an additional 2.5 mM of $MgCl_2$ in excess of that already contained in the provided reaction buffer and 300 mM DMSO). Otherwise, reaction parameters used conformed to the product protocol and the NEB Tm calculator.

All PCR amplicons and restriction digest fragments used in cloning were purified by gel electrophoresis on a 1% agarose gel at 110V for 25-60 minutes, followed by extraction of the appropriately-sized DNA band using a Zymoclean Gel DNA Recovery Kit (Zymo Research; Irvine, CA) following the standard protocol.

Sequences for the sfGFP, eforRed, aeBlue, sacB, kanR, specR, gentR, chlorR, ccdB, $oriT_{pRK2}$, FLP recombinase, Cre recombinase, colE1 ori, BBR ori, RK2 ori, VS1 ori, and Sa ori molecular parts were amplified by PCR using plasmid templates obtained from Addgene or sent to us directly by colleagues (all listed in Table S2). The $recA_{C58}$ coding sequence was amplified from C58 wild-type genomic DNA.

BFP, amilCP, tsPurple, apraR, pheS*, $codA^{D314A}$, and lacZα molecular parts as well as a library of promoter and terminator sequences ($P_{J23119}$, $P_{J23100}$, $P_{int}$, $P_{Atu8024}$, T3, T7, and rrnB T1) were synthesized as gBlock fragments from Integrated DNA Technologies (IDT; Coralville, IA) (see Dataset S2 for sequences). All sequences in the AGROtools kit intended to be utilized for assembling HR constructs were purged of recognition sites for the type IIS restriction enzymes *PaqCI/AarI*, *BsaI*, *BbsI*, *BsmBI/Esp3I*, and *SapI*. Thus, these sites were all excluded from the synthetic sequences by manual inspection following any codon-optimization prior to ordering.

Plasmid cloning was accomplished using either golden gate assembly, Gibson assembly, or restriction digestion/ligation.

In general for golden gate assemblies, all constituent fragments for producing a given plasmid were added in equimolar quantities (totaling no more than 1 µg of total DNA) to a 0.2 mL PCR tube along with 0.5 µL of the appropriate type IIS restriction enzyme, 1.0 µL of T4 DNA ligase enzyme, and 2.0 µL of T4 ligase buffer, and 0.8 µL of the PaqCI activator reagent if PaqCI was the enzyme used for assembly (all of these reagents from New England Biolabs; Ipswich, MA) were then added, along with enough sterile distilled water to bring the total volume to 20 µL. After mixing, assembly reactions were placed in a thermal cycler on a program cycling between 37°C for 5 minutes and

16°C for 10 minutes for 10-20 cycles, followed by a final 15-minute incubation at 37°C and a 10-minute inactivation step at 80°C and ending in an indefinite hold at 12°C.

For Gibson assembly schemes, input fragments were mixed in equimolar ratios totaling no more than 150 ng of total DNA in a 0.2 mL PCR tube with sterile distilled water bringing the total volume to 3 µL. This was then mixed with 3 µL of NEBuilder® HiFi DNA Assembly Master Mix (New England Biolabs) and the tube placed in a thermal cycler incubation the reaction for 30 minutes at 50°C, followed by an indefinite hold at 12°C as per the product protocol.

For restriction digestion/ligation cloning, insert and backbone fragments which had been digested with the appropriate enzymes following the manufacturer's recommendation (New England Biolabs, in all cases) and purified from agarose gel as described above were added in a 2:1 molar ratio (not exceeding 50 ng total DNA) to a 0.2 mL PCR tube with sterile distilled water bringing the total volume to 3 µL. This was then mixed with 3 µL of Instant Sticky-end Ligase Master Mix (New England Biolabs), pipette mixed for 8-10 strokes, and placed on ice according to the product protocol.

2 µL of each completed reaction mixture was used for transformation of chemically competent cells (or 1 µL was used for electroporation) using *E. coli* strains and on media and conditions described under the "Strains, media, and growth conditions" heading in the main text methods. Whenever the transformed plasmid carried a visual reporter, colonies were selected from Petri plates based on screening for its expression. Candidate colonies were used to inoculate two 3-mL cultures for each plasmid. to inoculate two 3-mL cultures for each plasmid. The ZR Plasmid Miniprep – Classic Kit (Zymo Research) was used to extract plasmid DNA following overnight growth, and clones were sequence-confirmed by Oxford-Nanopore based whole plasmid sequencing by Plasmidsaurus Inc. (Louisville, KY).

**Method S2**: Cloning and production of agrobacterial strain variants for evaluating visual reporters

Synthetic fragments produced for the promoters $P_{J23100}$, $P_{J23119}$, $P_{int}$, and $P_{Atu8024}$ (a suggestion from Roger Thilmony) and for the T3 and T7 terminator sequences were used to assemble a combinatorial expression cassette series for evaluation of the set of promoter elements. Each promoter was assembled with coding sequences for the sfGFP and eforRed markers, both amplified by PCR with an *Agrobacterium*-specific Shine-Dalgarno sequence (Golshani *et al.*, 2000) introduced via the forward primer for each, while the T3 terminator was paired with sfGFP and the T7 terminator with eforRed. The ends of all fragments contained *PaqCI* recognition sites and overhangs enabling assembly of expression cassettes in an agrobacteria-compatible (VS1 replicon) recipient plasmid backbone with spectinomycin resistance (pVS-rec [S]).

These constructs were generated by golden gate assembly with the PaqCI enzyme following the protocol described in Method S1 and were named pVS-sfGFP-[1-4] and pVS-eforRed-[1-4] with 1-4 corresponding to each cassette being driven by promoters $P_{J23100}$, $P_{J23119}$, $P_{int}$, and $P_{Atu8024}$, respectively. Additionally, the original cassettes for sfGFP and eforRed contained in the plasmids pEN-L1-PJ23119-PplpT-sfGFP-TrrfB-BsaI-Scaf-L2 and "eforRed chromoprotein", respectively, which had been designed for expression in *E. coli*, but failed to express when deployed in agrobacteria were amplified in their entirety and cloned into the same pVS-rec [S] backbone, yielding the control plasmids pVS-sfGFP-0 and pVS-eforRed-0. The primer sequences used in assembly of these constructs are given in Dataset S3. Each of these constructs was introduced into each of three chromosomally diverse agrobacterial strain backgrounds (AGL1, biovar 1; CM65/95, biovar 2, and T268/95, biovar 3), creating a strain by promoter combinatorial library, which was then utilized in "quantitative analysis of reporter expression" as described under that heading in the main text methods and additionally in Method S10 below.

Alternative visual reporter cassettes used in the AGROtools collection were assembled using the same methods and nomenclature described for sfGFP and eforRed. Synthetic coding sequence fragments for BFP (Bluebonnet2; Shroff *et al.*, 2020), amilCP, aeBlue, and tsPurple (Liljeruhm *et al.*, 2018) were each assembled with promoter $P_{int}$ and terminator T7 (with the exception of terminator rrnB T1 being used for BFP) to generate pVS-BFP-3, pVS-amilCP-3, pVS-aeBlue-3, and pVS-tsPurple-3. These were used as templates for downstream cloning.

**Method S3**: Assembly of all AGROtools plasmid components

The $P_{J23119}$::sfGFP::T3t and $P_{int}$::eforRed::T7t expression cassettes (in pVS-sfGFP-2 and pVS-eforRed-3) were selected to serve as the standard visual reporter parts for replacement and eviction sequences in AGROtools, respectively. These, along with all template-amplified toolkit components were purged of internal recognition sites for the set of type IIS restriction enzymes listed in Method S1, by site-directed mutagenesis using the set of primers listed in Dataset S4. In most cases, the resulting plasmid templates were named the same as their progenitor sequences with the term "-purged" appended. Due to the large size of the original plasmid template, the purged RK2 replicon sequence was re-assembled with a spectinomycin resistance marker yielding pRK2 (S), while this template was then used to generate a backbone for purging the gentamicin and chloramphenicol resistance gene markers, resulting in pRK2 (Ge) and pRK2 (Cm), respectively.

A series of 6 intermediate cloning constructs based on a homology arm design for disarming agrobacterial strains carrying type Ia Ti plasmids (named pDisarm_TiIa_v[1-6]) were assembled using *BsaI*-mediated golden gate assembly reactions with 7-8 fragment inputs each. Features including the ordered type IIP restriction sites intervening each molecular part and the pairs of recombinase sites flanking replacement sequence cassettes were incorporated into these constructs via 5' extensions to the primers used for amplifying each fragment. Each of these constructs differed in the individual molecular parts contained in their respective replacement and eviction sequences. The assembled plasmids described up to this point, along with the set of plasmids listed in Table S2 were then used as templates to construct all AGROtools modules using primers as described in Dataset S5.

pDisarm_TiIa_v[1-6] were used as templates to amplify cargos cloned into module 2 and module 4 recipient fragments. Additional variants of module 2 and module 4 components were constructed using restriction/ligation cloning to replace visual and selectable marker sequences by leveraging the designed restriction sites and inserting PCR-generated fragments containing new visual and selectable marker sequences digested with identical enzymes. Gibson assembly was used to construct some additional module 2 variants.

Backbone and excision plasmid components were also assembled using BsaI-mediated golden gate assembly reactions. For backbone components, the input fragments for each reaction consisted of a replicon, the canthaxanthin biosynthesis pathway gene cluster (split into 2 segments), a segment containing the origin of transfer sequence derived from plasmid pRK2 ($oriT_{pRK2}$), and an antibiotic resistance gene cassette. For excision plasmid components, the input fragments for each reaction consisted of a segment containing the BBR replicon and a *sacB* gene expression cassette, an antibiotic resistance gene cassette, a tyrosine recombinase (Cre or FLP) expression cassette, and an eforRed visual reporter expression cassette.

Modular plasmid variants carrying the CRISPR/dCas9 base editor cassette, the wild-type SpCas9 expression cassette, the INTEGRATE CRISPR-associated transposase polycistron, sgRNA or crRNA spacer sequence entry cassettes, INTEGRATE transposon cargo entry fragment, or recombineering operons, were each generated by BsaI-mediated (or BbsI-mediated, when the resulting plasmid contained *BsaI* sites) golden gate assembly.

Base plasmids for modules 1-4 were assembled by ligation of a short fragment carrying a designed multiple cloning site generated by oligonucleotide annealing into a BsaI-digested recipient fragment for each module. New insert entry variants for each of the modules were generated by BsaI-mediated golden gate assembly using parts consisting of each module backbone, a *lacZα* fragment, and a *ccdB* toxin gene fragment. These assembly mixtures were transformed into *E. coli* DB3.1 (ccdB survival) cells by electroporation and then plated on LB solid media amended with carbenicillin, IPTG (250 μg/mL, and X-gal (40 μg/mL). Blue colonies formed on these plates were selected as clone candidates.

**Method S4**: Assembly of disarmament and modification constructs

Target strains for disarmament were selected based on their genome having already been assembled, and on their genomic diversity, based on both chromosomal ancestry and virulence plasmid type. Constructs for facilitating disarmament or a chromosomal modification were designed by first identifying suitable left and right homology arm (HA) sequences, each between 800 and 1,400 bp in length, from genomic sites flanking the target sequence to be deleted. Assembled genomes were first annotated using the 'Beav' software package (Jung *et al.*, 2024). The resulting Genbank files were then imported into Geneious Prime 2026.1.2 for further analysis. Sequences repeated in the genome were annotated using the 'Repeat Finder' software plugin for Geneious (Volfovsky *et al.*, 2001). Gene sequences near the genomic regions flanking the targets that were marked as repeats, or which were annotated either as transposable elements or tyrosine recombinase family members were avoided and the HA window shifted to minimize the possibility of off-target homologous recombination. HA candidate sequences were then used as a query to search against a custom BLAST database consisting of the target strain's whole genome sequence to ensure that there were no partially overlapping sequences present elsewhere in the genome. Primers pairs were designed for amplifying each HA segment from each target strain's genomic DNA (or that of another strain carrying a closely related Ti plasmid). 5' extensions incorporating PaqCI recognition sites and the unique 4-base overhangs required for positional assembly within the constructs were added to each of these primers. PCR with Q5 polymerase (using the modified recipe detailed above) was performed with each of the primer pairs using cells of the target strain directly as reaction template. Fragments were then gel-purified as described in Method S1. For construct assembly, approximately 30 ng of each purified PCR product for the left and right HAs was added to a 0.2 mL PCR tube along with approximately 150 ng each of selected module 2 and module 4 components and approximately 400 ng of a selected backbone component (typically pAGRObb_000). The golden gate assembly reaction was carried out using parameters described above. Reaction mixtures were transformed into *E. coli*, colonies selected, plasmids extracted, and clones sequence confirmed as described previously.

Disarmament or modification constructs were named according to their purpose, the target strain, and the suffixes of the specific module 2, module 4, and backbone plasmids used to generate them, each separated by underscore characters, following the format: [pDisarm/pDelete]_[strain name]_[if applicable, target gene name]_[module 2 suffix]_[module 4 suffix]_[backbone suffix]. Sequences for each homology arm amplification primer used are described in Dataset S6, while Dataset S7 shows the names of the specific disarmament or modification constructs used for targeting each strain.

**Method S5**: Construct introduction into agrobacteria cells by electroporation

Preparation of agrobacterial competent cells was performed similar to that described in Nair *et al.* (2003). Briefly, a single colony was used to inoculate a 3 mL culture in MGY broth, which was then grown to a density of about $OD_{600}$=1.0. 1 mL of culture was then spun down and the pellet washed four times in sterile distilled water before resuspending it in 50 µL of water. Approximately 250 ng of construct was then mixed with these cells and electroporation was performed at a 1.85 kV setting using a T-720 Transporator Plus Electropulsator (BTX; Holliston, MA) in a cuvette with a 0.1 cm gap length. 1 mL of MGY broth was then added immediately and cultures were subjected to a 2-3 hour outgrowth period before being plated. For plating, cultures were spun down for 30 seconds, 900 µL of supernatant was removed, the pellet was resuspended in the remaining supernatant, and then the entire culture was plated on MGY solid media amended with the appropriate antibiotic for selection of primary recombinant clones after 2-3 days of incubation. Electroporations in strain Ach5 deviated from this protocol in that all of a 5 mL overnight culture was harvested per electroporated sample.

**Method S6**: Construct introduction into agrobacteria cells by triparental mating

We employed a triparental mating protocol modified from that described by Wise *et al.* (2006). Briefly, a donor *E. coli* strain carrying the disarmament construct, the helper *E. coli* strain (HB101 with pRK2013), and the target agrobacterial strain were each grown to saturation in liquid cultures and then used to inoculate secondary cultures, which were harvested when they reached $OD_{600}$=0.6-1.0. For all three cultures, 1-mL aliquots were washed once in a sterile 0.8% NaCl solution, resuspended in 100 µL of sterile distilled water, and 100 µL of each was mixed in equal proportions before pipetting the entire volume of the mixture onto a sterile 80-mm cellulose filters placed on an MGY agar plate without antibiotics. After approximately 48 hours of incubation at

28°C, the culture was collected by vortex mixing the cellulose filter in a 50-mL Falcon tube with 5 mL of sterile 0.8% NaCl solution. A 1-mL aliquot of this was then pelleted, the supernatant was removed and then the pellet resuspended in 100 µL of sterile distilled water. This entire volume was then plated on MG solid media (MGY recipe sans the yeast extract) amended with the appropriate antibiotic for selection of primary recombinant clones after 2-3 days of incubation. Since both HB101 and TOP10 (10β) strain backgrounds are auxotrophic for leucine biosynthesis, they are unable to grow on media without a source of free amino acids (*i.e.*, yeast extract). Thus, the recipient strain is efficiently selected to the exclusion of the donor and helper strain backgrounds following this method.

**Method S7**: Detailed plate reader assay

Each of the strain derivatives carrying the sfGFP and eforRed expression cassette variants described in Method S2 (30 total) were streaked from glycerol stocks onto fresh MGY media plates amended with 150 µg/mL of spectinomycin. Sectors of these plates containing single colonies were microscope imaged using green or red fluorescence filter sets, respectively.

For the sfGFP expression series (15 total), colonies from each culture were scraped from plates and suspended in 1 mL of MGY broth. The density of each of these suspensions was then adjusted to roughly $OD_{600}$=0.1 and 100-µL aliquots of each were transferred in triplicate sets into a 96-well plate along with suspensions of empty vector strains and MGY broth only as blank controls. Fluorescence intensity data for the entire plate was then collected using the microplate reader described in the main text methods. The raw measurements for each well taken by the microplate reader are provided in Dataset S8. Intensity values for each well were normalized according to their respective measured inoculum density readings. Readings for the three wells in each treatment were averaged and then baseline adjusted to readings acquired for empty vector strains. Scale bars show standard deviation across technical replicates.

Characterization of promoter elements was used to inform how expression cassettes were constructed as molecular parts incorporated into AGROtools. To avoid repeated use of the same expression elements in assembled constructs, we assigned particular promoter and terminator elements to each type of molecular part. The $P_{J23119}$::sfGFP::T3t and $P_{int}$::eforRed::T7t cassettes were selected as the primary sequences for inclusion in the replacement and eviction sequences cargos, respectively, whereas $P_{J23100}$ paired with rrnB T1 was used to drive each of the alternative counterselection marker sequences. $P_{Atu8024}$ was used to drive the *recA*$_{C58}$ sequence included in some eviction sequence cargos.

**Method S8**: Disarmament of wild-type strains and marker excision

Primary recombinant clones forming on plates following construct introduction into a target strain were screened for expression of both visual reporters using the NIGHTSEA fluorescence visualization systems. Colonies positive for both green and red fluorescence were then suspended in 100 µL of MGY broth, and then 50 µL of this culture was plated on MGY media amended with the corresponding selective antibiotic and sucrose at 2.5% w/v to select for secondary recombinant clones after 2-3 additional days of incubation. Apparent double crossover disarmed colonies formed on this plate (positive for green fluorescence, but negative for red fluorescence) were validated as disarmed by the presence of 4-5 kbp bands after colony PCR using “flanking” primers specifically designed for this purpose (see Dataset S9 for specific flanking primer sequences and Figure 3 for a visual representation of the strategy).

Strain derivatives confirmed as disarmed were then transformed with an AGROtools plasmid component of the “excision” class (pAGROex) appropriate for removing the pair of replacement sequence marker cassettes (see Dataset S7 for the specific pAGROex plasmid used for marker excision in each strain) either using electroporation or triparental mating as described above. Selection for these marker-excised derivative clones required plating on MGY (or MG) media amended with only the antibiotic selecting for maintenance of the excision plasmid and then screening for colonies which are negative for green fluorescence, but positive for red fluorescence after 2-3 days of incubation. These colonies were then suspended in 100 µL of MGY broth and 50 µL of this culture was plated on MGY media amended only with sucrose at 2.5% w/v to evict the excision plasmid. After 2-3 days of incubation, colonies which did not express either of the fluorescent signals were selected and successful marker excision was validated by colony PCR, again using a corresponding “flanking” primer pair, which should now yield a shorter (2-3 kbp) band than the progenitor disarmed strain still carrying the marker cassettes.

**Method S9**: Genome assembly and analysis of modified strains

Raw Oxford-Nanopore reads for each whole genome sequencing sample obtained from Plasmidsaurus were imported into Geneious Prime for assembly and analysis. Reads were trimmed and those <1,000 bp filtered out using the plugin for BBDuk with default settings. If needed, the read sets were downsampled to approximately 40X coverage using BBNorm (Bushnell *et al.*, 2017). De novo assembly was performed on the resulting read set using Flye (Kolmogorov *et al.*, 2019). Resulting contigs for each sample were annotated with Beav. The residual fragment containing the recombinase site used in the disarmament process for each strain was searched against the genome assemblies, with only one match for this sequence being found in each strain’s genome. Sequences of other individual molecular parts used in the introduced disarmament construct were also used as queries for BLAST searches, but no positive hits were found for any strain genome.

**Method S10**: Quantitative assay of reporter expression from agroinfiltrated disarmed strains

Disarmed strains carrying binary plasmid pT200-GFP-HygR (sequence given in Dataset S10) were infiltrated into 3-4 week old *N. benthamiana* plants as described in the main text methods. Each strain was injected into a single leaf on three different plants. Four leaf disks measuring approximately 6 mm were excised from each infiltrated leaf (12 total replicates per strain; 3 biological replicates and 4 technical replicates) and placed adaxial side up in microplate wells containing 100 µL sterile deionized $H_2O$. The plate was then run on the microplate reader described in the main text methods, with four measurements taken per well. The average well intensity values for each sample were recorded for 12 total measurements per strain. Mean and standard deviation were calculated, and any measurement which was over 2.5 standard deviations from the mean was dropped. ANOVA residual normality and homogeneity of variance were assessed with Shapiro-Wilk and Levene's tests, respectively. Because these assumptions were not met, statistical analysis of strain backgrounds was implemented using the Kruskal-Wallis non-parametric omnibus rank-sum test followed by post-hoc Dunn's pairwise comparisons and Benjamini-Hochberg correction for multiple hypothesis testing. Compact letters displayed in Figure S5 were derived from Benjamini-Hochberg adjusted Dunn *p*-values ($\alpha = 0.05$). Strains sharing a letter showed no significant difference (adjusted *p*-value > 0.05). Blank controls were retained for data visualization but excluded from statistical analyses.

**Method S11**: Counterselection agent/marker testing assay

Three constructs for C58 disarmament were assembled as previously described using the standard left and right homology arm fragments flanking the T-DNA on pTiC58, pAGROm2_112, pAGRObb_000, and either pAGROm4_210, pAGROm4_220, or pAGROm4_230. The final constructs were named pDisarm_C58_112_210_000, pDisarm_C58_112_220_000, and pDisarm_C58_112_230_000 differed only as to whether they carried the *sacB*, *pheS**, or *codA*$^{D314A}$ counterselection marker genes in their replacement sequences. Each of these constructs was introduced into the C58 wild-type background by electroporation to obtain a set of primary recombinant derivatives. Secondary recombinant clones were also obtained by applying sucrose counterselection to primary clones carrying *sacB*.

Each of the primary recombinant derivatives were streaked in triplicate onto fresh media plates alongside triplicate secondary recombinant clones, which functioned as a control treatment. After 2 days of growth, cells from each culture were scraped from plates and suspended in 1 mL of MGY broth. The density of each of these suspensions was then adjusted to $OD_{600}$=0.2 and 50 µL of each culture was transferred to each of the 12 wells in the top row of a 96-well plate. The wells below each sample were then used for a 10X serial dilution series relative to the contents of the well above, so that suspensions in the bottom row had a concentration of $1 \times 10^{-7}$ relative to that of the corresponding well in the top row.

A 96-pin plate replicator tool was then used to transfer this sample set onto a series of 150 mm x 10 mm Petri plates containing MGY solid media amended with 150 µg/mL spectinomycin and reagents corresponding to each counterselection marker gene: sucrose at concentrations of 30, 75, and 150 mM; 4-chlorophenylalanine (4-CP; Santa Cruz Biotechnology; Dallas, TX) at concentrations of 1, 5, and 20 mM; 5-fluorocytosine (5-FC; Tokyo Chemical Industry; Tokyo, Japan) at concentrations of 0.5, 2, and 5 mM. Plates amended with only spectinomycin served as a control treatment. Duplicate plates were made for each individual media formulation. Plates were incubated for 2 days at 28$^{o}$C, when at least some growth was apparent in the top row for each of the clones affected by the corresponding counterselection agent. Images of plates were collected in bright field, green fluorescence, and red fluorescence channels on sectors of 4 x 4 spots that included one clone of each of the three different single recombinants and a double-crossover clone at $1 \times 10^{0}$, $1 \times 10^{-1}$, $1 \times 10^{-2}$, and $1 \times 10^{-3}$ concentrations.

**Method S12**: Assessment of dual paired reporter constructs for disarmament

To test the utility of the visual reporters in selecting correctly recombined colonies, we produced constructs differing from our typical design only in the absence of expression cassettes for both visual reporters. To accomplish this, we first generated the accessory plasmids (not included in the AGROtools set) pAGROm2_002 and pAGROm4_010 using primers and strategies detailed in Dataset S11. Standard and visual reporter-free disarmament construct variants were generated for strains C58 and Ach5, named pDisarm_C58_112_210_000, pDisarm_C58_002_010_000, pDisarm_Ach5_102_210_000, and pDisarm_Ach5_002_010_000. These were each introduced into their respective wild-type target strains by electroporation.

For each construct-transformed treatment, 10 possible primary recombinant colonies were re-streaked to fresh media prior to colony PCR validation to ameliorate false positive signal due to residual construct DNA remaining on the selection plate. Colony selection was either guided by the expression of both fluorescent reporters or selected randomly in the treatments for which these were absent. The primers specR_seq-F and specR_seq-R were used for colony PCR, resulting in a 256 bp band for confirmed recombinant colonies (sequences in Dataset S11). PCR-confirmed primary recombinant clones were then pooled (within each treatment) and resuspended in 100 µL of MGY broth and then plated to MGY plates containing sucrose at 2.5% w/v to filter for secondary recombinants. 10 possible secondary recombinant colonies then underwent PCR validation using the pTiC58_Left_flanking-F + pTiC58_Right_flanking-R and pTiAch5_Left_flanking-F + pTiAch5_Right_flanking-R primer pairs (Dataset S9) for C58 and Ach5 derivatives, respectively.

**Method S13**: Homology arm length evaluation

The forward primers pTiC58_Left_HA-1200F, pTiC58_Left_HA-600F, pTiC58_Left_HA-300F were each paired with the reverse primer pTiC58_Left_HA-R to produce left homology arm fragments for C58 disarmament which were approximately 1,200 bp, 600 bp, and 300 bp in length respectively. Right homology arms of the same lengths were generated using the pTiC58_Right_HA-F forward primer paired with each of the pTiC58_Right_HA-1200R, pTiC58_Right_HA-600R, and pTiC58_Right_HA-300R reverse primers (sequences in Dataset S6). Homology arm sets of the roughly equivalent lengths were then input into three separate assemblies with pAGROm2_112, pAGROm4_210, and pAGRObb_000 to generate the constructs pDisarm_C58_112_210_000 [1200 HA], pDisarm_C58_112_210_000 [600 HA], and pDisarm_C58_112_210_000 [300 HA], respectively. These were then introduced in parallel into the C58 wild-type strain by electroporation in molar equivalent quantities to account for differences in construct size. Primary recombinant selection plates for each treatment were imaged after 3 days of incubation.

**Method S14**: Knockout of *recF* in the AGL1 background

Homology arms flanking the *recF* chromosomal locus were amplified from strain AGL1 colonies and used to assemble two constructs differing only in whether they contained a *recA* ectopic expression cassette in the eviction sequence. pDelete_C58_recF_112_211_000 carried *recA*, while pDelete_C58_recF_112_210_000, lacking *recA*, functioned as a negative control. The constructs were introduced into the AGL1 strain background by electroporation. Primary recombinant colonies were obtained only for the pDelete_C58_recF_112_211_000 construct. These were carried forward through the process of obtaining secondary recombinants and marker excision as described above to obtain the resulting derivative AGL1 *recF*- [ME].

**Method S15**: Gene deletion in *Pseudomonas*

Homology arms approximately 500 bp in length flanking the target gene (locus tag PSPTO_5060) in *Pseudomonas syringae* pv. *tomato* DC3000 were PCR-amplified and assembled with pAGROm2_101, pAGROm4_210, and pAGRObb_000 into a gene deletion construct using methods described above. This construct was introduced into DC3000 cells by triparental mating using King's B medium (King *et al.*, 1954) and maintained at 28°C. Primary recombinant clones were identified by eforRed expression after 2 days on kanamycin selection at 50 μg/mL. These were then transferred to media containing 5% sucrose to recover secondary recombinants. Deletion of the target operon was confirmed by colony PCR with flanking primers (sequences in Dataset S9).

| Plasmid short name | Plasmid contents | Antibiotic / other amendments to media | Use cases / description / notes | Sequence |
|---|---|---|---|---|
| pAGROm1_000 | module1_[base with MCS linker] | carb100 | Assembling a construct with an empty module 1 slot; Inserting a [left] homology arm into module 1 slot fo | TTGAGATCCTTTTT[illegible] |
| pAGROm1_000L | module1_[base with lacZa-ccdB placeholder] | carb200, IPTG250, X-gal50 | Blue/white cloning compatible with IPTG and X-gal added to media plate; Inserting a [left] homology arm i | TTGAGATCCTTTTT[illegible] |
| pAGROm2_000 | module2_[base with MCS linker] | carb100 | Assembling a construct with an empty module 2 slot; Introducing a replacement sequence or other new tc | TTGAGATCCTTTTT[illegible] |
| pAGROm2_000L | module2_[base with lacZa-ccdB placeholder] | carb200, IPTG250, X-gal50 | Blue/white cloning compatible with IPTG and X-gal added to media plate; Introducing a replacement seque | TTGAGATCCTTTTT[illegible] |
| pAGROm3_000 | module3_[base with MCS linker] | carb100 | Assembling a construct with an empty module 3 slot; Inserting a [right] homology arm into module 3 slot f | TTGAGATCCTTTTT[illegible] |
| pAGROm3_000L | module3_[base with lacZa-ccdB placeholder] | carb200, IPTG250, X-gal50 | Blue/white cloning compatible with IPTG and X-gal added to media plate; Inserting a [right] homology arm | TTGAGATCCTTTTT[illegible] |
| pAGROm4_000 | module4_[base with MCS linker] | carb100 | Assembling a construct with an empty module 4 slot; Introducing an eviction sequence or other new toolk | TTGAGATCCTTTTT[illegible] |
| pAGROm4_000L | module4_[base with lacZa-ccdB placeholder] | carb200, IPTG250, X-gal50 | Blue/white cloning compatible with IPTG and X-gal added to media plate; Introducing an eviction sequenc | TTGAGATCCTTTTT[illegible] |
| pAGROm2_101 | module2_FRT^sfGFP-kanR^FRT | carb100, kan50 | "Replacement sequence" cargo for assembling a homologous recombination-based disarmament or deleti | TTGAGATCCTTTTT[illegible] |
| pAGROm2_102 | module2_FRT^sfGFP-specR^FRT | carb100, spec100 | "Replacement sequence" cargo for assembling a homologous recombination-based disarmament or deleti | TTGAGATCCTTTTT[illegible] |
| pAGROm2_103 | module2_FRT^sfGFP-gentR^FRT | carb100, gent10 | "Replacement sequence" cargo for assembling a homologous recombination-based disarmament or deleti | TTGAGATCCTTTTT[illegible] |
| pAGROm2_104 | module2_FRT^sfGFP-apraR^FRT | carb100, apra50 | "Replacement sequence" cargo for assembling a homologous recombination-based disarmament or deleti | TTGAGATCCTTTTT[illegible] |
| pAGROm2_111 | module2_LoxP^sfGFP-kanR^LoxP | carb100, kan50 | "Replacement sequence" cargo for assembling a homologous recombination-based disarmament or deleti | TTGAGATCCTTTTT[illegible] |
| pAGROm2_112 | module2_LoxP^sfGFP-specR^LoxP | carb100, spec100 | "Replacement sequence" cargo for assembling a homologous recombination-based disarmament or deleti | TTGAGATCCTTTTT[illegible] |
| pAGROm2_113 | module2_LoxP^sfGFP-gentR^LoxP | carb100, gent10 | "Replacement sequence" cargo for assembling a homologous recombination-based disarmament or deleti | TTGAGATCCTTTTT[illegible] |
| pAGROm2_114 | module2_LoxP^sfGFP-apraR^LoxP | carb100, apra50 | "Replacement sequence" cargo for assembling a homologous recombination-based disarmament or deleti | TTGAGATCCTTTTT[illegible] |
| pAGROm2_121 | module2_Lox2272^sfGFP-kanR^Lox2272 | carb100, kan50 | "Replacement sequence" cargo for assembling a homologous recombination-based disarmament or deleti | TTGAGATCCTTTTT[illegible] |
| pAGROm2_122 | module2_Lox2272^sfGFP-specR^Lox2272 | carb100, spec100 | "Replacement sequence" cargo for assembling a homologous recombination-based disarmament or deleti | TTGAGATCCTTTTT[illegible] |
| pAGROm2_123 | module2_Lox2272^sfGFP-gentR^Lox2272 | carb100, gent10 | "Replacement sequence" cargo for assembling a homologous recombination-based disarmament or deleti | TTGAGATCCTTTTT[illegible] |
| pAGROm2_124 | module2_Lox2272^sfGFP-apraR^Lox2272 | carb100, apra50 | "Replacement sequence" cargo for assembling a homologous recombination-based disarmament or deleti | TTGAGATCCTTTTT[illegible] |
| pAGROm2_131 | module2_Lox5171^sfGFP-kanR^Lox5171 | carb100, kan50 | "Replacement sequence" cargo for assembling a homologous recombination-based disarmament or deleti | TTGAGATCCTTTTT[illegible] |
| pAGROm2_132 | module2_Lox5171^sfGFP-specR^Lox5171 | carb100, spec100 | "Replacement sequence" cargo for assembling a homologous recombination-based disarmament or deleti | TTGAGATCCTTTTT[illegible] |
| pAGROm2_133 | module2_Lox5171^sfGFP-gentR^Lox5171 | carb100, gent10 | "Replacement sequence" cargo for assembling a homologous recombination-based disarmament or deleti | TTGAGATCCTTTTT[illegible] |
| pAGROm2_134 | module2_Lox5171^sfGFP-apraR^Lox5171 | carb100, apra50 | "Replacement sequence" cargo for assembling a homologous recombination-based disarmament or deleti | TTGAGATCCTTTTT[illegible] |
| pAGROm2_301 | module2_FRT^BFP-kanR^FRT | carb100, kan50 | "Replacement sequence" cargo for assembling a homologous recombination-based disarmament or deleti | TTGAGATCCTTTTT[illegible] |
| pAGROm2_311 | module2_LoxP^BFP-kanR^LoxP | carb100, kan50 | "Replacement sequence" cargo for assembling a homologous recombination-based disarmament or deleti | TTGAGATCCTTTTT[illegible] |
| pAGROm2_321 | module2_Lox2272^BFP-kanR^Lox2272 | carb100, kan50 | "Replacement sequence" cargo for assembling a homologous recombination-based disarmament or deleti | TTGAGATCCTTTTT[illegible] |
| pAGROm2_331 | module2_Lox5171^BFP-kanR^Lox5171 | carb100, kan50 | "Replacement sequence" cargo for assembling a homologous recombination-based disarmament or deleti | TTGAGATCCTTTTT[illegible] |
| pAGROm4_210 | module4_eforRed-sacB | carb100 | "Eviction sequence" cargo for assembling a homologous recombination-based disarmament, deletion, or r | TTGAGATCCTTTTT[illegible] |
| pAGROm4_211 | module4_eforRed-sacB-recA$_{C58}$ | carb100 | "Eviction sequence" cargo for assembling a homologous recombination-based disarmament, deletion, or r | TTGAGATCCTTTTT[illegible] |
| pAGROm4_220 | module4_eforRed-pheS | carb100 | "Eviction sequence" cargo for assembling a homologous recombination-based disarmament, deletion, or r | TTGAGATCCTTTTT[illegible] |
| pAGROm4_221 | module4_eforRed-pheS-recA$_{C58}$ | carb100 | "Eviction sequence" cargo for assembling a homologous recombination-based disarmament, deletion, or r | TTGAGATCCTTTTT[illegible] |
| pAGROm4_230 | module4_eforRed-codA | carb100 | "Eviction sequence" cargo for assembling a homologous recombination-based disarmament, deletion, or r | TTGAGATCCTTTTT[illegible] |
| pAGROm4_231 | module4_eforRed-codA-recA$_{C58}$ | carb100 | "Eviction sequence" cargo for assembling a homologous recombination-based disarmament, deletion, or r | TTGAGATCCTTTTT[illegible] |
| pAGRObb_000 | colE1_chlorR_oriT$_{RK2}$ [CXN placeholder] | chlor25 | Non-replicating ("suicide") backbone for assembling any type of construct. Contains a chloramphenicol res | TTGAGATCCTTTTT[illegible] |
| pAGRObb_001 | colE1-kanR-oriT$_{RK2}$ [CXN placeholder] | kan50 | Non-replicating ("suicide") backbone for assembling any type of construct. Contains a kanamycin resistanc | TTGAGATCCTTTTT[illegible] |
| pAGRObb_002 | colE1_specR_oriT$_{RK2}$ [CXN placeholder] | spec100 | Non-replicating ("suicide") backbone for assembling any type of construct. Contains a spectinomycin resist | TTGAGATCCTTTTT[illegible] |
| pAGRObb_003 | colE1_gentR_oriT$_{RK2}$ [CXN placeholder] | gent20 | Non-replicating ("suicide") backbone for assembling any type of construct. Contains a gentamicin resistanc | TTGAGATCCTTTTT[illegible] |
| pAGRObb_004 | colE1_apraR_oriT$_{RK2}$ [CXN placeholder] | apra50 | Non-replicating ("suicide") backbone for assembling any type of construct. Contains a apramycin resistanc | TTGAGATCCTTTTT[illegible] |
| pAGRObb_101 | pBBR-kanR-oriT$_{RK2}$ [CXN placeholder] | kan50 | Backbone with BBR replicon for assembling any type of construct. Contains a kanamycin resistance marker | GCGGCCACCGGC[illegible] |
| pAGRObb_102 | pBBR-specR-oriT$_{RK2}$ [CXN placeholder] | spec100 | Backbone with BBR replicon for assembling any type of construct. Contains a spectinomycin resistance ma | GCGGCCACCGGC[illegible] |

| | | | | |
|---|---|---|---|---|
| pAGRObb_103 | pBBR-gentR-oriT$_{RK2}$ [CXN placeholder] | gent10 | Backbone with BBR replicon for assembling any type of construct. Contains a gentamicin resistance marke | GCGGCCACCGGCTGGCTCGCTTCGCTCGGCCCGTGGACAACCCTG[illegible] |
| pAGRObb_104 | pBBR-apraR-oriT$_{RK2}$ [CXN placeholder] | apra50 | Backbone with BBR replicon for assembling any type of construct. Contains a apramycin resistance marker | GCGGCCACCGGCTGGCTCGCTTCGCTCGGCCCGTGGACAACCCTG[illegible] |
| pAGRObb_201 | pRK2-kanR-oriT$_{RK2}$ [CXN placeholder] | kan50 | Backbone with RK2 replicon for assembling any type of construct. Contains a kanamycin resistance marker | GGCCGCCGGCGTTGTGGATACCTCGCGGAAAACTTGGCCCTCACT[illegible] |
| pAGRObb_202 | pRK2-specR-oriT$_{RK2}$ [CXN placeholder] | spec100 | Backbone with RK2 replicon for assembling any type of construct. Contains a spectinomycin resistance ma | GGCCGCCGGCGTTGTGGATACCTCGCGGAAAACTTGGCCCTCACT[illegible] |
| pAGRObb_203 | pRK2-gentR-oriT$_{RK2}$ [CXN placeholder] | gent10 | Backbone with RK2 replicon for assembling any type of construct. Contains a gentamicin resistance marke | GGCCGCCGGCGTTGTGGATACCTCGCGGAAAACTTGGCCCTCACT[illegible] |
| pAGRObb_204 | pRK2-apraR-oriT$_{RK2}$ [CXN placeholder] | apra50 | Backbone with RK2 replicon for assembling any type of construct. Contains a apramycin resistance marker | GGCCGCCGGCGTTGTGGATACCTCGCGGAAAACTTGGCCCTCACT[illegible] |
| pAGRObb_301 | pVS1-kanR-oriT$_{RK2}$ [CXN placeholder] | kan50 | Backbone with VS1 replicon (and colE1 for *E. coli* replication) for assembling any type of construct. Contair | TTGAAATCCTTTTTTTCTGCGCGTAATCTGCTGCTTGCAAACAAAAA[illegible] |
| pAGRObb_302 | pVS1-specR-oriT$_{RK2}$ [CXN placeholder] | spec100 | Backbone with VS1 replicon (and colE1 for *E. coli* replication) for assembling any type of construct. Contair | TTGAAATCCTTTTTTTCTGCGCGTAATCTGCTGCTTGCAAACAAAAA[illegible] |
| pAGRObb_303 | pVS1-gentR-oriT$_{RK2}$ [CXN placeholder] | gent10 | Backbone with VS1 replicon (and colE1 for *E. coli* replication) for assembling any type of construct. Contair | TTGAAATCCTTTTTTTTCTGCGCGTAATCTGCTGCTTGCAAACAAAA[illegible] |
| pAGRObb_304 | pVS1-apraR-oriT$_{RK2}$ [CXN placeholder] | apra50 | Backbone with VS1 replicon (and colE1 for *E. coli* replication) for assembling any type of construct. Contair | TTGAAATCCTTTTTTTCTGCGCGTAATCTGCTGCTTGCAAACAAAAA[illegible] |
| pAGRObb_401 | pSa-kanR-oriT$_{RK2}$ [CXN placeholder] | kan50 | Backbone with Sa replicon for assembling any type of construct. Contains a kanamycin resistance marker a | CCATCCAACAGCCCGCCGTCGAGCGGGCTTTTTTATCCCCGGAAGC[illegible] |
| pAGRObb_402 | pSa-specR-oriT$_{RK2}$ [CXN placeholder] | spec100 | Backbone with Sa replicon for assembling any type of construct. Contains a spectinomycin resistance mark | CCATCCAACAGCCCGCCGTCGAGCGGGCTTTTTTATCCCCGGAAGC[illegible] |
| pAGRObb_403 | pSa-gentR-oriT$_{RK2}$ [CXN placeholder] | gent10 | Backbone with Sa replicon for assembling any type of construct. Contains a gentamicin resistance marker a | CCATCCAACAGCCCGCCGTCGAGCGGGCTTTTTTATCCCCGGAAGC[illegible] |
| pAGRObb_404 | pSa-apraR-oriT$_{RK2}$ [CXN placeholder] | apra50 | Backbone with Sa replicon for assembling any type of construct. Contains a apramycin resistance marker a | CCATCCAACAGCCCGCCGTCGAGCGGGCTTTTTTATCCCCGGAAGC[illegible] |
| pAGROex_201 | pBBR-kanR-FLP-eforRed-sacB | kan50 | FLP recombinase expression cassette in a replicative backbone with a BBR replicon containing a kanamycir | GCGGCCACCGGCTGGCTCGCTTCGCTCGGCCCGTGGACAACCCTG[illegible] |
| pAGROex_202 | pBBR-specR-FLP-eforRed-sacB | spec100 | FLP recombinase expression cassette in a replicative backbone with a BBR replicon containing a spectinom | GCGGCCACCGGCTGGCTCGCTTCGCTCGGCCCGTGGACAACCCTG[illegible] |
| pAGROex_203 | pBBR-gentR-FLP-eforRed-sacB | gent10 | FLP recombinase expression cassette in a replicative backbone with a BBR replicon containing a gentamici | GCGGCCACCGGCTGGCTCGCTTCGCTCGGCCCGTGGACAACCCTG[illegible] |
| pAGROex_204 | pBBR-apraR-FLP-eforRed-sacB | apra50 | FLP recombinase expression cassette in a replicative backbone with a BBR replicon containing an apramyci | GCGGCCACCGGCTGGCTCGCTTCGCTCGGCCCGTGGACAACCCTG[illegible] |
| pAGROex_211 | pBBR-kanR-Cre-eforRed-sacB | kan50 | Cre recombinase expression cassette in a replicative backbone with a BBR replicon containing a kanamycir | GCGGCCACCGGCTGGCTCGCTTCGCTCGGCCCGTGGACAACCCTG[illegible] |
| pAGROex_212 | pBBR-specR-Cre-eforRed-sacB | spec100 | Cre recombinase expression cassette in a replicative backbone with a BBR replicon containing a spectinom | GCGGCCACCGGCTGGCTCGCTTCGCTCGGCCCGTGGACAACCCTG[illegible] |
| pAGROex_213 | pBBR-gentR-Cre-eforRed-sacB | gent10 | Cre recombinase expression cassette in a replicative backbone with a BBR replicon containing a gentamici | GCGGCCACCGGCTGGCTCGCTTCGCTCGGCCCGTGGACAACCCTG[illegible] |
| pAGROex_214 | pBBR-apraR-Cre-eforRed-sacB | apra50 | Cre recombinase expression cassette in a replicative backbone with a BBR replicon containing an apramyci | GCGGCCACCGGCTGGCTCGCTTCGCTCGGCCCGTGGACAACCCTG[illegible] |
| pAGROm2_412 | module2_LoxP^amilCP-specR^LoxP | carb100, spec100 | "Replacement sequence" cargo for assembling a homologous recombination-based disarmament or deleti | TTGAGATCCTTTTTTTCTGCGCGTAATCTGCTGCTTGCAAACAAAAA[illegible] |
| pAGROm2_512 | module2_LoxP^aeBlue-specR^LoxP | carb100, spec100 | "Replacement sequence" cargo for assembling a homologous recombination-based disarmament or deleti | TTGAGATCCTTTTTTTCTGCGCGTAATCTGCTGCTTGCAAACAAAAA[illegible] |
| pAGROm2_612 | module2_LoxP^tsPurple-specR^LoxP | carb100, spec100 | "Replacement sequence" cargo for assembling a homologous recombination-based disarmament or deleti | TTGAGATCCTTTTTTTCTGCGCGTAATCTGCTGCTTGCAAACAAAAA[illegible] |
| pAGROm4_410 | module4_amilCP-sacB | carb100 | "Eviction sequence" cargo for assembling a homologous recombination-based disarmament, deletion, or r | TTGAGATCCTTTTTTTCTGCGCGTAATCTGCTGCTTGCAAACAAAAA[illegible] |
| pAGROm4_510 | module4_aeBlue-sacB | carb100 | "Eviction sequence" cargo for assembling a homologous recombination-based disarmament, deletion, or r | TTGAGATCCTTTTTTTCTGCGCGTAATCTGCTGCTTGCAAACAAAAA[illegible] |
| pAGROm4_610 | module4_tsPurple-sacB | carb100 | "Eviction sequence" cargo for assembling a homologous recombination-based disarmament, deletion, or r | TTGAGATCCTTTTTTTCTGCGCGTAATCTGCTGCTTGCAAACAAAAA[illegible] |
| pAGROm2_102LC | module2_FRT^sfGFP-specR^FRT BsaI-[lacZa placeholder]-BsaI | carb100, spec100, IPTG250, X-gal50 | "Replacement sequence" entry plasmid intended for assembling a homologous recombination-based large | TTGAGATCCTTTTTTTCTGCGCGTAATCTGCTGCTTGCAAACAAAAA[illegible] |
| pAGROm2_112LC | module2_LoxP^sfGFP-specR^LoxP BsaI-[lacZa placeholder]-BsaI | carb100, spec100, IPTG250, X-gal50 | "Replacement sequence" entry plasmid intended for assembling a homologous recombination-based large | TTGAGATCCTTTTTTTCTGCGCGTAATCTGCTGCTTGCAAACAAAAA[illegible] |
| pAGROm1_int | module1_PJ23119-VchTnsABC-TniQ-Cas876 [INTEGRATE proteins] | carb100 | Module 1 containing an operon encoding the 7 proteins constituting the machinery used for INTEGRATE (( | TTGAGATCCTTTTTTTCTGCGCGTAATCTGCTGCTTGCAAACAAAAA[illegible] |
| pAGROm1_cbe0 | module1_pvirB-dCas9-CDA-UL-T3t | carb100 | Module 1 containing an acetosyringone-inducible cassette driven by an *A. tumefaciens* virB promoter and | TTGAGATCCTTTTTTTCTGCGCGTAATCTGCTGCTTGCAAACAAAAA[illegible] |
| pAGROm2_cas0 | module2_sgRNA entry_PJ23119-BsaI-[sfGFP-T3t placeholder]-BsaI-Scaf | carb100 | Module 2 entry plasmid for cloning a single sgRNA spacer into an expression cassette downstream from th | TTGAGATCCTTTTTTTCTGCGCGTAATCTGCTGCTTGCAAACAAAAA[illegible] |
| pAGROm2_cas1 | module2_sgRNA entry_PJ23100-BsaI-[sfGFP-T3t placeholder]-BsaI-Scaf | carb100 | Module 2 entry plasmid for cloning a single sgRNA spacer into an expression cassette downstream from th | TTGAGATCCTTTTTTTCTGCGCGTAATCTGCTGCTTGCAAACAAAAA[illegible] |
| pAGROm2_cas2 | module2_sgRNA entry_PJ23107-BsaI-[sfGFP-T3t placeholder]-BsaI-Scaf | carb100 | Module 2 entry plasmid for cloning a single sgRNA spacer into an expression cassette downstream from th | TTGAGATCCTTTTTTTCTGCGCGTAATCTGCTGCTTGCAAACAAAAA[illegible] |
| pAGROm2_int0 | module2_INTEGRATE crRNA entry_Vchrep-BsaI-[pint-sfGFP-T3T placeholder]-BsaI-Vchrep-T7T | carb100 | Module 2 entry plasmid for cloning crRNA spacers (singly or in an array of up to three) for targeting loci of | TTGAGATCCTTTTTTTCTGCGCGTAATCTGCTGCTTGCAAACAAAAA[illegible] |
| pAGROm3_cas0 | module3_sgRNA entry_PJ23119-BsaI-[sfGFP-T3t placeholder]-BsaI-Scaf | carb100 | Module 3 entry plasmid for cloning a single sgRNA spacer into an expression cassette downstream from th | TTGAGATCCTTTTTTTCTGCGCGTAATCTGCTGCTTGCAAACAAAAA[illegible] |
| pAGROm3_cas1 | module3_sgRNA entry_PJ23100-BsaI-[sfGFP-T3t placeholder]-BsaI-Scaf | carb100 | Module 3 entry plasmid for cloning a single sgRNA spacer into an expression cassette downstream from th | TTGAGATCCTTTTTTTCTGCGCGTAATCTGCTGCTTGCAAACAAAAA[illegible] |

| | | | | |
|---|---|---|---|---|
| pAGROm3_cas2 | module3_sgRNA entry_PJ23107-BsaI-[sfGFP-T3t placeholder]-BsaI-Scaf | carb100 | Module 3 entry plasmid for cloning a single sgRNA spacer into an expression cassette downstream from th | TTGAGATCCTTTT[illegible] |
| pAGROm3_int0 | module3_INTEGRATE cargo entry_VchINT R end-BbsI-[lacZa placeholder]-BbsI-VchINT L end | carb100, IPTG250, X-gal50 | Module 3 entry plasmid for cloning custom cargos for targeted insertion using the INTEGRATE system. A la | TTGAGATCCTTTT[illegible] |
| pAGROm1_cas0 | module1_pvirB-SpCas9-T3T | carb100 | Module 1 containing the wild-type *Streptococcus pyogenes* Cas9 nuclease under an *A. tumefaciens* virB (ac | TTGAGATCCTTTT[illegible] |
| pAGROm3_rec145 | module3_tonBt-TetR/$P_{Tet}$-pluγ-RecET_rhi145-rrnB_t1t2 | carb100 | Module 3 containing a recombineering gene cluster constristing of an anhydrotetracycline-inducible polyci | TTGAGATCCTTTT[illegible] |
| pAGROm3_rec483 | module3_tonBt-TetR/$P_{Tet}$-pluγ-RecETh_rhi483-rrnB_t1t2 | carb100 | Module 3 containing a recombineering gene cluster constristing of an anhydrotetracycline-inducible polyci | TTGAGATCCTTTT[illegible] |
| pAGROm3_recB6 | module3_tonBt-TetR/$P_{Tet}$-RecETh1h2h3h4_agrob6-rrnB_t1t2 | carb100 | Module 3 containing a recombineering gene cluster constristing of an anhydrotetracycline-inducible polyci | TTGAGATCCTTTT[illegible] |

| Gene fragment name | Sequence | Enzyme sites included | Purpose |
|---|---|---|---|
| PJ23100_Paq_GG_s1 | TTCAATACTTTCATATATCCAACACCTGCAAAACGTACCAACC | PaqCI | Promoter for assembly of multiple transcriptional units (used to drive pheS* expression in pAGROm4_220 and pAGROm4_221, and and codAD314A expression in pAGROm4_230, and pAGROm4_231) |
| PJ23119_Paq_GG_s1 | TTCAATACTTTCATATATCCAACACCTGCAAAACGTACCAACC | PaqCI | Promoter for assembly of multiple transcriptional units (used to drive sfGFP expression in pAGROm2_1XX, pAGROm2_casX, and pAGROm3_casX series) |
| Pint_Paq_GG_s1 | TTCAATACTTTCATATATCCAACACCTGCAAAACGTACCAACC | PaqCI | Promoter for assembly of multiple transcriptional units (used to drive eforRed expression in pAGROm4_2XX and pAGROex_2XX series, sfGFP in pAGROm2_int0, amilCP [pAGROm2_412, pAGROm4_410], aeBlue [pAGROm2_512, pAGROm4_510], and tsPurple [pAGROm2_612, pAGROm4_610]) |
| PAtu0824_Paq_GG_s1 | TTCAATACTTTCATATATCCAACACCTGCAAAACGTACCAACC | PaqCI | Promoter for assembly of multiple transcriptional units (used to drive recAC58 expression in pAGROm4_2X1 series |
| T3term_Paq_GG_s3 | GGGGCAAAGTGTCACCTGCAAAACCGCAATAAGCAAACCC | PaqCI | Terminator for assembly of multiple transcriptional units (used for sfGFP expression cassettes in pAGROm2_1XX, pAGROm2_casX, and pAGROm3_casX series) |
| T7term_Paq_GG_s3 | GGGGCAAAGTGTCACCTGCAAAACCGCATAGCATAACCCC | PaqCI | Terminator for assembly of multiple transcriptional units (used for eforRed expression cassettes in pAGROm4_2XX and pAGROex_2XX series, Cre expression in pAGROex_21X series, amilCP [pAGROm2_412, pAGROm4_410], aeBlue [pAGROm2_512, pAGROm4_510], and tsPurple [pAGROm2_612, pAGROm4_610]) |
| rrnB-T1term_Paq_GG_s3 | GGGGCAAAGTGTCACCTGCAAAACCGCCAAATAAAACGAAA | PaqCI | Terminator for assembly of multiple transcriptional units (used for sacB in pAGROm4_210 and pAGROm4_211, pheS* in pAGROm4_220 and pAGROm4_221, codAD314A in pAGROm4_230 and pAGROm4_231, and recAC58 in pAGROm4_2X1 series |
| Bluebonnet2_CDS_RhiLeg-co_Paq_s | TAGTCTTTCAGCACTCCTCAACCACCTGCAAAAATCATTTATC | PaqCI | Coding sequence for assembly of transcriptional unit ultimately used in pAGROm2_301, pAGROm2_311, pAGROm2_321, and pAGROm2_331 |
| amilCP_CDS_RhiLeg-co_Paq_s2 | TAGTCTTTCAGCACTCCTCAACCACCTGCAAAAATCATTTATC | PaqCI | Coding sequence for assembly of transcriptional unit ultimately used in pAGROm2_412 and pAGROm4_410 |
| tsPurple_CDS_RhiLeg-co_Paq_s2 | TAGTCTTTCAGCACTCCTCAACCACCTGCAAAAATCATTTATC | PaqCI | Coding sequence for assembly of transcriptional unit ultimately used in pAGROm2_612 and pAGROm4_610 |
| PN25::ApmA::T0term_RhiLeg-co_Paq_s1-3 (apraR) | ATTCAATACTTTAACACCTGCAAAACGTACACTGCTCAGACC | PaqCI | Whole transcriptional unit for apraR cassette used in pAGROm2_XX4, pAGRObb_X04, and pAGROex_2X4 series |
| pheS-T259A-A312G_CDS_Rrhi_LBA9402_RhiLeg-co_Paq_s2 | TAGTCTTTCAGCACTCCTCAACCACCTGCAAAAATCATTTATC | PaqCI | Coding sequence for assembly of transcriptional unit ultimately used in pAGROm4_220 and pAGROm4_221 |
| codA-D314A_CDS_RhiLeg-co_Paq_s2 | TAGTCTTTCAGCACTCCTCAACCACCTGCAAAAATCATTTATC | PaqCI | Coding sequence for assembly of transcriptional unit ultimately used in pAGROm4_230 and pAGROm4_231 |
| lacZa_MCS_del | GGGATTCAATACTTTCCAGCGCAACGCAATTAATGTGAGTTA | None | Used as PCR template for assembly into pAGROmX_000L entry module series, pAGROm2_1X2LC series, and pAGROm3_int0 |

| Primer name | Sequence | Template | Enzyme ends | Purpose |
|---|---|---|---|---|
| ccdB-CmR_amp-F | GTCCTTGCATGCACTTTTTGCAGGTGACCAC | pTRANS_220 | SphI | Amplifying a fragment containing the ccdB gene and a chloramphenicol resistance marker from pTRANS_220 for digestion with SphI and KpnI enzymes followed by ligation into same digested pEA186 backbone fragment, yielding pVS1 (S) |
| ccdB-CmR_amp-R | TTCGTACGACCCTCGGTA | pTRANS_220 | KpnI (within ter | Amplifying a fragment containing the ccdB gene and a chloramphenicol resistance marker from pTRANS_220 for digestion with SphI and KpnI enzymes followed by ligation into same digested pEA186 backbone fragment, yielding pVS1 (S) |
| ccdB-BsaI-del-frag1-F | GCTTGGTCTCTAAACCGGCACACTGGC | pVS1 (S) | BsaI | Amplifying a fragment from pVS1 (S) for removal of internal BsaI sites, re-assembly with two other fragments yields pVS-rec (S) via golden gate assembly using BsaI |
| ccdB-BsaI-del-frag1-R | GCTTGGTCTCGGGACCTACCAAGGCAACG | pVS1 (S) | BsaI | Amplifying a fragment from pVS1 (S) for removal of internal BsaI sites, re-assembly with two other fragments yields pVS-rec (S) via golden gate assembly using BsaI |
| ccdB-BsaI-del-frag2-F | GCTTGGTCTCTGTCCAGCGGCGGAG | pVS1 (S) | BsaI | Amplifying a fragment from pVS1 (S) for removal of internal BsaI sites, re-assembly with two other fragments yields pVS-rec (S) via golden gate assembly using BsaI |
| ccdB-BsaI-del-frag2-R | GCTTGGTCTCTCCATGCAGCGTACGAAGAAG | pVS1 (S) | BsaI | Amplifying a fragment from pVS1 (S) for removal of internal BsaI sites, re-assembly with two other fragments yields pVS-rec (S) via golden gate assembly using BsaI |
| ccdB-BsaI-del-frag3-F | GCTTGGTCTCTATGGCAACGTGCGTGG | pVS1 (S) | BsaI | Amplifying a fragment from pVS1 (S) for removal of internal BsaI sites, re-assembly with two other fragments yields pVS-rec (S) via golden gate assembly using BsaI |
| ccdB-BsaI-del-frag3-R | GCTTGGTCTCTGTTTCCGTTATCGGGGAAGAAGTG | pVS1 (S) | BsaI | Amplifying a fragment from pVS1 (S) for removal of internal BsaI sites, re-assembly with two other fragments yields pVS-rec (S) via golden gate assembly using BsaI |
| GG_sfGFP_SD_CDS_amp-F | CTGTCACCTGCGCTTATCAATTTATCTTCTACAAGGAGGTTAAGAA | pEN-L1-PJ23119-PplpT-sfGFP-T | PaqCI | Amplifying the sfGFP coding sequence with a Shine-Dalgarno sequence for assembling expression cassettes with synthetic promoter and terminator sequence fragments into pVS-rec (S) to make pVS-sfGFP-1, pVS-sfGFP-2, pVS-sfGFP-3, and pVS-sfGFP-4 via golden gate assembly using PaqCI |
| GG_sfGFP_CDS_amp-R | CTGTCACCTGCGCTTGCGGTCATTTGTACAGTTCATCCATACC | pEN-L1-PJ23119-PplpT-sfGFP-T | PaqCI | Amplifying the sfGFP coding sequence with a Shine-Dalgarno sequence for assembling expression cassettes with synthetic promoter and terminator sequence fragments into pVS-rec (S) to make pVS-sfGFP-1, pVS-sfGFP-2, pVS-sfGFP-3, and pVS-sfGFP-4 via golden gate assembly using PaqCI |
| GG_efRed_SD_CDS_amp-F | CTGTCACCTGCGCTTATCAATTTATCTTCTACAAGGAGGTTAAGAA | eforRed chromoprotein (Addgen | PaqCI | Amplifying the eforRed coding sequence with a Shine-Dalgarno sequence for assembling expression cassettes with synthetic promoter and terminator sequence fragments into pVS-rec (S) to make pVS-eforRed-1, pVS-eforRed-2, pVS-eforRed-3, and pVS-eforRed-4 via golden gate assembly using PaqCI |
| GG_efRed_CDS_amp-R | CTGTCACCTGCGCTTGCGGTTATGGGAGAGCCTTCGG | eforRed chromoprotein (Addgen | PaqCI | Amplifying the eforRed coding sequence with a Shine-Dalgarno sequence for assembling expression cassettes with synthetic promoter and terminator sequence fragments into pVS-rec (S) to make pVS-eforRed-1, pVS-eforRed-2, pVS-eforRed-3, and pVS-eforRed-4 via golden gate assembly using PaqCI |
| GG_aeBlue_SD_CDS_amp-F | CTGTCACCTGCGCTTATCAATTTATCTTCTACAAGGAGGTTAAGAA | aeBlue chromoprotein (Addgen | PaqCI | Amplifying the aeBlue coding sequence with a Shine-Dalgarno sequence for assembling expression cassettes with synthetic promoter and terminator sequence fragments into pVS-rec (S) to make pVS-aeBlue-1, pVS-aeBlue-2, pVS-aeBlued-3, and pVS-aeBlue-4 via golden gate assembly using PaqCI |
| GG_aeBlue_CDS_amp-R | CTGTCACCTGCGCTTGCGGTTAGTGATGCCCTAATTTACTCGG | aeBlue chromoprotein (Addgen | PaqCI | Amplifying the aeBlue coding sequence with a Shine-Dalgarno sequence for assembling expression cassettes with synthetic promoter and terminator sequence fragments into pVS-rec (S) to make pVS-aeBlue-1, pVS-aeBlue-2, pVS-aeBlued-3, and pVS-aeBlue-4 via golden gate assembly using PaqCI |
| sfGFP_cass_amp-F | CTGTCACCTGCGCTTCACTGAAAGTGAAACGTGATTTCATGCG | pEN-L1-PJ23119-PplpT-sfGFP-T | PaqCI | Amplifying the sfGFP whole expression cassette (which was non-functional in agrobacteria) from pEN-L1-PJ23119-PplpT-sfGFP-TrrfB-BsaI-Scaf-L2 for assembly into pVS-rec (S) to make pVS-sfGFP-0 via golden gate assembly using PaqCI |
| sfGFP_cass_amp-R | CTGTCACCTGCGCTTGATCTTCTATAAACGCAGAAAGGCCC | pEN-L1-PJ23119-PplpT-sfGFP-T | PaqCI | Amplifying the sfGFP whole expression cassette (which was non-functional in agrobacteria) from pEN-L1-PJ23119-PplpT-sfGFP-TrrfB-BsaI-Scaf-L2 for assembly into pVS-rec (S) to make pVS-sfGFP-0 via golden gate assembly using PaqCI |
| color_cass_amp-F | CTGTCACCTGCGCTTCACTCGCTTCTAGAGTTTACGGCTA | eforRed chromoprotein (Addgen | PaqCI | Amplifying the eforRed and aeBlue whole expression cassettes (which was non-functional in agrobacteria) from "eforRed chromoprotein" and "eforRed chromoprotein" plasmids, respectively for assembly into pVS-rec (S) to make pVS-eforRed-0 and pVS-aeBlue-0 via golden gate assembly using PaqCI |
| color_cass_amp-R | CTGTCACCTGCGCTTGATCCGCGAAGTAATCTTTTCGGTTTTAAA[illegible] | eforRed chromoprotein (Addgen | PaqCI | Amplifying the eforRed and aeBlue whole expression cassettes (which was non-functional in agrobacteria) from "eforRed chromoprotein" and "eforRed chromoprotein" plasmids, respectively for assembly into pVS-rec (S) to make pVS-eforRed-0 and pVS-aeBlue-0 via golden gate assembly using PaqCI |

| Primer name | Sequence | Template | Enzyme ends | Purpose |
|---|---|---|---|---|
| AeBlue-fxPq1-F | GTCTGGTCTCTCATCTGCGCACTACCTATCGTTC | aeBlue chromoprotein (Addgene | BsaI | Removal of a PaqCI site in the aeBlue coding sequence in "aeBlue chromoprotein," yielding "aeBlue chromoprotein-purged" via golden gate assembly using BsaI |
| AeBlue-fxPq1-hlpR | GTCTGGTCTCTATAGTGTTCACCCTTGTTACACC | aeBlue chromoprotein (Addgene | BsaI | Removal of a PaqCI site in the aeBlue coding sequence in "aeBlue chromoprotein," yielding "aeBlue chromoprotein-purged" via golden gate assembly using BsaI |
| AeBlue-fxPq1-hlpF | GTCTGGTCTCTCTATCCCATATCACCAGCTCAC | aeBlue chromoprotein (Addgene | BsaI | Removal of a PaqCI site in the aeBlue coding sequence in "aeBlue chromoprotein," yielding "aeBlue chromoprotein-purged" via golden gate assembly using BsaI |
| AeBlue-fxPq1-R | GTCTGGTCTCTGATGGGACGTCAGATGATTGC | aeBlue chromoprotein (Addgene | BsaI | Removal of a PaqCI site in the aeBlue coding sequence in "aeBlue chromoprotein," yielding "aeBlue chromoprotein-purged" via golden gate assembly using BsaI |
| GG_sfGFP_SapI_del-F | ATCAATTTATCTTCTACAAGGAGGTTAAGCATGCGTAAAGGCGAA | N/A (used for annealing with par | N/A | Removal of a SapI site in the sfGFP coding sequence by assembly with PJ23119 promoter and T3 terminator sequence fragments into pVS-rec (S), yielding pVS-sfGFP-2-purged via golden gate assembly using PaqCI |
| GG_sfGFP_SapI_del-R | CAGTTCTTCGCCTTTACGCATGCTTAACCTCCTTGTAGAAGATAAA | N/A (used for annealing with par | N/A | Removal of a SapI site in the sfGFP coding sequence by assembly with PJ23119 promoter and T3 terminator sequence fragments into pVS-rec (S), yielding pVS-sfGFP-2-purged via golden gate assembly using PaqCI |
| GG_sfGFP_trCDS_amp-F | CTGTCACCTGCGCTTACTGTTCACTGGTGTCGTCC | pVS-sfGFP-2 | PaqCI | Removal of a SapI site in the sfGFP coding sequence by assembly with PJ23119 promoter and T3 terminator sequence fragments into pVS-rec (S), yielding pVS-sfGFP-2-purged via golden gate assembly using PaqCI |
| GG_sfGFP_trCDS_amp-R | CTGTCACCTGCGCTTGCGGAAGCTTGGTCATTTGTACAGTTCATC | pVS-sfGFP-2 | PaqCI | Removal of a SapI site in the sfGFP coding sequence by assembly with PJ23119 promoter and T3 terminator sequence fragments into pVS-rec (S), yielding pVS-sfGFP-2-purged via golden gate assembly using PaqCI |
| GG_efRed_BbsI_del-F | ATCAATTTATCTTCTACAAGGAGGTTACATATGTCAGTGATTAAGCA | N/A (used for annealing with par | N/A | Removal of a BbsI site in the eforRed coding sequence by assembly with Pint promoter and T7 terminator sequence fragments into pVS-rec (S), yielding pVS-eforRed-3-purged via golden gate assembly using PaqCI |
| GG_efRed_BbsI_del-R | GGTTTTCATTACCTGCTTAATCACTGACATATGTAACCTCCTTGTAC | N/A (used for annealing with par | N/A | Removal of a BbsI site in the eforRed coding sequence by assembly with Pint promoter and T7 terminator sequence fragments into pVS-rec (S), yielding pVS-eforRed-3-purged via golden gate assembly using PaqCI |
| GG_efRed_trCDS_amp-F | CTGTCACCTGCGCTTAACCAAGTTGCACCTTGAGG | pVS-eforRed-3 | PaqCI | Removal of a BbsI site in the eforRed coding sequence by assembly with Pint promoter and T7 terminator sequence fragments into pVS-rec (S), yielding pVS-eforRed-3-purged via golden gate assembly using PaqCI |
| GG_efRed_trCDS_amp-R | CTGTCACCTGCGCTTGCGGCCTGCAGGTTTATGGGAGAGCCTTC | pVS-eforRed-3 | PaqCI | Removal of a BbsI site in the eforRed coding sequence by assembly with Pint promoter and T7 terminator sequence fragments into pVS-rec (S), yielding pVS-eforRed-3-purged via golden gate assembly using PaqCI |
| oriT_amp-F | GTCTGGTCTCTGTCACTGTGACTGGTGAGTATTCCA | pCGT4464A (provided by Roger | BsaI | Removal of a SapI site in the oriT sequence in pCGT4464A by re-assembly, yielding pCGT4464A-purged via golden gate assembly using BsaI |
| oriT_SapI_del_amp-R | GTCTGGTCTCTCTTGGTTCGTCTGGAAGGCAGTAC | pCGT4464A (provided by Roger | BsaI | Removal of a SapI site in the oriT sequence in pCGT4464A by re-assembly, yielding pCGT4464A-purged via golden gate assembly using BsaI |
| oriT_SapI_del_amp-F | GTCTGGTCTCTCAAGAGCGATTGAGGAAAAGGC | pCGT4464A (provided by Roger | BsaI | Removal of a SapI site in the oriT sequence in pCGT4464A by re-assembly, yielding pCGT4464A-purged via golden gate assembly using BsaI |
| oriT_amp-R | GTCTGGTCTCTGGCGACTTGGTTGAGTGATCCG | pCGT4464A (provided by Roger | BsaI | Removal of a SapI site in the oriT sequence in pCGT4464A by re-assembly, yielding pCGT4464A-purged via golden gate assembly using BsaI |
| GG_CrecA_CDS_amp-F | CTGTCACCTGCGCTTATCAATTTATCTTCTACAAGGAGGTTAACCA | C58 genomic DNA | PaqCI | Amplifying the native *A. tumefaciens* C58 *recA* coding sequence a Shine-Dalgarno sequence for assembling expression cassettes with synthetic promoter and terminator sequence fragments into pVS-rec (S) to make pVS-recA-4 via golden gate assembly using PaqCI |
| GG_CrecA_CDS_amp-R | CTGTCACCTGCGCTTGCGGCCTGCAGGAATCAGCCCTCGTCGC | C58 genomic DNA | PaqCI | Amplifying the native *A. tumefaciens* C58 *recA* coding sequence a Shine-Dalgarno sequence for assembling expression cassettes with synthetic promoter and terminator sequence fragments into pVS-rec (S) to make pVS-recA-4 via golden gate assembly using PaqCI |
| GG_C58recA_RE_del-0F | ATCAATTTATCTTCTACAAGGAGGTTAACCATGGCACAAAATTCTTT | N/A (used for annealing with par | N/A | Removal of Esp3I and BbsI sites in the native C58 recA coding sequence in pVS-recA-4 by assembly with PAtu0824 promoter and rrnB T1 terminator sequence fragments into pVS-rec (S), yielding pVS-recA-4-purged via golden gate assembly using PaqCI |
| GG_C58recA_RE_del-0R | GAGCCGCAAAGAATTTTGTGCCATGGTTAACCTCCTTGTAGAAGA | N/A (used for annealing with par | N/A | Removal of Esp3I and BbsI sites in the native C58 recA coding sequence in pVS-recA-4 by assembly with PAtu0824 promoter and rrnB T1 terminator sequence fragments into pVS-rec (S), yielding pVS-recA-4-purged via golden gate assembly using PaqCI |
| GG_C58recA_RE_del-1F | CTGTCACCTGCGCTTGCTCGTAGAGGATAAATCGGTGG | pVS-recA-4 | PaqCI | Removal of Esp3I and BbsI sites in the native C58 recA coding sequence in pVS-recA-4 by assembly with PAtu0824 promoter and rrnB T1 terminator sequence fragments into pVS-rec (S), yielding pVS-recA-4-purged via golden gate assembly using PaqCI |
| GG_C58recA_RE_del-1R | CTGTCACCTGCGCTTAAGGCCCGGCAGGCTATCG | pVS-recA-4 | PaqCI | Removal of Esp3I and BbsI sites in the native C58 recA coding sequence in pVS-recA-4 by assembly with PAtu0824 promoter and rrnB T1 terminator sequence fragments into pVS-rec (S), yielding pVS-recA-4-purged via golden gate assembly using PaqCI |
| GG_C58recA_RE_del-2F | CTGTCACCTGCGCTTCCTTCAGGCCCGTCTGATGAG | pVS-recA-4 | PaqCI | Removal of Esp3I and BbsI sites in the native C58 recA coding sequence in pVS-recA-4 by assembly with PAtu0824 promoter and rrnB T1 terminator sequence fragments into pVS-rec (S), yielding pVS-recA-4-purged via golden gate assembly using PaqCI |
| GG_C58recA_RE_del-2R | CTGTCACCTGCGCTTGCGGCGCGCCTCAGCCCTCGTCGCTG | pVS-recA-4 | PaqCI | Removal of Esp3I and BbsI sites in the native C58 recA coding sequence in pVS-recA-4 by assembly with PAtu0824 promoter and rrnB T1 terminator sequence fragments into pVS-rec (S), yielding pVS-recA-4-purged via golden gate assembly using PaqCI |
| 2299_RK2_bb-amp-p1F | GTCTGGTCTCTTCCTAGATCAAAGGATCTTCTTGAGATCCT | pKL2299A | BsaI | Removal of three PaqCI sites and an Esp3I site in the RK2 replicon from pKL2299A by assembly of 5 parts yielding pRK2 (S) via golden gate assembly using BsaI |
| 2299_RK2_bb-amp-p1R | GTCTGGTCTCTAGCCGACGCGAACGCTCC | pKL2299A | BsaI | Removal of three PaqCI sites and an Esp3I site in the RK2 replicon from pKL2299A by assembly of 5 parts yielding pRK2 (S) via golden gate assembly using BsaI |
| 2299_RK2_bb-amp-p2F | GTCTGGTCTCTGGCTCGAACAGGAGGCGG | pKL2299A | BsaI | Removal of three PaqCI sites and an Esp3I site in the RK2 replicon from pKL2299A by assembly of 5 parts yielding pRK2 (S) via golden gate assembly using BsaI |
| 2299_RK2_bb-amp-p2R | GTCTGGTCTCTCTTGCTGCCACACCAGTTC | pKL2299A | BsaI | Removal of three PaqCI sites and an Esp3I site in the RK2 replicon from pKL2299A by assembly of 5 parts yielding pRK2 (S) via golden gate assembly using BsaI |
| 2299_RK2_bb-amp-p3F | GTCTGGTCTCTCAAGTGTTGGAGTACGCGAAGC | pKL2299A | BsaI | Removal of three PaqCI sites and an Esp3I site in the RK2 replicon from pKL2299A by assembly of 5 parts yielding pRK2 (S) via golden gate assembly using BsaI |
| 2299_RK2_bb-amp-p3R | GTCTGGTCTCTGAGGTGGCTGCTGAACC | pKL2299A | BsaI | Removal of three PaqCI sites and an Esp3I site in the RK2 replicon from pKL2299A by assembly of 5 parts yielding pRK2 (S) via golden gate assembly using BsaI |
| 2299_RK2_bb-amp-p4F | GTCTGGTCTCTCCTCCATCGCCCTAGGTCCC | pKL2299A | BsaI | Removal of three PaqCI sites and an Esp3I site in the RK2 replicon from pKL2299A by assembly of 5 parts yielding pRK2 (S) via golden gate assembly using BsaI |
| 2299_RK2_bb-amp-p4R | GTCTGGTCTCTTACGCGAATGCGCGAGGTCTAT | pKL2299A | BsaI | Removal of three PaqCI sites and an Esp3I site in the RK2 replicon from pKL2299A by assembly of 5 parts yielding pRK2 (S) via golden gate assembly using BsaI |
| specR_new_amp-F | GTCTGGTCTCTCGTATCCTGCTTTTGATCGGTACCA | pVS-rec (S) | BsaI | Removal of three PaqCI sites and an Esp3I site in the RK2 replicon from pKL2299A by assembly of 5 parts yielding pRK2 (S) via golden gate assembly using BsaI |
| specR_new_amp-R | GTCTGGTCTCTAGGAGTAGGGCTTATTATGCACGCTT | pVS-rec (S) | BsaI | Removal of three PaqCI sites and an Esp3I site in the RK2 replicon from pKL2299A by assembly of 5 parts yielding pRK2 (S) via golden gate assembly using BsaI |
| gentR_start-F | GTCTGGTCTCTTTCGCACCGTGGAAACGGATGAA | pKL2299A (Addgene #: 222105) | BsaI | Removal of Esp3I site in the aacC1 gentamicin resistance cassette from pKL2299A by assembly into pRK2 (S) backbone, yielding pRK2 (Ge) via golden gate assembly using BsaI |
| gentR_Esp3I_amp-R | GTCTGGTCTCTCACCGAACTCACGACCGAAA | pKL2299A (Addgene #: 222105) | BsaI | Removal of Esp3I site in the aacC1 gentamicin resistance cassette from pKL2299A by assembly into pRK2 (S) backbone, yielding pRK2 (Ge) via golden gate assembly using BsaI |
| gentR_Esp3I_amp-F | GTCTGGTCTCTGGTGACGTAGCCACCTACTCC | pKL2299A (Addgene #: 222105) | BsaI | Removal of Esp3I site in the aacC1 gentamicin resistance cassette from pKL2299A by assembly into pRK2 (S) backbone, yielding pRK2 (Ge) via golden gate assembly using BsaI |
| gentR_end-R | GTCTGGTCTCTAGGACGTTGTGACAATTTACCGAACAAC | pKL2299A (Addgene #: 222105) | BsaI | Removal of Esp3I site in the aacC1 gentamicin resistance cassette from pKL2299A by assembly into pRK2 (S) backbone, yielding pRK2 (Ge) via golden gate assembly using BsaI |
| Tyr-rec_cass-v2F | CTGTCACCTGCGCTTCACTCCTTTACGGCTAGCTCAGC | pKL2315 (Addgene #: 187876) | PaqCI | Amplifying the Cre expression cassette in pKL2315 for assembly into pVS-rec (S) to make pVS-Cre-A via golden gate assembly using PaqCI |
| Tyr-rec_cass-v2R | CTGTCACCTGCGCTTGATCTAAGCAGAAAGTCAAAAGCCTCC | pKL2315 (Addgene #: 187876) | PaqCI | Amplifying the Cre expression cassette in pKL2315 for assembly into pVS-rec (S) to make pVS-Cre-A via golden gate assembly using PaqCI |
| Cre-cass_tswap-F | CTGTCACCTGCGCTTCGTACTTTACGGCTAGCTCAGCC | pVS-Cre-A | PaqCI | Amplifying the Cre coding sequence and PJ23107 promoter in pVS-Cre-A for assembly with the T7 termintor synthetic fragment into pVS-rec (S) to make pVS-Cre-A-purged via golden gate assembly using PaqCI, removing a BsaI site to the 3' of the coding sequence in original construct |
| Cre-cass_tswap-R | CTGTCACCTGCGCTTGCGGTTCAGTCACCATCTTCCAGG | pVS-Cre-A | PaqCI | Amplifying the Cre coding sequence and PJ23107 promoter in pVS-Cre-A for assembly with the T7 termintor synthetic fragment into pVS-rec (S) to make pVS-Cre-A-purged via golden gate assembly using PaqCI, removing a BsaI site to the 3' of the coding sequence in original construct |
| chlorR_amp-1F | GTCTGGTCTCTGTCATGACGTTGATCGGCACGTAA | pCGT4464A (provided by Roger | BsaI | Removal of Esp3I site in the CAT chloramphenicol resistance cassette from pCGT4464A by assembly into pRK2 (S) backbone, yielding pRK2 (Cm) via golden gate assembly using BsaI |
| chlorR_amp-1R | CTGTGGTCTCTACACGAAAAACATATTCTCAATAAACCCTTTAGG | pCGT4464A (provided by Roger | BsaI | Removal of Esp3I site in the CAT chloramphenicol resistance cassette from pCGT4464A by assembly into pRK2 (S) backbone, yielding pRK2 (Cm) via golden gate assembly using BsaI |
| chlorR_amp-2F | CTGTGGTCTCTGTGTCAGCCAATCCCTGGG | pCGT4464A (provided by Roger | BsaI | Removal of Esp3I site in the CAT chloramphenicol resistance cassette from pCGT4464A by assembly into pRK2 (S) backbone, yielding pRK2 (Cm) via golden gate assembly using BsaI |

| | | | | |
|---|---|---|---|---|
| chlorR_amp-2R | GTCTGGTCTCTTAGTCTGCCACTCATCGCAGTA | pCGT4464A (provided by Roger ¯ | BsaI | Removal of Esp3I site in the CAT chloramphenicol resistance cassette from pCGT4464A by assembly into pRK2 (S) backbone, yielding pRK2 (Cm) via golden gate assembly using BsaI |

| Primer name | Sequence | Template | Enzyme ends | Purpose |
|---|---|---|---|---|
| oriT-fixed_amp-F | GTCTGGTCTCTGTCACTGTGACTGGTGAGTATTCCA | pCGT4464A-purged | BsaI | Amplifying the $oriT_{pRK2}$ sequence from pCGT4464A-purged for assembly of pDisarm_TiIa_v[1-6] via golden gate assembly using BsaI |
| oriT-fixed_amp-R | GTCTGGTCTCTCTGACACTTGGTTGAGTGATCC | pCGT4464A-purged | BsaI | Amplifying the $oriT_{pRK2}$ sequence from pCGT4464A-purged for assembly of pDisarm_TiIa_v[1-6] via golden gate assembly using BsaI |
| carbR-colE1_amp-F | GTCTGGTCTCGTCAGGTGGCACTTTTCG | pMOD_B2303 | BsaI | Amplifying the carbencillin resistance marker and colE1 ori sequence fragment from pMOD_B2303 for assembly of pDisarm_TiIa_v[1-6] via golden gate assembly using BsaI |
| carbR-colE1_amp-R | GTCTGGTCTCTGGCGTTCTGGGCGCTCTCCC | pMOD_B2303 | BsaI | Amplifying the carbencillin resistance marker and colE1 ori sequence fragment from pMOD_B2303 for assembly of pDisarm_TiIa_v[1-6] via golden gate assembly using BsaI |
| pTi_Type_Ia_L_HA-F | GTCTGGTCTCTCGCCCAGAGCTTGTCAGCTACC | Strain X1/95 genomic DNA | BsaI | Amplifying a left homology arm fragment from a Type Ia Ti plasmid for assembly of pDisarm_TiIa_v[1-6] via golden gate assembly using BsaI |
| pTi_Type_Ia_L_HA-R | GTCTGGTCTCTGCATGACTTTGCTCATGTTACCGATG | Strain X1/95 genomic DNA | BsaI | Amplifying a left homology arm fragment from a Type Ia Ti plasmid for assembly of pDisarm_TiIa_v[1-6] via golden gate assembly using BsaI |
| pTi_Type_Ia_R_HA-F | GTCTGGTCTCTTCCTGTAAGAGCGTGAAAGACTCATTC | Strain X1/95 genomic DNA | BsaI | Amplifying a right homology arm fragment from a Type Ia Ti plasmid for assembly of pDisarm_TiIa_v[1-6] via golden gate assembly using BsaI |
| pTi_Type_Ia_R_HA-R | GTCTGGTCTCTCGGTCGCAATTGCCAACCTCAAC | Strain X1/95 genomic DNA | BsaI | Amplifying a right homology arm fragment from a Type Ia Ti plasmid for assembly of pDisarm_TiIa_v[1-6] via golden gate assembly using BsaI |
| FRT_sfGFP-F | GTCTGGTCTCTATGCGGCGCGCCGAAGTTCCTATTCTCTAGAAA | pVS-sfGFP-2-purged | BsaI | Amplifying the sfGFP expression cassette from pVS-sfGFP-2-purged for assembly of pDisarm_TiIa_v1 and pDisarm_TiIa_v5 via golden gate assembly using BsaI |
| LoxP_sfGFP-F | GTCTGGTCTCTATGCGGCGCGCCATAACTTCGTATAATGTATGCT | pVS-sfGFP-2-purged | BsaI | Amplifying the sfGFP expression cassette from pVS-sfGFP-2-purged for assembly of pDisarm_TiIa_v2 and pDisarm_TiIa_v6 via golden gate assembly using BsaI |
| Lox2272_sfGFP-F | GTCTGGTCTCTATGCGGCGCGCCATAACTTCGTATAAAGTATCCT | pVS-sfGFP-2-purged | BsaI | Amplifying the sfGFP expression cassette from pVS-sfGFP-2-purged for assembly of pDisarm_TiIa_v3 via golden gate assembly using BsaI |
| Lox5171_sfGFP-F | GTCTGGTCTCTATGCGGCGCGCCATAACTTCGTATAATGTGTACT | pVS-sfGFP-2-purged | BsaI | Amplifying the sfGFP expression cassette from pVS-sfGFP-2-purged for assembly of pDisarm_TiIa_v4 via golden gate assembly using BsaI |
| sfGFP_amp1-R | GTCTGGTCTCTCGAATGAGCTCGTCCACATTGCGTAGACTGT | pVS-sfGFP-2-purged | BsaI | Amplifying the sfGFP expression cassette from pVS-sfGFP-2-purged for assembly of pDisarm_TiIa_v[1-6] via golden gate assembly using BsaI |
| kanR_amp1-F | GTCTGGTCTCTTTCGCGGCCGCCTTGTGTCTCAAAATCTCTGATG | pMIN-VS1 | BsaI | Amplifying the kanamycin resistance marker from pMIN-VS1 for assembly of pDisarm_TiIa_v[1-4] via golden gate assembly using BsaI |
| FRT_kanR_amp-R | GTCTGGTCTCTAGGACCCGGGGAAGTTCCTATACTTTCTAGAGA | pMIN-VS1 | BsaI | Amplifying the kanamycin resistance marker from pMIN-VS1 for assembly of pDisarm_TiIa_v1 via golden gate assembly using BsaI |
| LoxP_kanR_amp-R | GTCTGGTCTCTAGGACCCGGGATAACTTCGTATAGCATACATTAT | pMIN-VS1 | BsaI | Amplifying the kanamycin resistance marker from pMIN-VS1 for assembly of pDisarm_TiIa_v2 via golden gate assembly using BsaI |
| Lox2272_kanR_amp-R | GTCTGGTCTCTAGGACCCGGGATAACTTCGTATAGGATACTTTAT | pMIN-VS1 | BsaI | Amplifying the kanamycin resistance marker from pMIN-VS1 for assembly of pDisarm_TiIa_v3 via golden gate assembly using BsaI |
| Lox5171_kanR_amp-R | GTCTGGTCTCTAGGACCCGGGATAACTTCGTATAGTACACATTAT | pMIN-VS1 | BsaI | Amplifying the kanamycin resistance marker from pMIN-VS1 for assembly of pDisarm_TiIa_v4 via golden gate assembly using BsaI |
| specR_amp1-F | GTCTGGTCTCTTTCGCGGCCGCCTGCTTTTGATCGGTACCA | pVS-rec (S) | BsaI | Amplifying the spectinomycin resistance marker from pVS-rec (S) for assembly of pDisarm_TiIa_v5 and pDisarm_TiIa_v6 via golden gate assembly using BsaI |
| FRT_specR_amp-R | GTCTGGTCTCTAGGACCCGGGGAAGTTCCTATACTTTCTAGAGA | pVS-rec (S) | BsaI | Amplifying the spectinomycin resistance marker from pVS-rec (S) for assembly of pDisarm_TiIa_v5 via golden gate assembly using BsaI |
| LoxP_specR_amp-R | GTCTGGTCTCTAGGACCCGGGATAACTTCGTATAGCATACATTAT | pVS-rec (S) | BsaI | Amplifying the spectinomycin resistance marker from pVS-rec (S) for assembly of pDisarm_TiIa_v6 via golden gate assembly using BsaI |
| eforRed_cass_amp-F | GTCTGGTCTCTACCGGCGCGCCGTCGACCCAACCGAAAGGTG | pVS-eforRed-3-purged | BsaI | Amplifying the eforRed expression cassette from pVS-eforRed-3-purged for assembly of pDisarm_TiIa_v[1-6] via golden gate assembly using BsaI |
| eforRed_cass_amp-R | GTCTGGTCTCTTCGCGAGCTCGTCCACATTGCGTAGACTGT | pVS-eforRed-3-purged | BsaI | Amplifying the eforRed expression cassette from pVS-eforRed-3-purged for assembly of pDisarm_TiIa_v[1-6] via golden gate assembly using BsaI |
| sacB_amp-F | GTCTGGTCTCTGCGACACATATACCTGCCGTTCACT | pFlp-Km | BsaI | Amplifying the sacB counterselection marker from pFlp-Km for assembly of pDisarm_TiIa_v1 and pDisarm_TiIa_v2 via golden gate assembly using BsaI |
| sacB_amp-endR | GTCTGGTCTCTTGACGGAAGTCCATCGGTAACAGTCCCGGGGA | pFlp-Km | BsaI | Amplifying the sacB counterselection marker from pFlp-Km for assembly of pDisarm_TiIa_v1 via golden gate assembly using BsaI |
| sacB_amp-altR | GTCTGGTCTCTATCTGCGGCCGCCTTATTTGTTAACTGTTAATTGT | pFlp-Km | BsaI | Amplifying the sacB counterselection marker from pFlp-Km for assembly of pDisarm_TiIa_v2 via golden gate assembly using BsaI |
| pheS/codA_cass_amp-F | GTCTGGTCTCTGCGAGCCAACCGAAAGGTGACTCTA | pVS-pheS-1; pVS-codA-1 | BsaI | Amplifying the pheS* counterselection marker expression cassette from pVS-pheS-1 for assembly of pDisarm_TiIa_v3 and pDisarm_TiIa_v4 via golden gate assembly using BsaI; amplifying the $codA^{D314A}$ counterselection marker expression cassette from pVS-codA-1 for assembly of pDisarm_TiIa_v5 and pDisarm_TiIa_v6 via golden gate assembly using BsaI |
| pheS/codA_cass_amp-endR | GTCTGGTCTCTTGACGGAAGTCCATCGGTAACAGTCCCGGGGA | pVS-pheS-1; pVS-codA-1 | BsaI | Amplifying the pheS* counterselection marker expression cassette from pVS-pheS-1 for assembly of pDisarm_TiIa_v3 via golden gate assembly using BsaI; amplifying the $codA^{D314A}$ counterselection marker expression cassette from pVS-codA-1 for assembly of pDisarm_TiIa_v5 via golden gate assembly using BsaI |
| pheS/codA_cass_amp-altR | GTCTGGTCTCTATCTGCGGCCGCGTCCACATTGCGTAGACTGT | pVS-pheS-1; pVS-codA-1 | BsaI | Amplifying the pheS* counterselection marker expression cassette from pVS-pheS-1 for assembly of pDisarm_TiIa_v4 via golden gate assembly using BsaI; amplifying the $codA^{D314A}$ counterselection marker expression cassette from pVS-codA-1 for assembly of pDisarm_TiIa_v6 via golden gate assembly using BsaI |
| C58recA_cass_amp-F | GTCTGGTCTCTAGATCCAACCGAAAGGTGACTCTA | pVS-recA-4-purged | BsaI | Amplifying the $recA_{C58}$ expression cassette from pVS-recA-4-purged for assembly of pDisarm_TiIa_v2, pDisarm_TiIa_v4, and pDisarm_TiIa_v6 via golden gate assembly using BsaI |
| C58recA_cass_amp-R | GTCTGGTCTCTTGACGGAAGTCCATCGGTAACAGTCCCGGGGA | pVS-recA-4-purged | BsaI | Amplifying the $recA_{C58}$ expression cassette from pVS-recA-4-purged for assembly of pDisarm_TiIa_v2, pDisarm_TiIa_v4, and pDisarm_TiIa_v6 via golden gate assembly using BsaI |
| mod1_BB_amp-F | GTCTGGTCTCTCCGCATCAAAAAGCAGGTGTGTCAGGTGGCACT | pMOD_B2303 | BsaI | Amplifying the carbencillin resistance marker and colE1 ori sequence fragment from pMOD_B2303, installing PaqCI sites and overhangs for module 1 to construct pAGROm1_000L and pAGROm1_cbe0, via golden gate assembly using BsaI |
| mod1_BB_amp-R | GTCTGGTCTCTCGCCTACGTTTTGCAGGTGTGGCGCTCTCCCGC | pMOD_B2303 | BsaI | Amplifying the carbencillin resistance marker and colE1 ori sequence fragment from pMOD_B2303, installing PaqCI sites and overhangs for module 1 to construct pAGROm1_000L and pAGROm1_cbe0, via golden gate assembly using BsaI |
| mod2_BB_amp-F | GTCTGGTCTCTCCGCCCGCAAAAGCAGGTGTGTCAGGTGGCAC | pMOD_B2303 | BsaI | Amplifying the carbencillin resistance marker and colE1 ori sequence fragment from pMOD_B2303, installing PaqCI sites and overhangs for module 2 to construct pAGROm2_000L and remaining pAGROm2_[XXX] series via golden gate assembly using BsaI |
| mod2_BB_amp-R | GTCTGGTCTCTCGCCTGATTTTTGCAGGTGTGGCGCTCTCCCGC | pMOD_B2303 | BsaI | Amplifying the carbencillin resistance marker and colE1 ori sequence fragment from pMOD_B2303, installing PaqCI sites and overhangs for module 2 to construct pAGROm2_000L and remaining pAGROm2_[XXX] series via golden gate assembly using BsaI |
| mod2_cargo_amp-F | GTCTGGTCTCTATCACATCGGTAACATGAGCAAAGTC | pDisarm_TiIa_v[1-6] | BsaI | Amplifying replacement sequence fragments from pDisarm_TiIa_v[1-6] series to assemble with the mod2_BB_amp fragment, yielding pAGROm2_101, pAGROm2_111, pAGROm2_121, pAGROm2_131, pAGROm2_102, and pAGROm2_112, respectively, via golden gate assembly using BsaI |
| mod2_cargo_amp-R | GTCTGGTCTCTGCGGCGAATGAGTCTTTCACGCTCTTAC | pDisarm_TiIa_v[1-6] | BsaI | Amplifying replacement sequence fragments from pDisarm_TiIa_v[1-6] series to assemble with the mod2_BB_amp fragment, yielding pAGROm2_101, pAGROm2_111, pAGROm2_121, pAGROm2_131, pAGROm2_102, and pAGROm2_112, respectively, via golden gate assembly using BsaI |
| mod3_BB_amp-F | GTCTGGTCTCTCCGCGGGAAAAAGCAGGTGTGTCAGGTGGCAC | pMOD_B2303 | BsaI | Amplifying the carbencillin resistance marker and colE1 ori sequence fragment from pMOD_B2303, installing PaqCI sites and overhangs for module 3 to construct pAGROm3_000L via golden gate assembly using BsaI |
| mod3_BB_amp-R | GTCTGGTCTCTCGCCGCGGTTTTGCAGGTGTGGCGCTCTCCCG | pMOD_B2303 | BsaI | Amplifying the carbencillin resistance marker and colE1 ori sequence fragment from pMOD_B2303, installing PaqCI sites and overhangs for module 3 to construct pAGROm3_000L via golden gate assembly using BsaI |
| mod4_BB_amp-F | GTCTGGTCTCTCCGCTCAAAAAAGCAGGTGTGTCAGGTGGCACT | pMOD_B2303 | BsaI | Amplifying the carbencillin resistance marker and colE1 ori sequence fragment from pMOD_B2303, installing PaqCI sites and overhangs for module 4 to construct pAGROm4_000L and remaining pAGROm4_[XXX] series via golden gate assembly using BsaI |
| mod4_BB_amp-R | GTCTGGTCTCTCGCCTCCCTTTTGCAGGTGTGGCGCTCTCCCGC | pMOD_B2303 | BsaI | Amplifying the carbencillin resistance marker and colE1 ori sequence fragment from pMOD_B2303, installing PaqCI sites and overhangs for module 4 to construct pAGROm4_000L and remaining pAGROm4_[XXX] series via golden gate assembly using BsaI |
| mod4_cargo_amp-F | GTCTGGTCTCTGGGATTGAGGTTGGCAATTGCGA | pDisarm_TiIa_v[1-6] | BsaI | Amplifying replacement sequence fragments from pDisarm_TiIa_v[1-6] series to assemble with the mod4_BB_amp fragment, yielding pAGROm4_210, pAGROm4_211, pAGROm4_220, pAGROm4_221, pAGROm4_230, and pAGROm4_231, respectively, via golden gate assembly using BsaI |
| mod4_cargo_amp-R | GTCTGGTCTCTTTGAGGAAGTCCATCGGTAACAGT | pDisarm_TiIa_v[1-6] | BsaI | Amplifying replacement sequence fragments from pDisarm_TiIa_v[1-6] series to assemble with the mod4_BB_amp fragment, yielding pAGROm4_210, pAGROm4_211, pAGROm4_220, pAGROm4_221, pAGROm4_230, and pAGROm4_231, respectively, via golden gate assembly using BsaI |
| lacZa_amp1-F | GTCTGGTCTCTGGCGCGCCTTCCTGCAGGCGTCGACTAGCGCA | lacZa_MCS_del (synthetic fragm | BsaI | Amplifying the lacZα fragment from lacZa_MCS_del to assemble with mod[X]_BB_amp fragments and ccdB_amp1 fragment, yielding pAGROm[1-4]_000L via golden gate assembly using BsaI |
| lacZa_amp1-R | GTCTGGTCTCTACGATGGACTCCAACGTCAAAGG | lacZa_MCS_del (synthetic fragm | BsaI | Amplifying the lacZα fragment from lacZa_MCS_del to assemble with mod[X]_BB_amp fragments and ccdB_amp1 fragment, yielding pAGROm[1-4]_000L via golden gate assembly using BsaI |

| | | | | |
|---|---|---|---|---|
| ccdB_amp1-F | GTCTGGTCTCTTCGTAAGCCAGATAACAGTATGCGTATTT | pVS-rec (S) | BsaI | Amplifying the ccdB fragment from pVS-rec (S) to assemble with mod[X]_BB_amp fragments and lacZa_amp1 fragment, yielding pAGROm[1-4]_000L via golden gate assembly using BsaI |
| ccdB_amp1-R | GTCTGGTCTCTGCGGCCTTAACCCGGGTGACGTCAGCGGCCGC | pVS-rec (S) | BsaI | Amplifying the ccdB fragment from pVS-rec (S) to assemble with mod[X]_BB_amp fragments and lacZa_amp1 fragment, yielding pAGROm[1-4]_000L via golden gate assembly using BsaI |
| mod_000_oligo-F | CGCGCCTTCCTGCAGGCGTCGACAGAGCTCAGCGGCCGCTTG | N/A (used for annealing with par | N/A | Ligating annealed dsOligo pair consisting of an MCS linker sequence into AscI and XmaI-digested pAGROm[X]_000L backbone fragments, yielding pAGROm1_000, pAGROm2_000, pAGROm3_000, and pAGROm4_000 base module plasmids |
| mod_000_oligo-R | CCGGTTAACCCGGGTGACGTCAAGCGGCCGCTGAGCTCTGTCG | N/A (used for annealing with par | N/A | Ligating annealed dsOligo pair consisting of an MCS linker sequence into AscI and XmaI-digested pAGROm[X]_000L backbone fragments, yielding pAGROm1_000, pAGROm2_000, pAGROm3_000, and pAGROm4_000 base module plasmids |
| specR_NotI_amp-F | GGATTTAACTGCGGCCGCCTGCTTTTGATCGGTACCA | pVS-rec (S) | NotI | Amplifying the spectinomycin resistance marker from pVS-rec (S) for digestion with NotI and AatII enzymes followed by ligation into same digested pAGROm2_121 and pAGROm2_131 backbone fragments, yielding pAGROm2_122 and pAGROm2_132, respectively |
| specR_AatII_amp-R | GGATTTGGATGACGTCGTAGGGCTTATTATGCACGCTT | pVS-rec (S) | AatII | Amplifying the spectinomycin resistance marker from pVS-rec (S) for digestion with NotI and AatII enzymes followed by ligation into same digested pAGROm2_121 and pAGROm2_131 backbone fragments, yielding pAGROm2_122 and pAGROm2_132, respectively |
| gentR_NotI_amp-F | GGATTTAACTGCGGCCGCTCACCGTGGAAACGGATGAA | pRK2 (Ge) | NotI | Amplifying the gentamicin resistance marker from pRK2 (Ge) for digestion with NotI and AatII enzymes followed by ligation into same digested pAGROm2_101, pAGROm2_111, pAGROm2_121, and pAGROm2_131 backbone fragments, yielding pAGROm2_103, pAGROm2_113, pAGROm2_123 and pAGROm2_133, respectively |
| gentR_AatII_amp-R | GGATTTGGATGACGTCCGTTGTGACAATTTACCGAACAAC | pRK2 (Ge) | AatII | Amplifying the gentamicin resistance marker from pRK2 (Ge) for digestion with NotI and AatII enzymes followed by ligation into same digested pAGROm2_101, pAGROm2_111, pAGROm2_121, and pAGROm2_131 backbone fragments, yielding pAGROm2_103, pAGROm2_113, pAGROm2_123 and pAGROm2_133, respectively |
| apraR_NotI_amp-F | GGATTTAACTGCGGCCGCTCACTGCTCAGACCAAGGATA | PN25::ApmA::T0term_RhiLeg-co | NotI | Amplifying the apramycin resistance marker cassette from PN25::ApmA::T0term_RhiLeg-co_Paq_s1-3 for digestion with NotI and AatII enzymes followed by ligation into same digested pAGROm2_101, pAGROm2_111, pAGROm2_121, and pAGROm2_131 backbone fragments, yielding pAGROm2_104, pAGROm2_114, pAGROm2_124 and pAGROm2_134, respectively |
| apraR_AatII_amp-R | GGATTTGGATGACGTCGGACAGTTTGCTGAGATTGC | PN25::ApmA::T0term_RhiLeg-co | AatII | Amplifying the apramycin resistance marker cassette from PN25::ApmA::T0term_RhiLeg-co_Paq_s1-3 for digestion with NotI and AatII enzymes followed by ligation into same digested pAGROm2_101, pAGROm2_111, pAGROm2_121, and pAGROm2_131 backbone fragments, yielding pAGROm2_104, pAGROm2_114, pAGROm2_124 and pAGROm2_134, respectively |
| GG_pro_Paq_Sal_amp-F | CTGTCACCTGCAAGTCGACCCAACCGAAAGGTGACTCTA | pVS-BFP-3; pVS-amilCP-3; pVS-a | PaqCI/SalI | Amplifying the BFP expression cassette from pVS-BFP-3 for digestion with SalI and SacI enzymes followed by ligation into same digested pAGROm2_101, pAGROm2_111, pAGROm2_121, and pAGROm2_131 backbone fragments, yielding pAGROm2_301, pAGROm2_311, pAGROm2_321 and pAGROm2_331, respectively; amplifying the amilCP, aeBlue, and tsPurple expression cassettes from pVS-amilCP-3, pVS-aeBlue-3, and pVS-tsPurple-3 for digestion with SalI and SacI enzymes followed by ligation of each combinatorially into same digested pAGROm2_112 and pAGROm4_210 backbone fragments, yielding pAGROm2_412, pAGROm2_512, pAGROm2_612, pAGROm4_410, pAGROm4_510, and pAGROm4_610, respectively |
| GG_ter_Paq_Sac_amp-R | CTGTCACCTGCGAGCTCGTCCACATTGCGTAGACTGT | pVS-BFP-3; pVS-amilCP-3; pVS-a | PaqCI/SacI | Amplifying the BFP expression cassette from pVS-BFP-3 for digestion with SalI and SacI enzymes followed by ligation into same digested pAGROm2_101, pAGROm2_111, pAGROm2_121, and pAGROm2_131 backbone fragments, yielding pAGROm2_301, pAGROm2_311, pAGROm2_321 and pAGROm2_331, respectively; amplifying the amilCP, aeBlue, and tsPurple expression cassettes from pVS-amilCP-3, pVS-aeBlue-3, and pVS-tsPurple-3 for digestion with SalI and SacI enzymes followed by ligation of each combinatorially into same digested pAGROm2_112 and pAGROm4_210 backbone fragments, yielding pAGROm2_412, pAGROm2_512, pAGROm2_612, pAGROm4_410, pAGROm4_510, and pAGROm4_610, respectively |
| CXN_amp-1F | GTCTGGTCTCTCGCCCGTAAAAAGCAGGTGCAGTGGTATGGGGT | pAGM4673 | BsaI | Amplifying an upstream segment of the operon encoding the canthaxanthin biosynthetic pathway from pAGM4673 (removing an internal PaqCI site) for assembly of all pAGRObb_[X0X] via golden gate assembly using BsaI |
| CXN_amp-1R | CTGTGGTCTCTCAGATGGTGTTCGTGATGATAACCG | pAGM4673 | BsaI | Amplifying an upstream segment of the operon encoding the canthaxanthin biosynthetic pathway from pAGM4673 (removing an internal PaqCI site) for assembly of all pAGRObb_[X0X] via golden gate assembly using BsaI |
| CXN_amp-2F | CTGTGGTCTCTTCTGCACCCGACGGT | pAGM4673 | BsaI | Amplifying a downstream segment of the operon encoding the canthaxanthin biosynthetic pathway from pAGM4673 (removing an internal PaqCI site) for assembly of all pAGRObb_[X0X] via golden gate assembly using BsaI |
| CXN_amp-2R | GTCTGGTCTCTATCTTTGATTTTGCAGGTGTTCTTCCACTTGAGTGC | pAGM4673 | BsaI | Amplifying a downstream segment of the operon encoding the canthaxanthin biosynthetic pathway from pAGM4673 (removing an internal PaqCI site) for assembly of all pAGRObb_[X0X] via golden gate assembly using BsaI |
| oriT_amp-F | GTCTGGTCTCTAGATGCTGTGACTGGTGAGTATTCCA | pCGT4464A-purged | BsaI | Amplifying the $oriT_{pRK2}$ fragment from pCGT4464A-purged for assembly of all pAGRObb_[X0X] via golden gate assembly using BsaI |
| oriT_amp-v3R | GTCTGGTCTCTCGAAACTTGGTTGAGTGATCCGG | pCGT4464A-purged | BsaI | Amplifying the $oriT_{pRK2}$ fragment from pCGT4464A-purged for assembly of all pAGRObb_[X0X] via golden gate assembly using BsaI |
| chlorR_amp0-F | GTCTGGTCTCTTTCGGACGTTGATCGGCACGTAA | pRK2 (Cm) | BsaI | Amplifying chloramphenicol resistance marker from pRK2 (Cm) for assembly of pAGRObb_000 via golden gate assembly using BsaI |
| chlorR_amp0-R | GTCTGGTCTCTAGGACTGCCACTCATCGCAGTA | pRK2 (Cm) | BsaI | Amplifying chloramphenicol resistance marker from pRK2 (Cm) for assembly of pAGRObb_000 via golden gate assembly using BsaI |
| colE1_only_amp-F | GTCTGGTCTCTTCCTGTCCGATTAGCGACATAGTCATC | pMOD_B2303 | BsaI | Amplifying the colE1 ori fragment from pMOD_B2303 for assembly of pAGRObb_[00X] series via golden gate assembly using BsaI |
| colE1_only_amp-R | GTCTGGTCTCTGGCGAATGGTAGCTGACAAGCTCTG | pMOD_B2303 | BsaI | Amplifying the colE1 ori fragment from pMOD_B2303 for assembly of pAGRObb_[00X] series via golden gate assembly using BsaI |
| pBBR_BB_amp-F | GTCTGGTCTCTTCCTAATAGAGTAACCCCCTACGGG | pFlp-Km | BsaI | Amplifying the BBR replicon fragment from pFlp-Km for assembly of pAGRObb_[10X] series via golden gate assembly using BsaI |
| pBBR_BB_amp-R | GTCTGGTCTCTGGCGAATGGTAGCTGACAAGCTCTATCAACGGG | pFlp-Km | BsaI | Amplifying the BBR replicon fragment from pFlp-Km for assembly of pAGRObb_[10X] series via golden gate assembly using BsaI |
| pRK2_BB_amp-F | GTCTGGTCTCTTCCTGGAGATCAAAGGATCTTCTTGAGATCC | pRK2 (S) | BsaI | Amplifying the RK2 replicon fragment from pRK2 (S) for assembly of pAGRObb_[20X] series via golden gate assembly using BsaI |
| pRK2_BB_amp-R | GTCTGGTCTCTGGCGAATGGTAGCTGACAAGCTCTCTTAGGGATA | pRK2 (S) | BsaI | Amplifying the RK2 replicon fragment from pRK2 (S) for assembly of pAGRObb_[20X] series via golden gate assembly using BsaI |
| pVS1_BB-amp-F | GTCTGGTCTCTTCCTCAGAGCATTACGCTGACTTGA | pMIN-VS1 | BsaI | Amplifying the VS1 replicon fragment from pMIN-VS1 for assembly of pAGRObb_[30X] series via golden gate assembly using BsaI |
| pVS1_BB-amp-R | GTCTGGTCTCTGGCGAATGGTAGCTGACAAGCTCTGTAGATTGCC | pMIN-VS1 | BsaI | Amplifying the VS1 replicon fragment from pMIN-VS1 for assembly of pAGRObb_[30X] series via golden gate assembly using BsaI |
| pSa_BB_amp-F | GTCTGGTCTCTTCCTGATCGAGCGGCATTTCATTCAGG | pWLlR5 | BsaI | Amplifying the Sa replicon fragment from pWLlR5 for assembly of pAGRObb_[40X] series via golden gate assembly using BsaI |
| pSa_BB_amp-R | GTCTGGTCTCTGGCGAATGGTAGCTGACAAGCTCTTTGTGGGCA | pWLlR5 | BsaI | Amplifying the Sa replicon fragment from pWLlR5 for assembly of pAGRObb_[40X] series via golden gate assembly using BsaI |
| kanR_amp0-F | GTCTGGTCTCTTTCGCTTGTGTCTCAAAATCTCTGATGTTAC | pMIN-VS1 | BsaI | Amplifying kanamycin resistance marker from pMIN-VS1 for assembly of pAGRObb_[X01] series and pAGROex_[2X1] series via golden gate assembly using BsaI |
| kanR_amp0-R | GTCTGGTCTCTAGGAAATTGATTCAACAAAGCCGC | pMIN-VS1 | BsaI | Amplifying kanamycin resistance marker from pMIN-VS1 for assembly of pAGRObb_[X01] series and pAGROex_[2X1] series via golden gate assembly using BsaI |
| specR_amp0-F | GTCTGGTCTCTTTCGCCTGCTTTTGATCGGTACCA | pVS-rec (S) | BsaI | Amplifying spectinomycin resistance marker from pVS-rec (S) for assembly of pAGRObb_[X02] series and pAGROex_[2X2] series via golden gate assembly using BsaI |
| specR_amp0-R | GTCTGGTCTCTAGGAGTAGGGCTTATTATGCACGCTT | pVS-rec (S) | BsaI | Amplifying spectinomycin resistance marker from pVS-rec (S) for assembly of pAGRObb_[X02] series and pAGROex_[2X2] series via golden gate assembly using BsaI |
| gentR_amp0-F | GTCTGGTCTCTTTCGCACCGTGGAAACGGATGAA | pRK2 (Ge) | BsaI | Amplifying gentamicin resistance marker from pRK2 (Ge) for assembly of pAGRObb_[X03] series and pAGROex_[2X3] series via golden gate assembly using BsaI |
| gentR_amp0-R | GTCTGGTCTCTAGGACGTTGTGACAATTTACCGAACAAC | pRK2 (Ge) | BsaI | Amplifying gentamicin resistance marker from pRK2 (Ge) for assembly of pAGRObb_[X03] series and pAGROex_[2X3] series via golden gate assembly using BsaI |
| apraR_amp0-F | GTCTGGTCTCTTTCGCACTGCTCAGACCAAGGATA | PN25::ApmA::T0term_RhiLeg-co | BsaI | Amplifying apramycin resistance marker from PN25::ApmA::T0term_RhiLeg-co_Paq_s1-3 for assembly of pAGRObb_[X04] series and pAGROex_[2X4] series via golden gate assembly using BsaI |
| apraR_amp0-R | GTCTGGTCTCTAGGACAGTTTGCTGAGATTGC | PN25::ApmA::T0term_RhiLeg-co | BsaI | Amplifying apramycin resistance marker from PN25::ApmA::T0term_RhiLeg-co_Paq_s1-3 for assembly of pAGRObb_[X04] series and pAGROex_[2X4] series via golden gate assembly using BsaI |
| sacB_BBRbb_amp-F | GTCTGGTCTCTGCGACCGAAGTTCCTATTCTCTAGGG | pFlp-Km | BsaI | Amplifying a fragment consisting of the sacB counterselection marker and the BBR replicon from pFlp-Km for assembly of all pAGROex_[2XX] via golden gate assembly using BsaI |
| sacB_BBRbb_amp-R | GTCTGGTCTCTCGAAGCTACTTTTCCGCTGCATAACC | pFlp-Km | BsaI | Amplifying a fragment consisting of the sacB counterselection marker and the BBR replicon from pFlp-Km for assembly of all pAGROex_[2XX] via golden gate assembly using BsaI |
| eforRed_cass_amp0-F | GTCTGGTCTCTACCGCCAACCGAAAGGTGACTCTA | pVS-eforRed-3-purged | BsaI | Amplifying the eforRed expression cassette from pVS-eforRed-3-purged for assembly of all pAGROex_[2XX] via golden gate assembly using BsaI |
| eforRed_cass_amp0-R | GTCTGGTCTCTTCGCGTCCACATTGCGTAGACTGT | pVS-eforRed-3-purged | BsaI | Amplifying the eforRed expression cassette from pVS-eforRed-3-purged for assembly of all pAGROex_[2XX] via golden gate assembly using BsaI |
| pBBR_bb_FLP_amp-F | GTCTGGTCTCTTCCTCAATTGCCTGGATTCTCACCAA | pFlp-Km | BsaI | Amplifying the FLP recombinase expression cassette from pFlp-Km for assembly of pAGROex_[20X] series via golden gate assembly using BsaI |
| pBBR_bb_FLP_amp-R | GTCTGGTCTCTCGGTGTCGGAATAGGAACTTCAAAGCG | pFlp-Km | BsaI | Amplifying the FLP recombinase expression cassette from pFlp-Km for assembly of pAGROex_[20X] series via golden gate assembly using BsaI |
| Cre-cass_amp-F | GTCTGGTCTCTTCCTCTTTACGGCTAGCTCAGCC | pVS-Cre-A-purged | BsaI | Amplifying the Cre recombinase expression cassette from pVS-Cre-A-purged for assembly of pAGROex_[21X] series via golden gate assembly using BsaI |
| Cre-cass_amp-R | GTCTGGTCTCTCGGTGTCCACATTGCGTAGACTGT | pVS-Cre-A-purged | BsaI | Amplifying the Cre recombinase expression cassette from pVS-Cre-A-purged for assembly of pAGROex_[21X] series via golden gate assembly using BsaI |

| | | | | |
|---|---|---|---|---|
| lacZa_Gib_FRT_HR-F | CTCTAGAAAGTATAGGAACTTCCCCGGATAATGAGACCTTAGCG | lacZa_MCS_del (synthetic fragm | N/A | Amplifying the lacZα fragment from lacZa_MCS_del flanked on the upstream end with a FRT site for Gibson assembly with an XmaI-digested pAGROm2_102 backbone fragment, yielding pAGROm2_102LC |
| lacZa_Gib_LoxP_HR-F | AATGTATGCTATACGAAGTTATCCCGGATAATGAGACCTTAGCG | lacZa_MCS_del (synthetic fragm | N/A | Amplifying the lacZα fragment from lacZa_MCS_del flanked on the upstream end with a LoxP site for Gibson assembly with an XmaI-digested pAGROm2_112 backbone fragment, yielding pAGROm2_112LC |
| lacZa_Gib_Um2R_HR-R | GAGTCTTTCACGCTCTTACAGGACGTACGTTCATGAGACCTTGGA | lacZa_MCS_del (synthetic fragm | N/A | Amplifying the lacZα fragment from lacZa_MCS_del flanked on the upstream end with either a FRT or LoxP site for Gibson assembly with an XmaI-digested pAGROm2_102 or pAGROm2_112 backbone fragment, yielding pAGROm2_102LC or pAGROm2_112LC, respectively |
| dCas9_CBE_amp-1F | GTCTGGTCTCTCGTAGGCGCGCCTCGAGCATTTTCGTTCTGAG | pVP143 | BsaI | Amplifying an upstream segment of the dCas9-CDA-UGI expression cassette from pVP143 for assembly with the downstream segment and the mod1_BB_amp fragment to make pAGROm1_cbe0 via golden gate assembly using BsaI |
| dCas9_CBE_amp-1R | GTCTGGTCTCTCTCTTTAAATGTCAAACTATCATCATGGATC | pVP143 | BsaI | Amplifying an upstream segment of the dCas9-CDA-UGI expression cassette from pVP143 for assembly with the downstream segment and the mod1_BB_amp fragment to make pAGROm1_cbe0 via golden gate assembly using BsaI |
| dCas9_CBE_amp-2F | GTCTGGTCTCTAGAGGACATTCAAAAAGCACAAGTGTC | pVP143 | BsaI | Amplifying a downstream segment of the dCas9-CDA-UGI expression cassette from pVP143 for assembly with the upstream segment and the mod1_BB_amp fragment to make pAGROm1_cbe0 via golden gate assembly using BsaI |
| dCas9_CBE_amp-2R | GTCTGGTCTCTTGATCCCGGGTTGACGTCTGTACAAACTTGCATA | pVP143 | BsaI | Amplifying a downstream segment of the dCas9-CDA-UGI expression cassette from pVP143 for assembly with the upstream segment and the mod1_BB_amp fragment to make pAGROm1_cbe0 via golden gate assembly using BsaI |
| ddCas9-frag1-F | GCTTGGTCTCTCTAATCAGAATAAGCAAACCCCTT | pAGROm1_cbe0 | BsaI | Amplifying the Cas9 coding sequence, expression elements, and backbone from pAGROm1_cbe0, reverting the D10A and H840A dCas9 mutations back to the wild-type for a 3-piece assembly yielding pAGROm1_cas0 via golden gate assembly using BsaI |
| ddCas9-frag1-R | GCTTGGTCTCTTGTCTAAGCCTATTGAGTATTTCTTATCC | pAGROm1_cbe0 | BsaI | Amplifying the Cas9 coding sequence, expression elements, and backbone from pAGROm1_cbe0, reverting the D10A and H840A dCas9 mutations back to the wild-type for a 3-piece assembly yielding pAGROm1_cas0 via golden gate assembly using BsaI |
| ddCas9-frag2-F | GCTTGGTCTCTGACATCGGCACAAATAGCGT | pAGROm1_cbe0 | BsaI | Amplifying the Cas9 coding sequence, expression elements, and backbone from pAGROm1_cbe0, reverting the D10A and H840A dCas9 mutations back to the wild-type for a 3-piece assembly yielding pAGROm1_cas0 via golden gate assembly using BsaI |
| ddCas9-frag2-R | GCTTGGTCTCTGTGATCGACATCATAATCACTTAAACG | pAGROm1_cbe0 | BsaI | Amplifying the Cas9 coding sequence, expression elements, and backbone from pAGROm1_cbe0, reverting the D10A and H840A dCas9 mutations back to the wild-type for a 3-piece assembly yielding pAGROm1_cas0 via golden gate assembly using BsaI |
| ddCas9-frag3-F | GCTTGGTCTCTTCACATTGTTCCACAAAGTTTCCT | pAGROm1_cbe0 | BsaI | Amplifying the Cas9 coding sequence, expression elements, and backbone from pAGROm1_cbe0, reverting the D10A and H840A dCas9 mutations back to the wild-type for a 3-piece assembly yielding pAGROm1_cas0 via golden gate assembly using BsaI |
| ddCas9-frag3-R | GCTTGGTCTCTTTAGTCACCTCCTAGCTGAC | pAGROm1_cbe0 | BsaI | Amplifying the Cas9 coding sequence, expression elements, and backbone from pAGROm1_cbe0, reverting the D10A and H840A dCas9 mutations back to the wild-type for a 3-piece assembly yielding pAGROm1_cas0 via golden gate assembly using BsaI |
| sfGFP_BbsI_sgRNA_ph-F | GTTCCTGAAGACTTAGTCGACAGTTGACAGCTAGCTCAGTCCT | pEN-L1-PJ23119-PplpT-sfGFP-T | BbsI | Amplifying the sgRNA spacer placeholder and scaffold sequence from pEN-L1-PJ23119-PplpT-sfGFP-TrrfB-BsaI-Scaf-L2 for assembly with either the m2BB_Bbs_sgR_amp or m3BB_Bbs_sgR_amp fragments to make pmod2_sgRNA_ph or pmod3_sgRNA_ph, respectively, via golden gate assembly using BbsI |
| sfGFP_BbsI_sgRNA_ph-R | GTTCCTGAAGACTTGCAGTCGTACAAGAAAGCTGGGTAATAC | pEN-L1-PJ23119-PplpT-sfGFP-T | BbsI | Amplifying the sgRNA spacer placeholder and scaffold sequence from pEN-L1-PJ23119-PplpT-sfGFP-TrrfB-BsaI-Scaf-L2 for assembly with either the m2BB_Bbs_sgR_amp or m3BB_Bbs_sgR_amp fragments to make pmod2_sgRNA_ph or pmod3_sgRNA_ph, respectively, via golden gate assembly using BbsI |
| m2BB_BbsI_sgRNA_amp-F | GTTCCTGAAGACTTCTGCCCGCAAAAGCAGGTGTGTCAGGTGGC | pMOD_B2303 | BbsI | Amplifying the carbencillin resistance marker and colE1 ori sequence fragment from pMOD_B2303, installing PaqCI sites and overhangs for module 2 for assembly with the sfGFP_BbsI-sgRNA_ph fragment, yielding pmod2_sgRNA_ph via golden gate assembly using BbsI |
| m2BB_BbsI_sgRNA_amp-R | GTTCCTGAAGACTTGACTTGATTTTTGCAGGTGTGGCGCTCTCCC | pMOD_B2303 | BbsI | Amplifying the carbencillin resistance marker and colE1 ori sequence fragment from pMOD_B2303, installing PaqCI sites and overhangs for module 2 for assembly with the sfGFP_BbsI-sgRNA_ph fragment, yielding pmod2_sgRNA_ph via golden gate assembly using BbsI |
| m3BB_BbsI_sgRNA_amp-F | GTTCCTGAAGACTTCTGCGGGAAAAAGCAGGTGTGTCAGGTGGC | pMOD_B2303 | BbsI | Amplifying the carbencillin resistance marker and colE1 ori sequence fragment from pMOD_B2303, installing PaqCI sites and overhangs for module 3 for assembly with the sfGFP_BbsI-sgRNA_ph fragment, yielding pmod3_sgRNA_ph via golden gate assembly using BbsI |
| m3BB_BbsI_sgRNA_amp-R | GTTCCTGAAGACTTGACTGCGGTTTTGCAGGTGTGGCGCTCTCC | pMOD_B2303 | BbsI | Amplifying the carbencillin resistance marker and colE1 ori sequence fragment from pMOD_B2303, installing PaqCI sites and overhangs for module 3 for assembly with the sfGFP_BbsI-sgRNA_ph fragment, yielding pmod3_sgRNA_ph via golden gate assembly using BbsI |
| sgRNA_ph_cass_rep-F | GTTCCTGAAGACTTTAGCGGAGACCAATTTATCTTCTACAAGGAG | pVS-sfGFP-2-purged | BbsI | Amplifying the sfGFP coding sequence and terminator from pVS-sfGFP-2-purged, digesting it with BbsI, and then ligating this into BsaI-digested backbone fragments of either pmod2_sgRNA_ph or pmod3_sgRNA_ph, yielding pAGROm2_cas0 or pAGROm3_cas0, respectively |
| sgRNA_ph_cass_rep-R | GTTCCTGAAGACTTAAACTGAGACCGTCCACATTGCGTAGACTGT | pVS-sfGFP-2-purged | BbsI | Amplifying the sfGFP coding sequence and terminator from pVS-sfGFP-2-purged, digesting it with BbsI, and then ligating this into BsaI-digested backbone fragments of either pmod2_sgRNA_ph or pmod3_sgRNA_ph, yielding pAGROm2_cas0 or pAGROm3_cas0, respectively |
| J23100_SalI_NheI_oligo-F | TCGACAGTTGACGGCTAGCTCAGTCCTAGGTACAGTG | N/A (used for annealing with par | N/A | Ligating annealed dsOligo pair containing the PJ23100 promoter sequence into SalI and NheI-digested pAGROm2_cas0 or pAGROm3_cas0 backbone fragments, yielding pAGROm2_cas1 and pAGROm3_cas1, respectively |
| J23100_SalI_NheI_oligo-R | CTAGCACTGTACCTAGGACTGAGCTAGCCGTCAACTG | N/A (used for annealing with par | N/A | Ligating annealed dsOligo pair containing the PJ23100 promoter sequence into SalI and NheI-digested pAGROm2_cas0 or pAGROm3_cas0 backbone fragments, yielding pAGROm2_cas1 and pAGROm3_cas1, respectively |
| J23107_SalI_NheI_oligo-F | TCGACAGTTTACGGCTAGCTCAGCCCTAGGTATTATG | N/A (used for annealing with par | N/A | Ligating annealed dsOligo pair containing the PJ23107 promoter sequence into SalI and NheI-digested pAGROm2_cas0 or pAGROm3_cas0 backbone fragments, yielding pAGROm2_cas2 and pAGROm3_cas2, respectively |
| J23107_SalI_NheI_oligo-R | CTAGCATAATACCTAGGGCTGAGCTAGCCGTAAACTG | N/A (used for annealing with par | N/A | Ligating annealed dsOligo pair containing the PJ23107 promoter sequence into SalI and NheI-digested pAGROm2_cas0 or pAGROm3_cas0 backbone fragments, yielding pAGROm2_cas2 and pAGROm3_cas2, respectively |
| m1BB_INTEGRATE_amp-F | GTCTGGTCTCTATCAAAAAGCAGGTGTGTCAGGTGGCACTTTTCG | pMOD_B2303 | BsaI | Amplifying the carbencillin resistance marker and colE1 ori sequence fragment from pMOD_B2303, installing PaqCI sites and overhangs for module 1 for assembly with the INTEGRATE_mach_amp fragment, yielding pAGROm1_int0 via golden gate assembly using BsaI |
| m1BB_INTEGRATE_amp-R | GTCTGGTCTCTTACGTTTTGCAGGTGTGGCGCTCTCCCGCTT | pMOD_B2303 | BsaI | Amplifying the carbencillin resistance marker and colE1 ori sequence fragment from pMOD_B2303, installing PaqCI sites and overhangs for module 1 for assembly with the INTEGRATE_mach_amp fragment, yielding pAGROm1_int0 via golden gate assembly using BsaI |
| INTEGRATE_mach_amp-F | GTCTGGTCTCTCGTATGGCGCGCCTACTTTGATCCCGAGGGGAA | pEA186 | BsaI | Amplifying the INTEGRATE machinery polycistron fragment from pEA186 for assembly with the m1_INTEGRATE_BB_amp fragment, yielding pAGROm1_int0 via golden gate assembly using BsaI |
| INTEGRATE_mach_amp-R | GTCTGGTCTCTTGATTCCTGCAGGCGTCAGATTAAGGGTACAGG | pEA186 | BsaI | Amplifying the INTEGRATE machinery polycistron fragment from pEA186 for assembly with the m1_INTEGRATE_BB_amp fragment, yielding pAGROm1_int0 via golden gate assembly using BsaI |
| m2BB_crRNA_amp-F | GTTCCTGAAGACTTCCGCAAAAGCAGGTGTGTCAGGTGGCACTT | pMOD_B2303 | BbsI | Amplifying the carbencillin resistance marker and colE1 ori sequence fragment from pMOD_B2303, installing PaqCI sites and overhangs for module 2 for assembly with the crRNA_ph_amp fragment, yielding pmod2_INT_crRNA via golden gate assembly using BbsI |
| m2BB_crRNA_amp-R | GTTCCTGAAGACTTTGATTTTTGCAGGTGTGGCGCTCTCCCGCTT | pMOD_B2303 | BbsI | Amplifying the carbencillin resistance marker and colE1 ori sequence fragment from pMOD_B2303, installing PaqCI sites and overhangs for module 2 for assembly with the crRNA_ph_amp fragment, yielding pmod2_INT_crRNA via golden gate assembly using BbsI |
| crRNA_ph_amp-F | GTTCCTGAAGACTTATCAGGCGCGCCTAATGGGTGAACTGCCGA | pEA186 | BbsI | Amplifying the crRNA spacer entry fragment from pEA186 for assembly with the m2_crRNA_BB_amp fragment, yielding pmod2_INT_crRNA via golden gate assembly using BbsI |
| crRNA_ph_amp-R | GTTCCTGAAGACTTGCGGGCGGCCGCTTATGCCTGAGGTTTCAG | pEA186 | BbsI | Amplifying the crRNA spacer entry fragment from pEA186 for assembly with the m2_crRNA_BB_amp fragment, yielding pmod2_INT_crRNA via golden gate assembly using BbsI |
| sfGFP_BbsI_crRNA_amp-F | GTTCCTGAAGACCT ATAACGAGACCAACCGAAAGGTGACTCTA | pVS-sfGFP-3 | BbsI | Amplifying the sfGFP expression cassette from pVS-sfGFP-3, digesting it with BbsI, and then ligating this into a BsaI-digested backbone fragment of pmod2_INT_crRNA, yielding pAGROm2_int0 |
| sfGFP_BbsI_crRNA_amp-R | GTTCCTGAAGACAGTTCACGAGACCGTCCACATTGCGTAGACTG | pVS-sfGFP-3 | BbsI | Amplifying the sfGFP expression cassette from pVS-sfGFP-3, digesting it with BbsI, and then ligating this into a BsaI-digested backbone fragment of pmod2_INT_crRNA, yielding pAGROm2_int0 |
| m3BB_cargo_amp-F | GTCTGGTCTCTGGGAAAAAGCAGGTGTGTCAGGTGGCACTTTTC | pMOD_B2303 | BsaI | Amplifying the carbencillin resistance marker and colE1 ori sequence fragment from pMOD_B2303, installing PaqCI sites and overhangs for module 3 for assembly with the INTEG_cargo_amp fragment, or one of the three recombineering system fragments (Ptet-pluγ-RecET_rhi145, Ptet-pluγ-RecETh_rhi483, Ptet-RecETh1h2h3h4_agroB6) yielding pmod3_INT_cargo, pAGROm3_rec145, pAGROm3_rec483, and pAGROm3_recB6, respectively, via golden gate assembly using BsaI |
| m3BB_cargo_amp-R | GTCTGGTCTCTGCGGTTTTGCAGGTGTGGCGCTCTCCCGCTT | pMOD_B2303 | BsaI | Amplifying the carbencillin resistance marker and colE1 ori sequence fragment from pMOD_B2303, installing PaqCI sites and overhangs for module 3 for assembly with the INTEG_cargo_amp fragment, or one of the three recombineering system fragments (Ptet-pluγ-RecET_rhi145, Ptet-pluγ-RecETh_rhi483, Ptet-RecETh1h2h3h4_agroB6) yielding pmod3_INT_cargo, pAGROm3_rec145, pAGROm3_rec483, and pAGROm3_recB6, respectively, via golden gate assembly using BsaI |
| INTEGRATE_cargo_amp-F | GTCTGGTCTCTCCGCGGCGCGCCGCTGAAACCTCAGGCATTTG | pEA186 | BsaI | Amplifying the INTEGRATE transposon cargo fragment from pEA186 for assembly with the m3_cargo_BB_amp fragment, yielding pmod3_INT_cargo via golden gate assembly using BsaI |
| INTEGRATE_cargo_amp-R | GTCTGGTCTCTTCCCGCGGCCGCCTCCCTGTTGATGCAACCATA | pEA186 | BsaI | Amplifying the INTEGRATE transposon cargo fragment from pEA186 for assembly with the m3_cargo_BB_amp fragment, yielding pmod3_INT_cargo via golden gate assembly using BsaI |
| lacZa_ph_Gib_BbsI_amp-F | CGACATTTTCCTCGAGGTCATTTCAAGGTAGTCTTCAAAGCGCAA | lacZa_MCS_del (synthetic fragm | N/A | Amplifying the lacZα fragment from lacZa_MCS_del with added divergent BbsI sites and overhangs for Gibson assembly with a PstI and XhoI-digested pmod3_INT_cargo backbone fragment, yielding pAGROm3_int0 |
| lacZa_ph_Gib_BbsI_amp-R | ATTTCAGCAAAACTACTGCAGTAAGGAACTAGTCTTCAATGGACT | lacZa_MCS_del (synthetic fragm | N/A | Amplifying the lacZα fragment from lacZa_MCS_del with added divergent BbsI sites and overhangs for Gibson assembly with a PstI and XhoI-digested pmod3_INT_cargo backbone fragment, yielding pAGROm3_int0 |
| Ptet_[tonB]_amp-F | GTCTGGTCTCTCCGCAGTCAAAAGCCTCCGGTCGGAGGCTTTTG | pBBR1-kan-Ptet-plu-gamma-ET_ | BsaI | Amplifying the TetR/TetO sequence fragment from pBBR1-kan-Ptet-plu-gamma-ET_rhi145 with the bidirectional tonB terminator added at the upstream end for assembly with the m3_cargo_BB_amp fragment, the rrnB_T1T2_term fragment, and segments of either pluγ-RecET_rhi145, pluγ-RecETh_rhi483, or RecETh1h2h3h4_agroB6 recombineering systems yielding pAGROm3_rec145, pAGROm3_rec483, and pAGROm3_recB6, respectively, via golden gate assembly using BsaI |
| Ptet_amp-R | GTCTGGTCTCTATTCTCTCTATCACTGATAGGGAGTG | pBBR1-kan-Ptet-plu-gamma-ET_ | BsaI | Amplifying the TetR/TetO sequence fragment from pBBR1-kan-Ptet-plu-gamma-ET_rhi145 with the bidirectional tonB terminator added at the upstream end for assembly with the m3_cargo_BB_amp fragment, the rrnB_T1T2_term fragment, and segments of either pluγ-RecET_rhi145, pluγ-RecETh_rhi483, or RecETh1h2h3h4_agroB6 recombineering systems yielding pAGROm3_rec145, pAGROm3_rec483, and pAGROm3_recB6, respectively, via golden gate assembly using BsaI |
| plu-gamma-F | GTCTGGTCTCTGAATTCATATGAACCCATATGCAGTTTATG | pBBR1-kan-Ptet-plu-gamma-ET_ | BsaI | Amplifying a fragment containing the *Photorhabdus luminescens* "γ" ssDNA binding protein open reading frame and a segment of either the RecET gene cluster from *Rhizobium* sp. LC145 or the RecETh cluster from *Rhizobium* sp. Root483D2 from pBBR1-kan-Ptet-plu-gamma-ET_rhi145 or pBBR1-kan-Ptet-plu-gamma-ETh_rhi483 for assembly with the m3_cargo_BB_amp fragment, the rrnB_T1T2_term fragment, and the TetR/TetO sequence fragment with the remaining segments of either RecET_rhi145 or RecETh_rhi483 recombineering systems, yielding pAGROm3_rec145 or pAGROm3_rec483, respectively, via golden gate assembly using BsaI |
| rhi145_GG_elim_p1-R | GTCTGGTCTCTCGTGACAGATATCGGCTGGTT | pBBR1-kan-Ptet-plu-gamma-ET_ | BsaI | Amplifying a fragment containing the *Photorhabdus luminescens* "γ" ssDNA binding protein open reading frame and a segment of the RecET gene cluster from *Rhizobium* sp. LC145 from pBBR1-kan-Ptet-plu-gamma-ET_rhi145, eliminating internal PaqCI and BsaI recognition sequences, for assembly with the m3_cargo_BB_amp fragment, the rrnB_T1T2_term fragment, the TetR/TetO sequence fragment, and the remaining segments of the RecET_rhi145 recombineering system, yielding pAGROm3_rec145 via golden gate assembly using BsaI |
| rhi145_GG_elim_p2-F | GTCTGGTCTCTCACGTGCATCATCGAGCATTCCG | pBBR1-kan-Ptet-plu-gamma-ET_ | BsaI | Amplifying a second segment of the RecET gene cluster from *Rhizobium* sp. LC145 from pBBR1-kan-Ptet-plu-gamma-ET_rhi145, eliminating internal PaqCI and BsaI recognition sequences, for assembly with the m3_cargo_BB_amp fragment, the rrnB_T1T2_term fragment, the TetR/TetO sequence fragment, and the remaining segments of the RecET_rhi145 recombineering system, yielding pAGROm3_rec145 via golden gate assembly using BsaI |

| | | | | |
|---|---|---|---|---|
| rhi145_GG_elim_p2-R | GTCTGGTCTCTGCTCCGCGATCAGCTCG | pBBR1-kan-Ptet-plu-gamma-ET_ | BsaI | Amplifying a second segment of the RecET gene cluster from *Rhizobium* sp. LC145 from pBBR1-kan-Ptet-plu-gamma-ET_rhi145, eliminating internal PaqCI and BsaI recognition sequences, for assembly with the m3_cargo_BB_amp fragment, the rrnB_T1T2_term fragment, the TetR/TetO sequence fragment, and the remaining segments of the RecET_rhi145 recombineering system, yielding pAGROm3_rec145 via golden gate assembly using BsaI |
| rhi145_GG_elim_p3-F | GTCTGGTCTCTGAGCGCCTGACGGGTGTTG | pBBR1-kan-Ptet-plu-gamma-ET_ | BsaI | Amplifying a third segment of the RecET gene cluster from *Rhizobium* sp. LC145 from pBBR1-kan-Ptet-plu-gamma-ET_rhi145, eliminating internal PaqCI and BsaI recognition sequences, for assembly with the m3_cargo_BB_amp fragment, the rrnB_T1T2_term fragment, the TetR/TetO sequence fragment, and the remaining segments of the RecET_rhi145 recombineering system, yielding pAGROm3_rec145 via golden gate assembly using BsaI |
| rhi145_term_p3-R | GTCTGGTCTCTCTGTCTCCTTTATGCGGCCTCC | pBBR1-kan-Ptet-plu-gamma-ET_ | BsaI | Amplifying a third segment of the RecET gene cluster from *Rhizobium* sp. LC145 from pBBR1-kan-Ptet-plu-gamma-ET_rhi145, eliminating internal PaqCI and BsaI recognition sequences, for assembly with the m3_cargo_BB_amp fragment, the rrnB_T1T2_term fragment, the TetR/TetO sequence fragment, and the remaining segments of the RecET_rhi145 recombineering system, yielding pAGROm3_rec145 via golden gate assembly using BsaI |
| rhi483_GG_elim_p1-R | GTCTGGTCTCTGCCTTCGACGTCCACCA | pBBR1-kan-Ptet-plu-gamma-ET | BsaI | Amplifying a fragment containing the *Photorhabdus luminescens* "γ" ssDNA binding protein open reading frame and a segment of the RecETh cluster from *Rhizobium* sp. Root483D2 from pBBR1-kan-Ptet-plu-gamma-ETh_rhi483, eliminating internal PaqCI and BsaI recognition sequences, for assembly with the m3_cargo_BB_amp fragment, the rrnB_T1T2_term fragment, the TetR/TetO sequence fragment, and the remaining segment of the RecETh_rhi483 recombineering system, yielding pAGROm3_rec483 via golden gate assembly using BsaI |
| rhi483_GG_elim_p2-F | GTCTGGTCTCTAGGCCTCTTGGAGATTAAGTGCCCC | pBBR1-kan-Ptet-plu-gamma-ET | BsaI | Amplifying a second segment of the RecETh cluster from *Rhizobium* sp. Root483D2 from pBBR1-kan-Ptet-plu-gamma-ETh_rhi483, eliminating internal PaqCI and BsaI recognition sequences, for assembly with the m3_cargo_BB_amp fragment, the rrnB_T1T2_term fragment, the TetR/TetO sequence fragment, and the remaining segment of the RecETh_rhi483 recombineering system, yielding pAGROm3_rec483 via golden gate assembly using BsaI |
| rhi483_term_p2-R | GTCTGGTCTCTCTGTCTCCTTCATGACGCCCT | pBBR1-kan-Ptet-plu-gamma-ET | BsaI | Amplifying a second segment of the RecETh cluster from *Rhizobium* sp. Root483D2 from pBBR1-kan-Ptet-plu-gamma-ETh_rhi483, eliminating internal PaqCI and BsaI recognition sequences, for assembly with the m3_cargo_BB_amp fragment, the rrnB_T1T2_term fragment, the TetR/TetO sequence fragment, and the remaining segment of the RecETh_rhi483 recombineering system, yielding pAGROm3_rec483 via golden gate assembly using BsaI |
| agrob6_start-F | GTCTGGTCTCTGAATTCATATGGAAAAGATAATCGACAAAGAC | pBBR1-kan-Ptet-RecETh1h2h3h | BsaI | Amplifying a segment of the RecETh1h2h3h4 gene cluster from *Agrobacterium tumefaciens* str. B6 from pBBR1-kan-Ptet-RecETh1h2h3h4_agrob6, eliminating internal PaqCI and BsaI recognition sequences, for assembly with the m3_cargo_BB_amp fragment, the rrnB_T1T2_term fragment, the TetR/TetO sequence fragment, and the remaining segments of the RecETh1h2h3h4_agrob6 recombineering system, yielding pAGROm3_recB6 via golden gate assembly using BsaI |
| agrob6_GG_elim_p1-R | GTCTGGTCTCTAGGGGATGCGGTGAACCCTTCG | pBBR1-kan-Ptet-RecETh1h2h3h | BsaI | Amplifying a segment of the RecETh1h2h3h4 gene cluster from *Agrobacterium tumefaciens* str. B6 from pBBR1-kan-Ptet-RecETh1h2h3h4_agrob6, eliminating internal PaqCI and BsaI recognition sequences, for assembly with the m3_cargo_BB_amp fragment, the rrnB_T1T2_term fragment, the TetR/TetO sequence fragment, and the remaining segments of the RecETh1h2h3h4_agrob6 recombineering system, yielding pAGROm3_recB6 via golden gate assembly using BsaI |
| agrob6_GG_elim_p2-F | GTCTGGTCTCTCCCTGCCGCTGATGAGC | pBBR1-kan-Ptet-RecETh1h2h3h | BsaI | Amplifying a second segment of the RecETh1h2h3h4 gene cluster from *Agrobacterium tumefaciens* str. B6 from pBBR1-kan-Ptet-RecETh1h2h3h4_agrob6, eliminating internal PaqCI and BsaI recognition sequences, for assembly with the m3_cargo_BB_amp fragment, the rrnB_T1T2_term fragment, the TetR/TetO sequence fragment, and the remaining segments of the RecETh1h2h3h4_agrob6 recombineering system, yielding pAGROm3_recB6 via golden gate assembly using BsaI |
| agrob6_GG_elim_p2-R | GTCTGGTCTCTCGCCTTCGCCTCAAGGG | pBBR1-kan-Ptet-RecETh1h2h3h | BsaI | Amplifying a second segment of the RecETh1h2h3h4 gene cluster from *Agrobacterium tumefaciens* str. B6 from pBBR1-kan-Ptet-RecETh1h2h3h4_agrob6, eliminating internal PaqCI and BsaI recognition sequences, for assembly with the m3_cargo_BB_amp fragment, the rrnB_T1T2_term fragment, the TetR/TetO sequence fragment, and the remaining segments of the RecETh1h2h3h4_agrob6 recombineering system, yielding pAGROm3_recB6 via golden gate assembly using BsaI |
| agrob6_GG_elim_p3-F | GTCTGGTCTCTGGCGGGTGCAGCATGACTCG | pBBR1-kan-Ptet-RecETh1h2h3h | BsaI | Amplifying a third segment of the RecETh1h2h3h4 gene cluster from *Agrobacterium tumefaciens* str. B6 from pBBR1-kan-Ptet-RecETh1h2h3h4_agrob6, eliminating internal PaqCI and BsaI recognition sequences, for assembly with the m3_cargo_BB_amp fragment, the rrnB_T1T2_term fragment, the TetR/TetO sequence fragment, and the remaining segments of the RecETh1h2h3h4_agrob6 recombineering system, yielding pAGROm3_recB6 via golden gate assembly using BsaI |
| agrob6_term_p3-R | GTCTGGTCTCTCTGTGCTCCTTAAGCCTAGGTCAT | pBBR1-kan-Ptet-RecETh1h2h3h | BsaI | Amplifying a third segment of the RecETh1h2h3h4 gene cluster from *Agrobacterium tumefaciens* str. B6 from pBBR1-kan-Ptet-RecETh1h2h3h4_agrob6, eliminating internal PaqCI and BsaI recognition sequences, for assembly with the m3_cargo_BB_amp fragment, the rrnB_T1T2_term fragment, the TetR/TetO sequence fragment, and the remaining segments of the RecETh1h2h3h4_agrob6 recombineering system, yielding pAGROm3_recB6 via golden gate assembly using BsaI |
| rrnB_T1T2_term-F | GTCTGGTCTCTACAGGAGAGTAGGGAACTGCCAG | pEN-L1-PJ23119-PplpT-sfGFP-T | BsaI | Amplifying a fragment containing the rrnB T1 and T2 terminator sequences from pEN-L1-PJ23119-PplpT-sfGFP-TrrfB-BsaI-Scaf-L2 for assembly with the m3_cargo_BB_amp fragment, the TetR/TetO sequence fragment, and segments of either pluγ-RecET_rhi145, pluγ-RecETh_rhi483, or RecETh1h2h3h4_agroB6 recombineering systems yielding pAGROm3_rec145, pAGROm3_rec483, and pAGROm3_recB6, respectively, via golden gate assembly using BsaI |
| rrnB_T1T2_term-R | GTCTGGTCTCTTCCCAAAAGGCCATCCGTCAGG | pEN-L1-PJ23119-PplpT-sfGFP-T | BsaI | Amplifying a fragment containing the rrnB T1 and T2 terminator sequences from pEN-L1-PJ23119-PplpT-sfGFP-TrrfB-BsaI-Scaf-L2 for assembly with the m3_cargo_BB_amp fragment, the TetR/TetO sequence fragment, and segments of either pluγ-RecET_rhi145, pluγ-RecETh_rhi483, or RecETh1h2h3h4_agroB6 recombineering systems yielding pAGROm3_rec145, pAGROm3_rec483, and pAGROm3_recB6, respectively, via golden gate assembly using BsaI |

| Primer name | Sequence | Template | Enzyme ends | Purpose |
|---|---|---|---|---|
| pTiC58_Left_HA-1200F | CTGTCACCTGCGCTTCGTACATCACATCCCAGGTCGAA | strain C58 genomic DNA; strain | PaqCI | Amplifying a Left homology arm fragment ~1,200 bp in length from pTiC58 for assembly with the corresponding ~1,200 bp Right homology arm fragment, pAGROm2_112, pAGROm4_210, and pAGRObb_000, yielding pDisarm_C58_112_210_000, via golden gate assembly using PaqCI; amplifying a Left homology arm fragment from pTi3a for assembly with the corresponding Right homology arm fragment, pAGROm2_112, pAGROm4_210, and pAGRObb_000, yielding pDisarm_3a_112_210_000, via golden gate assembly using PaqCI; amplifying a Left homology arm fragment from pTiC58 for assembly with the corresponding Right homology arm fragment, pAGROm2_002, pAGROm4_010, and pAGRObb_000, yielding pDisarm_C58_002_010_000, via golden gate assembly using PaqCI (used as a control construct without visual reporter cassettes) |
| pTiC58_Left_HA-600F | CTGTCACCTGCGCTTCGTACGATCTTCTCGACAATGGCA | strain C58 genomic DNA; strain | PaqCI | Amplifying a Left homology arm fragment ~600 bp in length from pTiC58 for assembly with the corresponding ~600 bp Right homology arm fragment, pAGROm2_112, pAGROm4_210, and pAGRObb_000, yielding pDisarm_C58_112_210_000 [600 HA], via golden gate assembly using PaqCI |
| pTiC58_Left_HA-300F | CTGTCACCTGCGCTTCGTAATGTTCATCGCATCGAGGT | strain C58 genomic DNA | PaqCI | Amplifying a Left homology arm fragment ~300 bp in length from pTiC58 for assembly with the corresponding ~300 bp Right homology arm fragment, pAGROm2_112, pAGROm4_210, and pAGRObb_000, yielding pDisarm_C58_112_210_000 [300 HA], via golden gate assembly using PaqCI |
| pTiC58_Left_HA-R | CTGTCACCTGCGCTTTGATCCATCATCTCCCTCGCAGAGA | strain C58 genomic DNA | PaqCI | Amplifying a Left homology arm fragment ~1,200 bp in length from pTiC58 for assembly with the corresponding ~1,200 bp Right homology arm fragment, pAGROm2_112, pAGROm4_210, and pAGRObb_000, yielding pDisarm_C58_112_210_000, via golden gate assembly using PaqCI; amplifying a Left homology arm fragment from pTi3a for assembly with the corresponding Right homology arm fragment, pAGROm2_112, pAGROm4_210, and pAGRObb_000, yielding pDisarm_3a_112_210_000, via golden gate assembly using PaqCI; amplifying a Left homology arm fragment ~600 bp in length from pTiC58 for assembly with the corresponding ~600 bp Right homology arm fragment, pAGROm2_112, pAGROm4_210, and pAGRObb_000, yielding pDisarm_C58_112_210_000 [600 HA], via golden gate assembly using PaqCI; amplifying a Left homology arm fragment ~300 bp in length from pTiC58 for assembly with the corresponding ~300 bp Right homology arm fragment, pAGROm2_112, pAGROm4_210, and pAGRObb_000, yielding pDisarm_C58_112_210_000 [300 HA], via golden gate assembly using PaqCI |
| pTiC58_Right_HA-F | CTGTCACCTGCGCTTCCGCTTCAAACGTCGGAAAGCCTG | strain C58 genomic DNA; strain | PaqCI | Amplifying a Right homology arm fragment ~1,200 bp in length from pTiC58 for assembly with the corresponding ~1,200 bp Left homology arm fragment, pAGROm2_112, pAGROm4_210, and pAGRObb_000, yielding pDisarm_C58_112_210_000, via golden gate assembly using PaqCI; amplifying a Right homology arm fragment from pTi3a for assembly with the corresponding Left homology arm fragment, pAGROm2_112, pAGROm4_210, and pAGRObb_000, yielding pDisarm_3a_112_210_000, via golden gate assembly using PaqCI; amplifying a Right homology arm fragment ~600 bp in length from pTiC58 for assembly with the corresponding ~600 bp Left homology arm fragment, pAGROm2_112, pAGROm4_210, and pAGRObb_000, yielding pDisarm_C58_112_210_000 [600 HA], via golden gate assembly using PaqCI; amplifying aRight homology arm fragment ~300 bp in length from pTiC58 for assembly with the corresponding ~300 bp Left homology arm fragment, pAGROm2_112, pAGROm4_210, and pAGRObb_000, yielding pDisarm_C58_112_210_000 [300 HA], via golden gate assembly using PaqCI; |
| pTiC58_Right_HA-1200R | CTGTCACCTGCGCTTTCCCTTCAAGAAACGGGTAGTTACCG | strain C58 genomic DNA; strain | PaqCI | Amplifying a Right homology arm fragment ~1,200 bp in length from pTiC58 for assembly with the corresponding ~1,200 bp Left homology arm fragment, pAGROm2_112, pAGROm4_210, and pAGRObb_000, yielding pDisarm_C58_112_210_000, via golden gate assembly using PaqCI; amplifying a Right homology arm fragment from pTi3a for assembly with the corresponding Left homology arm fragment, pAGROm2_112, pAGROm4_210, and pAGRObb_000, yielding pDisarm_3a_112_210_000, via golden gate assembly using PaqCI; amplifying a Right homology arm fragment from pTiC58 for assembly with the corresponding Left homology arm fragment, pAGROm2_002, pAGROm4_010, and pAGRObb_000, yielding pDisarm_C58_002_010_000, via golden gate assembly using PaqCI (used as a control construct without visual reporter cassettes) |
| pTiC58_Right_HA-600R | CTGTCACCTGCGCTTTCCCTGATCGGATACGACTTTACGACA | strain C58 genomic DNA | PaqCI | Amplifying a Right homology arm fragment ~600 bp in length from pTiC58 for assembly with the corresponding ~600 bp Left homology arm fragment, pAGROm2_112, pAGROm4_210, and pAGRObb_000, yielding pDisarm_C58_112_210_000 [600 HA], via golden gate assembly using PaqCI |
| pTiC58_Right_HA-300R | CTGTCACCTGCGCTTTCCCAGATATCTTGCTGCGTTCGGAT | strain C58 genomic DNA | PaqCI | Amplifying a Right homology arm fragment ~300 bp in length from pTiC58 for assembly with the corresponding ~300 bp Left homology arm fragment, pAGROm2_112, pAGROm4_210, and pAGRObb_000, yielding pDisarm_C58_112_210_000 [300 HA], via golden gate assembly using PaqCI |
| pTiAch5_Left_HA-F | CTGTCACCTGCGCTTCGTAAGCCGCATATTGTGATCTGG | strain Ach5 genomic DNA | PaqCI | Amplifying a Left homology arm fragment from pTiAch5 for assembly with the corresponding Right homology arm fragment, pAGROm2_102, pAGROm4_210, and pAGRObb_000, yielding pDisarm_Ach5_102_210_000, via golden gate assembly using PaqCI; amplifying a Left homology arm fragment from pTiAch5 for assembly with the corresponding Right homology arm fragment, pAGROm2_002, pAGROm4_010, and pAGRObb_000, yielding pDisarm_Ach5_002_010_000, via golden gate assembly using PaqCI (used as a control construct without visual reporter cassettes) |
| pTiAch5_Left_HA-R | CTGTCACCTGCGCTTTGATGCTCATTGTGCTCGCAAAC | strain Ach5 genomic DNA | PaqCI | Amplifying a Left homology arm fragment from pTiAch5 for assembly with the corresponding Right homology arm fragment, pAGROm2_102, pAGROm4_210, and pAGRObb_000, yielding pDisarm_Ach5_102_210_000, via golden gate assembly using PaqCI; amplifying a Left homology arm fragment from pTiAch5 for assembly with the corresponding Right homology arm fragment, pAGROm2_002, pAGROm4_010, and pAGRObb_000, yielding pDisarm_Ach5_002_010_000, via golden gate assembly using PaqCI (used as a control construct without visual reporter cassettes) |
| pTiAch5_Right_HA-F | CTGTCACCTGCGCTTCCGCTGGAAGCCCTATCCAAGTTC | strain Ach5 genomic DNA | PaqCI | Amplifying a Right homology arm fragment from pTiAch5 for assembly with the corresponding Left homology arm fragment, pAGROm2_102, pAGROm4_210, and pAGRObb_000, yielding pDisarm_Ach5_102_210_000, via golden gate assembly using PaqCI; amplifying a Right homology arm fragment from pTiAch5 for assembly with the corresponding Left homology arm fragment, pAGROm2_002, pAGROm4_010, and pAGRObb_000, yielding pDisarm_Ach5_002_010_000, via golden gate assembly using PaqCI (used as a control construct without visual reporter cassettes) |
| pTiAch5_Right_HA-R | CTGTCACCTGCGCTTTCCCGACGGATCACTATCAGAACGG | strain Ach5 genomic DNA | PaqCI | Amplifying a Right homology arm fragment from pTiAch5 for assembly with the corresponding Left homology arm fragment, pAGROm2_102, pAGROm4_210, and pAGRObb_000, yielding pDisarm_Ach5_102_210_000, via golden gate assembly using PaqCI; amplifying a Right homology arm fragment from pTiAch5 for assembly with the corresponding Left homology arm fragment, pAGROm2_002, pAGROm4_010, and pAGRObb_000, yielding pDisarm_Ach5_002_010_000, via golden gate assembly using PaqCI (used as a control construct without visual reporter cassettes) |
| pTiAF82/95_Left_HA-F | CTGTCACCTGCGCTTCGTAAGCCGCATATTGTGATCTGG | strain AF82/95 genomic DNA | PaqCI | Amplifying a Left homology arm fragment from pTiAF82/95 for assembly with the corresponding Right homology arm fragment, pAGROm2_114, pAGROm4_210, and pAGRObb_000, yielding pDisarm_AF82/95_114_210_000, via golden gate assembly using PaqCI |
| pTiAF82/95_Left_HA-R | CTGTCACCTGCGCTTTGATGCTCATTGTGCTCGCAAAC | strain AF82/95 genomic DNA | PaqCI | Amplifying a Left homology arm fragment from pTiAF82/95 for assembly with the corresponding Right homology arm fragment, pAGROm2_114, pAGROm4_210, and pAGRObb_000, yielding pDisarm_AF82/95_114_210_000, via golden gate assembly using PaqCI |
| pTiAF82/95_Right_HA-F | CTGTCACCTGCGCTTCCGCTAATGAGATCGTAACCGCGCT | strain AF82/95 genomic DNA | PaqCI | Amplifying a Right homology arm fragment from pTiAF82/95 for assembly with the corresponding Left homology arm fragment, pAGROm2_114, pAGROm4_210, and pAGRObb_000, yielding pDisarm_AF82/95_114_210_000, via golden gate assembly using PaqCI |
| pTiAF82/95_Right_HA-R | CTGTCACCTGCGCTTTCCCACGGCTGACTGGACGATATAT | strain AF82/95 genomic DNA | PaqCI | Amplifying a Right homology arm fragment from pTiAF82/95 for assembly with the corresponding Left homology arm fragment, pAGROm2_114, pAGROm4_210, and pAGRObb_000, yielding pDisarm_AF82/95_114_210_000, via golden gate assembly using PaqCI |
| pTiCM65/95_Left_HA-F | CTGTCACCTGCGCTTCGTATTCCGTCGATCGGATCG | strain CM65/95 genomic DNA (o | PaqCI | Amplifying a Left homology arm fragment from pTiCM65/95 for assembly with the corresponding Right homology arm fragment, pAGROm2_112, pAGROm4_210, and pAGRObb_000, yielding pDisarm_CM65/95_112_210_000, via golden gate assembly using PaqCI |
| pTiCM65/95_Left_HA-R | CTGTCACCTGCGCTTTGATACCACTCGATACAGGCAG | strain CM65/95 genomic DNA (o | PaqCI | Amplifying a Left homology arm fragment from pTiCM65/95 for assembly with the corresponding Right homology arm fragment, pAGROm2_112, pAGROm4_210, and pAGRObb_000, yielding pDisarm_CM65/95_112_210_000, via golden gate assembly using PaqCI |
| pTiCM65/95_Right_HA-F | CTGTCACCTGCGCTTCCGCTTGTACGTCCATATGGTCGG | strain CM65/95 genomic DNA (o | PaqCI | Amplifying a Right homology arm fragment from pTiCM65/95 for assembly with the corresponding Left homology arm fragment, pAGROm2_112, pAGROm4_210, and pAGRObb_000, yielding pDisarm_CM65/95_112_210_000, via golden gate assembly using PaqCI |
| pTiCM65/95_Right_HA-R | CTGTCACCTGCGCTTTCCCAGAATTTCATCCAGGCGTTTGC | strain CM65/95 genomic DNA (o | PaqCI | Amplifying a Right homology arm fragment from pTiCM65/95 for assembly with the corresponding Left homology arm fragment, pAGROm2_112, pAGROm4_210, and pAGRObb_000, yielding pDisarm_CM65/95_112_210_000, via golden gate assembly using PaqCI |
| pTiX1/95_Left_HA-F | CTGTCACCTGCGCTTCGTATTGAATGTTCCGTCGATCGG | strain X1/95 genomic DNA (or st | PaqCI | Amplifying a Left homology arm fragment from pTiX1/95 for assembly with the corresponding Right homology arm fragment, pAGROm2_112, pAGROm4_210, and pAGRObb_000, yielding pDisarm_X1/95_112_210_000, via golden gate assembly using PaqCI |
| pTiX1/95_Left_HA-R | CTGTCACCTGCGCTTTGATCGGGAGAGCCGTTGTAA | strain X1/95 genomic DNA (or st | PaqCI | Amplifying a Left homology arm fragment from pTiX1/95 for assembly with the corresponding Right homology arm fragment, pAGROm2_112, pAGROm4_210, and pAGRObb_000, yielding pDisarm_X1/95_112_210_000, via golden gate assembly using PaqCI |
| pTiX1/95_Right_HA-F | CTGTCACCTGCGCTTCCGCTTGTATGTGCATGCCAACCA | strain X1/95 genomic DNA (or st | PaqCI | Amplifying a Right homology arm fragment from pTiX1/95 for assembly with the corresponding Left homology arm fragment, pAGROm2_112, pAGROm4_210, and pAGRObb_000, yielding pDisarm_X1/95_112_210_000, via golden gate assembly using PaqCI |
| pTiX1/95_Right_HA-R | CTGTCACCTGCGCTTTCCCGAAAGCTTCGACAAGATTCCAC | strain X1/95 genomic DNA (or st | PaqCI | Amplifying a Right homology arm fragment from pTiX1/95 for assembly with the corresponding Left homology arm fragment, pAGROm2_112, pAGROm4_210, and pAGRObb_000, yielding pDisarm_X1/95_112_210_000, via golden gate assembly using PaqCI |
| pTi13-2099-1-2_Left_HA-F | CTGTCACCTGCGCTTCGTAAACCGTGTTGGAACGGTCTT | strain 13-2099-1-2 genomic DN | PaqCI | Amplifying a Left homology arm fragment from pTi13-2099-1-2 for assembly with the corresponding Right homology arm fragment, pAGROm2_112, pAGROm4_210, and pAGRObb_000, yielding pDisarm_13-2099-1-2_112_210_000, via golden gate assembly using PaqCI |
| pTi13-2099-1-2_Left_HA-R | CTGTCACCTGCGCTTTGATCGTTGCTGCCTGTGATCAAA | strain 13-2099-1-2 genomic DN | PaqCI | Amplifying a Left homology arm fragment from pTi13-2099-1-2 for assembly with the corresponding Right homology arm fragment, pAGROm2_112, pAGROm4_210, and pAGRObb_000, yielding pDisarm_13-2099-1-2_112_210_000, via golden gate assembly using PaqCI |
| pTi13-2099-1-2_Right_HA-F | CTGTCACCTGCGCTTCCGCGCAGCATTCCAAAAAGATCC | strain 13-2099-1-2 genomic DN | PaqCI | Amplifying a Right homology arm fragment from pTi13-2099-1-2 for assembly with the corresponding Left homology arm fragment, pAGROm2_112, pAGROm4_210, and pAGRObb_000, yielding pDisarm_13-2099-1-2_112_210_000, via golden gate assembly using PaqCI |
| pTi13-2099-1-2_Right_HA-R | CTGTCACCTGCGCTTTCCCCGTTCCGTTCATCAGTGTTG | strain 13-2099-1-2 genomic DN | PaqCI | Amplifying a Right homology arm fragment from pTi13-2099-1-2 for assembly with the corresponding Left homology arm fragment, pAGROm2_112, pAGROm4_210, and pAGRObb_000, yielding pDisarm_13-2099-1-2_112_210_000, via golden gate assembly using PaqCI |
| pTiM56/79_Left_HA-F | CTGTCACCTGCGCTTCGTACTCGCCACTTTCTTCACCCA | strain M56/79 genomic DNA | PaqCI | Amplifying a Left homology arm fragment from pTiM56/79 for assembly with the corresponding Right homology arm fragment, pAGROm2_111, pAGROm4_210, and pAGRObb_000, yielding pDisarm_M56/79_111_210_000, via golden gate assembly using PaqCI |
| pTiM56/79_Left_HA-R | CTGTCACCTGCGCTTTGATATCACCACTCGATACAGGCAG | strain M56/79 genomic DNA | PaqCI | Amplifying a Left homology arm fragment from pTiM56/79 for assembly with the corresponding Right homology arm fragment, pAGROm2_111, pAGROm4_210, and pAGRObb_000, yielding pDisarm_M56/79_111_210_000, via golden gate assembly using PaqCI |
| pTiM56/79_Right_HA-F | CTGTCACCTGCGCTTCCGCCTTCTGTTCAACCACCCAAACG | strain M56/79 genomic DNA | PaqCI | Amplifying a Right homology arm fragment from pTiM56/79 for assembly with the corresponding Left homology arm fragment, pAGROm2_111, pAGROm4_210, and pAGRObb_000, yielding pDisarm_M56/79_111_210_000, via golden gate assembly using PaqCI |
| pTiM56/79_Right_HA-R | CTGTCACCTGCGCTTTCCCGCACTTGCCAACCTCAACATC | strain M56/79 genomic DNA | PaqCI | Amplifying a Right homology arm fragment from pTiM56/79 for assembly with the corresponding Left homology arm fragment, pAGROm2_111, pAGROm4_210, and pAGRObb_000, yielding pDisarm_M56/79_111_210_000, via golden gate assembly using PaqCI |
| pTiT60/94_Left_HA-F | GTCTGGTCTCTCGCCCCTGTTCCAGTCGATTCAGATAG | strain T60/94 genomic DNA | BsaI | Amplifying a Left homology arm fragment from pTiT60/94 for assembly with the corresponding Right homology arm fragment, a fragment consisting of an sfGFP expression cassette flanked upstream by an FRT site (from pVS-sfGFP-2), a fragment consisting of a spectinomycin resistance marker cassette flanked downstream by an FRT site (from pVS-rec (S)), an eforRed expression cassette fragment (from pVS-eforRed-3), a sacB counterselection marker fragment (from pFlp-Km), an $oriT_{pRK2}$ fragment (from pCGT4464A), and a fragment consisting of a carbenicillin resistance marker and colE1 ori (from pMOD_B2303), yielding pDisarm_T60/94_FRT^sfGFP-specR^FRT_eforRed-sacB_$oriT_{pRK2}$_colE1-carbR, via golden gate assembly using BsaI |
| pTiT60/94_Left_HA-R | GTCTGGTCTCTGCATGTTCGGGTGTCGAGAGATG | strain T60/94 genomic DNA | BsaI | Amplifying a Left homology arm fragment from pTiT60/94 for assembly with the corresponding Right homology arm fragment, a fragment consisting of an sfGFP expression cassette flanked upstream by an FRT site (from pVS-sfGFP-2), a fragment consisting of a spectinomycin resistance marker cassette flanked downstream by an FRT site (from pVS-rec (S)), an eforRed expression cassette fragment (from pVS-eforRed-3), a sacB counterselection marker fragment (from pFlp-Km), an $oriT_{pRK2}$ fragment (from pCGT4464A), and a fragment consisting of a carbenicillin resistance marker and colE1 ori (from pMOD_B2303), yielding pDisarm_T60/94_FRT^sfGFP-specR^FRT_eforRed-sacB_$oriT_{pRK2}$_colE1-carbR, via golden gate assembly using BsaI |
| pTiT60/94_Right_HA-F | GTCTGGTCTCTTCCTTTCTCCGCTACCGGTC | strain T60/94 genomic DNA | BsaI | Amplifying a Right homology arm fragment from pTiT60/94 for assembly with the corresponding Left homology arm fragment, a fragment consisting of an sfGFP expression cassette flanked upstream by an FRT site (from pVS-sfGFP-2), a fragment consisting of a spectinomycin resistance marker cassette flanked downstream by an FRT site (from pVS-rec (S)), an eforRed expression cassette fragment (from pVS-eforRed-3), a sacB counterselection marker fragment (from pFlp-Km), an $oriT_{pRK2}$ fragment (from pCGT4464A), and a fragment consisting of a carbenicillin resistance marker and colE1 ori (from pMOD_B2303), yielding pDisarm_T60/94_FRT^sfGFP-specR^FRT_eforRed-sacB_$oriT_{pRK2}$_colE1-carbR, via golden gate assembly using BsaI |
| pTiT60/94_Right_HA-R | GTCTGGTCTCTCGGTGCTTTCGGCTGGAATCG | strain T60/94 genomic DNA | BsaI | Amplifying a Right homology arm fragment from pTiT60/94 for assembly with the corresponding Left homology arm fragment, a fragment consisting of an sfGFP expression cassette flanked upstream by an FRT site (from pVS-sfGFP-2), a fragment consisting of a spectinomycin resistance marker cassette flanked downstream by an FRT site (from pVS-rec (S)), an eforRed expression cassette fragment (from pVS-eforRed-3), a sacB counterselection marker fragment (from pFlp-Km), an $oriT_{pRK2}$ fragment (from pCGT4464A), and a fragment consisting of a carbenicillin resistance marker and colE1 ori (from pMOD_B2303), yielding pDisarm_T60/94_FRT^sfGFP-specR^FRT_eforRed-sacB_$oriT_{pRK2}$_colE1-carbR, via golden gate assembly using BsaI |
| pTiK224_T1_Left_HA-F | CTGTCACCTGCGCTTCGTAGGTAGGATCTCCACCGAG | strain K224 genomic DNA | PaqCI | Amplifying a Left homology arm fragment from pTiK224 surrounding T-DNA segment 1 for assembly with the corresponding Right homology arm fragment, pAGROm2_112, pAGROm4_210, and pAGRObb_000, yielding pDisarm_K224[T1]_112_210_000, via golden gate assembly using PaqCI |
| pTiK224_T1_Left_HA-R | CTGTCACCTGCGCTTTGATAGGTGCGGTAAGTCGATTTC | strain K224 genomic DNA | PaqCI | Amplifying a Left homology arm fragment from pTiK224 surrounding T-DNA segment 1 for assembly with the corresponding Right homology arm fragment, pAGROm2_112, pAGROm4_210, and pAGRObb_000, yielding pDisarm_K224[T1]_112_210_000, via golden gate assembly using PaqCI |
| pTiK224_T1_Right_HA-F | CTGTCACCTGCGCTTCCGCTTCAAGGGCATCGAGGAC | strain K224 genomic DNA | PaqCI | Amplifying a Right homology arm fragment from pTiK224 surrounding T-DNA segment 1 for assembly with the corresponding Left homology arm fragment, pAGROm2_112, pAGROm4_210, and pAGRObb_000, yielding pDisarm_K224[T1]_112_210_000, via golden gate assembly using PaqCI |
| pTiK224_T1_Right_HA-R | CTGTCACCTGCGCTTTCCCACAGATCTGTTTCCCATCGTG | strain K224 genomic DNA | PaqCI | Amplifying a Right homology arm fragment from pTiK224 surrounding T-DNA segment 1 for assembly with the corresponding Left homology arm fragment, pAGROm2_112, pAGROm4_210, and pAGRObb_000, yielding pDisarm_K224[T1]_112_210_000, via golden gate assembly using PaqCI |
| pTiK224_T2_Left_HA-F | CTGTCACCTGCGCTTCGTAGGCTGGCAAACGGATCATAA | strain K224 genomic DNA | PaqCI | Amplifying a Left homology arm fragment from pTiK224 surrounding T-DNA segment 2 for assembly with the corresponding Right homology arm fragment, pAGROm2_122, pAGROm4_210, and pAGRObb_000, yielding pDisarm_K224[T2]_122_210_000, via golden gate assembly using PaqCI |
| pTiK224_T2_Left_HA-R | CTGTCACCTGCGCTTTGATGTCGCTATTCCCGCTAACT | strain K224 genomic DNA | PaqCI | Amplifying a Left homology arm fragment from pTiK224 surrounding T-DNA segment 2 for assembly with the corresponding Right homology arm fragment, pAGROm2_122, pAGROm4_210, and pAGRObb_000, yielding pDisarm_K224[T2]_122_210_000, via golden gate assembly using PaqCI |
| pTiK224_T2_Right_HA-F | CTGTCACCTGCGCTTCCGCCTGCAATCTCCCAGGCAT | strain K224 genomic DNA | PaqCI | Amplifying a Right homology arm fragment from pTiK224 surrounding T-DNA segment 2 for assembly with the corresponding Left homology arm fragment, pAGROm2_122, pAGROm4_210, and pAGRObb_000, yielding pDisarm_K224[T2]_122_210_000, via golden gate assembly using PaqCI |
| pTiK224_T2_Right_HA-R | CTGTCACCTGCGCTTTCCCTCGGGTCAAACTTCTTTGGT | strain K224 genomic DNA | PaqCI | Amplifying a Right homology arm fragment from pTiK224 surrounding T-DNA segment 2 for assembly with the corresponding Left homology arm fragment, pAGROm2_122, pAGROm4_210, and pAGRObb_000, yielding pDisarm_K224[T2]_122_210_000, via golden gate assembly using PaqCI |
| C58_recF_Left_HA-F | CTGTCACCTGCGCTTCGTAGCCTGCAGCTTGGCGATA | strain C58 (or AGL1) genomic DN | PaqCI | Amplifying a Left homology arm fragment from the C58 circular chromosome adjacent to the *recF* locus for assembly with the corresponding Right homology arm fragment, pAGROm2_112, pAGROm4_211, and pAGRObb_000, yielding pDelete_C58_recF_112_211_000 (containing *recA* expression cassette), via golden gate assembly using PaqCI; amplifying a Left homology arm fragment from the C58 circular chromosome adjacent to the *recF* locus for assembly with the corresponding Right homology arm fragment, pAGROm2_112, pAGROm4_210, and pAGRObb_000, yielding pDelete_C58_recF_112_210_000 (the control construct lacking a *recA* expression cassette), via golden gate assembly using PaqCI |

| | | | | |
|---|---|---|---|---|
| C58_recF_Left_HA-R | CTGTCACCTGCGCTTTGATGCATCAAGCACAAGTGCGG | strain C58 (or AGL1) genomic DN | PaqCI | Amplifying a Left homology arm fragment from the C58 circular chromosome adjacent to the *recF* locus for assembly with the corresponding Right homology arm fragment, pAGROm2_112, pAGROm4_211, and pAGRObb_000, yielding pDelete_C58_recF_112_211_000 (containing *recA* expression cassette), via golden gate assembly using PaqCI; amplifying a Left homology arm fragment from the C58 circular chromosome adjacent to the *recF* locus for assembly with the corresponding Right homology arm fragment, pAGROm2_112, pAGROm4_210, and pAGRObb_000, yielding pDelete_C58_recF_112_210_000 (the control construct lacking a *recA* expression cassette), via golden gate assembly using PaqCI |
| C58_recF_Right_HA-F | CTGTCACCTGCGCTTCCGCTGGATGACAGGCTGAACC | strain C58 (or AGL1) genomic DN | PaqCI | Amplifying a Right homology arm fragment from the C58 circular chromosome adjacent to the *recF* locus for assembly with the corresponding Left homology arm fragment, pAGROm2_112, pAGROm4_211, and pAGRObb_000, yielding pDelete_C58_recF_112_211_000 (containing *recA* expression cassette), via golden gate assembly using PaqCI; amplifying a Righthomology arm fragment from the C58 circular chromosome adjacent to the *recF* locus for assembly with the corresponding Left homology arm fragment, pAGROm2_112, pAGROm4_210, and pAGRObb_000, yielding pDelete_C58_recF_112_210_000 (the control construct lacking a *recA* expression cassette), via golden gate assembly using PaqCI |
| C58_recF_Right_HA-R | CTGTCACCTGCGCTTTCCCCCGCTGGAGATTGGAGAG | strain C58 (or AGL1) genomic DN | PaqCI | Amplifying a Right homology arm fragment from the C58 circular chromosome adjacent to the *recF* locus for assembly with the corresponding Left homology arm fragment, pAGROm2_112, pAGROm4_211, and pAGRObb_000, yielding pDelete_C58_recF_112_211_000 (containing *recA* expression cassette), via golden gate assembly using PaqCI; amplifying a Righthomology arm fragment from the C58 circular chromosome adjacent to the *recF* locus for assembly with the corresponding Left homology arm fragment, pAGROm2_112, pAGROm4_210, and pAGRObb_000, yielding pDelete_C58_recF_112_210_000 (the control construct lacking a *recA* expression cassette), via golden gate assembly using PaqCI |
| C58DE_recA_blaTEM-F | CTGTCACCTGCGCTTCGTATCCGTATCATAATTGTCGTGCTT | strain AGL1 genomic DNA | PaqCI | Amplifying a fragment from the circular chromosome in the AGL1 strain containing the blaTEM ampicillin resistance marker at the inactive *recA* locus flanked on either side by sequence segments homologous to the wild-type *recA* locus in the C58 chromosomal background. This fragment was assembled with pAGROm4_210 and pAGRObb_000, yielding pDelete_C58_recA_blaTEM_210_000, via golden gate assembly using PaqCI |
| C58DE_recA_blaTEM-R | CTGTCACCTGCGCTTTCCCCGTTCATGAAGAGGATGTCGA | strain AGL1 genomic DNA | PaqCI | Amplifying a fragment from the circular chromosome in the AGL1 strain containing the blaTEM ampicillin resistance marker at the inactive *recA* locus flanked on either side by sequence segments homologous to the wild-type *recA* locus in the C58 chromosomal background. This fragment was assembled with pAGROm4_210 and pAGRObb_000, yielding pDelete_C58_recA_blaTEM_210_000, via golden gate assembly using PaqCI |
| C58DE_thyA_Left_HA-F | CTGTCACCTGCGCTTCGTAAGATTTCGTCCTTCGGACG | strain C58 (or AGL1) genomic DN | PaqCI | Amplifying a Left homology arm fragment from the C58 circular chromosome adjacent to the *thyA* locus for assembly with the corresponding Right homology arm fragment, pAGROm2_132, pAGROm4_211, and pAGRObb_000, yielding pDelete_C58_thyA_132_211_000, via golden gate assembly using PaqCI |
| C58DE_thyA_Left_HA-R | CTGTCACCTGCGCTTTGATTTCAAGCATCAAGGCACCG | strain C58 (or AGL1) genomic DN | PaqCI | Amplifying a Left homology arm fragment from the C58 circular chromosome adjacent to the *thyA* locus for assembly with the corresponding Right homology arm fragment, pAGROm2_132, pAGROm4_211, and pAGRObb_000, yielding pDelete_C58_thyA_132_211_000, via golden gate assembly using PaqCI |
| C58DE_thyA_Right_HA-F | CTGTCACCTGCGCTTCCGCCCGGAGAAGATCGAGATATTGTTT | strain C58 (or AGL1) genomic DN | PaqCI | Amplifying a Right homology arm fragment from the C58 circular chromosome adjacent to the *thyA* locus for assembly with the corresponding Left homology arm fragment, pAGROm2_132, pAGROm4_211, and pAGRObb_000, yielding pDelete_C58_thyA_132_211_000, via golden gate assembly using PaqCI |
| C58DE_thyA_Right_HA-R | CTGTCACCTGCGCTTTCCCGAAAAGCCGATGCCGTTT | strain C58 (or AGL1) genomic DN | PaqCI | Amplifying a Right homology arm fragment from the C58 circular chromosome adjacent to the *thyA* locus for assembly with the corresponding Left homology arm fragment, pAGROm2_132, pAGROm4_211, and pAGRObb_000, yielding pDelete_C58_thyA_132_211_000, via golden gate assembly using PaqCI |
| PstDC3000_5060_Left_HA-F | CTGTCACCTGCGCTTCGTAGTGCCTACAAGCCCAAAGA | *Pseudomonas syringae* pv. *toma* | PaqCI | Amplifying a Left homology arm fragment from the DC3000 chromosome adjacent to the operon containing ORFs 5060-5064 for assembly with the corresponding Right homology arm fragment, pAGROm2_101, pAGROm4_210, and pAGRObb_000, yielding pDelete_PstDC3000_[5060-64]_101_210_000, via golden gate assembly using PaqCI |
| PstDC3000_5060_Left_HA-R | CTGTCACCTGCGCTTTGATCTTCGGTCACGCTGGTG | *Pseudomonas syringae* pv. *toma* | PaqCI | Amplifying a Left homology arm fragment from the DC3000 chromosome adjacent to the operon containing ORFs 5060-5064 for assembly with the corresponding Right homology arm fragment, pAGROm2_101, pAGROm4_210, and pAGRObb_000, yielding pDelete_PstDC3000_[5060-64]_101_210_000, via golden gate assembly using PaqCI |
| PstDC3000_5064_Right_HA-F | CTGTCACCTGCGCTTCCGCCGCCATCAAGCGGTTTATCT | *Pseudomonas syringae* pv. *toma* | PaqCI | Amplifying a Right homology arm fragment from the DC3000 chromosome adjacent to the operon containing ORFs 5060-5064 for assembly with the corresponding Left homology arm fragment, pAGROm2_101, pAGROm4_210, and pAGRObb_000, yielding pDelete_PstDC3000_[5060-64]_101_210_000, via golden gate assembly using PaqCI |
| PstDC3000_5064_Right_HA-R | CTGTCACCTGCGCTTTCCCGCAACCACGACAACCCTT | *Pseudomonas syringae* pv. *toma* | PaqCI | Amplifying a Right homology arm fragment from the DC3000 chromosome adjacent to the operon containing ORFs 5060-5064 for assembly with the corresponding Left homology arm fragment, pAGROm2_101, pAGROm4_210, and pAGRObb_000, yielding pDelete_PstDC3000_[5060-64]_101_210_000, via golden gate assembly using PaqCI |

| Wild-type strain name | Modification construct used | Selection type / concentration (in µg/mL) used in target strain | Method of construct introduction | Excision plasmid used |
|---|---|---|---|---|
| 3a | pDisarm_3a_112_210_000 | spectinomycin 200 | Electroporation | pAGROex_211 |
| X1/95 | pDisarm_X1/95_112_210_000 | spectinomycin 200 | Electroporation | pAGROex_211 |
| O54/95 | pDisarm_X1/95_112_210_000 | spectinomycin 200 | Electroporation | pAGROex_211 |
| 13-2099-1-2 | pDisarm_13-2099-1-2_112_210_000 | spectinomycin 200 | Electroporation | pAGROex_211 |
| M56/79 | pDisarm_M56/79 _111_210_000 | kanamycin 100 | Electroporation | pAGROex_214 |
| CM65/95 | pDisarm_CM65/95_112_210_000 | spectinomycin 200 | Tri-parental mating | pAGROex_211 |
| N33/94 | pDisarm_CM65/95_112_210_000 | spectinomycin 200 | Tri-parental mating | pAGROex_214 |
| AF82/95 | pDisarm_AF82/95_114_210_000 | apramycin 100 | Tri-parental mating | pAGROex_212 |
| K224 | pDisarm_K224[T1]_112_210_000 and pDisarm_K224[T2]_122_210_000 | spectinomycin 200 | Tri-parental mating | pAGROex_214 |
| T60/94 | pDisarm_T60/94_102_210_000 | spectinomycin 200 | Tri-parental mating | pAGROex_201 |
| C58_DER | pDelete_C58_thyA_132_211_000 | spectinomycin 200 | Tri-parental mating | pAGROex_214 |
| AGL1 (C58) | pDelete_C58_recF_112_211_000 | spectinomycin 200 | Electroporation | pAGROex_211 |

| Well positions | Sample abbreviation | Strain background | Expression cassette | $OD_{600}$ absorbance reading | Fluorescence intensity (sfGFP) | Ratio of fluorescence intensity to cell density |
|---|---|---|---|---|---|---|
| A1 | AG-B | AGL1 | (empty) | 0.098364 | 45 | 457.4844455 |
| A2 | AG-B | AGL1 | (empty) | 0.097105 | 44 | 453.1177591 |
| A3 | AG-B | AGL1 | (empty) | 0.097393 | 45 | 462.0455269 |
| A4 | CM-B | CM65/95 | (empty) | 0.085899 | 47 | 547.154216 |
| A5 | CM-B | CM65/95 | (empty) | 0.085714 | 47 | 548.3351611 |
| A6 | CM-B | CM65/95 | (empty) | 0.085866 | 46 | 535.718445 |
| A7 | T-B | T268/95 | (empty) | 0.12917 | 48 | 371.6033135 |
| A8 | T-B | T268/95 | (empty) | 0.10721 | 48 | 447.7194292 |
| A9 | T-B | T268/95 | (empty) | 0.096529 | 47 | 486.9003098 |
| B1 | AG-0 | AGL1 | PglpT::sRBS-sfGFP::rrnB_T1 | 0.092287 | 50 | 541.7881175 |
| B2 | AG-0 | AGL1 | PglpT::sRBS-sfGFP::rrnB_T1 | 0.093096 | 50 | 537.0800034 |
| B3 | AG-0 | AGL1 | PglpT::sRBS-sfGFP::rrnB_T1 | 0.093033 | 50 | 537.4437028 |
| B4 | CM-0 | CM65/95 | PglpT::sRBS-sfGFP::rrnB_T1 | 0.093302 | 53 | 568.0478446 |
| B5 | CM-0 | CM65/95 | PglpT::sRBS-sfGFP::rrnB_T1 | 0.092568 | 51 | 550.9463313 |
| B6 | CM-0 | CM65/95 | PglpT::sRBS-sfGFP::rrnB_T1 | 0.093223 | 52 | 557.8022591 |
| B7 | T-0 | T268/95 | PglpT::sRBS-sfGFP::rrnB_T1 | 0.093971 | 48 | 510.7958838 |
| B8 | T-0 | T268/95 | PglpT::sRBS-sfGFP::rrnB_T1 | 0.094002 | 48 | 510.6274335 |
| B9 | T-0 | T268/95 | PglpT::sRBS-sfGFP::rrnB_T1 | 0.25627 | 52 | 202.9109923 |
| C1 | AG-1 | AGL1 | PJ23100::AtSD-sfGFP::T3T | 0.088691 | 611 | 6889.08683 |
| C2 | AG-1 | AGL1 | PJ23100::AtSD-sfGFP::T3T | 0.089022 | 610 | 6852.238772 |
| C3 | AG-1 | AGL1 | PJ23100::AtSD-sfGFP::T3T | 0.088869 | 611 | 6875.288346 |
| C4 | CM-1 | CM65/95 | PJ23100::AtSD-sfGFP::T3T | 0.094171 | 347 | 3684.786187 |
| C5 | CM-1 | CM65/95 | PJ23100::AtSD-sfGFP::T3T | 0.089767 | 347 | 3865.563069 |
| C6 | CM-1 | CM65/95 | PJ23100::AtSD-sfGFP::T3T | 0.090561 | 353 | 3897.925155 |
| C7 | T-1 | T268/95 | PJ23100::AtSD-sfGFP::T3T | 0.56136 | 1340 | 2387.059997 |
| C8 | T-1 | T268/95 | PJ23100::AtSD-sfGFP::T3T | 0.34727 | 1190 | 3426.728482 |
| C9 | T-1 | T268/95 | PJ23100::AtSD-sfGFP::T3T | 0.31323 | 863 | 2755.163937 |
| D1 | AG-2 | AGL1 | PJ23119::AtSD-sfGFP::T3T | 0.088763 | 1351 | 15220.30576 |
| D2 | AG-2 | AGL1 | PJ23119::AtSD-sfGFP::T3T | 0.089078 | 1346 | 15110.35272 |
| D3 | AG-2 | AGL1 | PJ23119::AtSD-sfGFP::T3T | 0.0901 | 1345 | 14927.85794 |
| D4 | CM-2 | CM65/95 | PJ23119::AtSD-sfGFP::T3T | 0.10212 | 610 | 5973.364669 |
| D5 | CM-2 | CM65/95 | PJ23119::AtSD-sfGFP::T3T | 0.10111 | 602 | 5953.911581 |
| D6 | CM-2 | CM65/95 | PJ23119::AtSD-sfGFP::T3T | 0.10316 | 601 | 5825.901512 |
| D7 | T-2 | T268/95 | PJ23119::AtSD-sfGFP::T3T | 0.089138 | 685 | 7684.71359 |
| D8 | T-2 | T268/95 | PJ23119::AtSD-sfGFP::T3T | 0.094663 | 686 | 7246.759558 |
| D9 | T-2 | T268/95 | PJ23119::AtSD-sfGFP::T3T | 0.089302 | 656 | 7345.860115 |
| E1 | AG-3 | AGL1 | Pint::AtSD-sfGFP::T3T | 0.08901 | 1182 | 13279.40681 |
| E2 | AG-3 | AGL1 | Pint::AtSD-sfGFP::T3T | 0.089564 | 1179 | 13163.77116 |
| E3 | AG-3 | AGL1 | Pint::AtSD-sfGFP::T3T | 0.089658 | 1192 | 13294.96531 |
| E4 | CM-3 | CM65/95 | Pint::AtSD-sfGFP::T3T | 0.09925 | 898 | 9047.858942 |
| E5 | CM-3 | CM65/95 | Pint::AtSD-sfGFP::T3T | 0.098247 | 884 | 8997.730211 |
| E6 | CM-3 | CM65/95 | Pint::AtSD-sfGFP::T3T | 0.098276 | 871 | 8862.794579 |
| E7 | T-3 | T268/95 | Pint::AtSD-sfGFP::T3T | 0.098315 | 1185 | 12053.09464 |
| E8 | T-3 | T268/95 | Pint::AtSD-sfGFP::T3T | 0.098703 | 1202 | 12177.94799 |
| E9 | T-3 | T268/95 | Pint::AtSD-sfGFP::T3T | 0.094419 | 1266 | 13408.31824 |

| | | | | | | |
|---|---|---|---|---|---|---|
| F1 | AG-4 | AGL1 | Atu0824p::AtSD-sfGFP::T3T | 0.093212 | 694 | 7445.393297 |
| F2 | AG-4 | AGL1 | Atu0824p::AtSD-sfGFP::T3T | 0.096906 | 686 | 7079.025035 |
| F3 | AG-4 | AGL1 | Atu0824p::AtSD-sfGFP::T3T | 0.093116 | 692 | 7431.590704 |
| F4 | CM-4 | CM65/95 | Atu0824p::AtSD-sfGFP::T3T | 0.10158 | 551 | 5424.296121 |
| F5 | CM-4 | CM65/95 | Atu0824p::AtSD-sfGFP::T3T | 0.1007 | 550 | 5461.767627 |
| F6 | CM-4 | CM65/95 | Atu0824p::AtSD-sfGFP::T3T | 0.10036 | 562 | 5599.840574 |
| F7 | T-4 | T268/95 | Atu0824p::AtSD-sfGFP::T3T | 0.089771 | 474 | 5280.101592 |
| F8 | T-4 | T268/95 | Atu0824p::AtSD-sfGFP::T3T | 0.094681 | 480 | 5069.654947 |
| F9 | T-4 | T268/95 | Atu0824p::AtSD-sfGFP::T3T | 0.097022 | 602 | 6204.778298 |

| Primer name | Sequence | Template | Enzyme ends | Purpose |
|---|---|---|---|---|
| pTiC58_Left_flanking-F | CGAGAGATCGATCCTGTCC | strain C58_DC or C58_DE genor | N/A | Colony PCR used to verify secondary recombination and marker excision status in disarmed derivatives of C58 or 3a |
| pTiC58_Right_flanking-R | TTCCCTTTGGTCTGATCCC | strain C58_DC or C58_DE genor | N/A | Colony PCR used to verify secondary recombination and marker excision status in disarmed derivatives of C58 or 3a |
| pTiAch5_Left_flanking-F | AATGACGTCACCTCATCGG | strain Ach5_DC or Ach5_DE gen | N/A | Colony PCR used to verify secondary recombination and marker excision status in disarmed derivatives of Ach5 |
| pTiAch5_Right_flanking-R | CGACTGGTTTCGACAGTCAT | strain Ach5_DC or Ach5_DE gen | N/A | Colony PCR used to verify secondary recombination and marker excision status in disarmed derivatives of Ach5 |
| pTiAF82/95_Left_flanking-F | CGATTATGCTTAAAGCGAAGCGTTA | strain AF82/95_DC or AF82/95_ | N/A | Colony PCR used to verify secondary recombination and marker excision status in disarmed derivatives of AF82/95 |
| pTiAF82/95_Right_flanking-R | CGTGAAACTGTGAATTCCGG | strain AF82/95_DC or AF82/95_ | N/A | Colony PCR used to verify secondary recombination and marker excision status in disarmed derivatives of AF82/95 |
| pTiCM65/95_Left_flanking-F | AATCAGATCTCGAACAGGTGC | strain CM65/95_DC or CM65/95 | N/A | Colony PCR used to verify secondary recombination and marker excision status in disarmed derivatives of CM65/95 and N33/94 |
| pTiCM65/95_Right_flanking-R | AAAGGTCACCGGTGTGATTAC | strain CM65/95_DC or CM65/95 | N/A | Colony PCR used to verify secondary recombination and marker excision status in disarmed derivatives of CM65/95 and N33/94 |
| pTiX1/95_Left_flanking-F | CAATCGAGATCGCCCAATTC | strain X1/95_DC or X1/95_DE ge | N/A | Colony PCR used to verify secondary recombination and marker excision status in disarmed derivatives of X1/95 and O54/95 |
| pTiX1/95_Right_flanking-R | CTGGCCGGCTACATTGAA | strain X1/95_DC or X1/95_DE ge | N/A | Colony PCR used to verify secondary recombination and marker excision status in disarmed derivatives of X1/95 and O54/95 |
| pTi13-2099-1-2_Left_flanking-F | GCTGATCTTCAGATCATCCG | strain 13-2099-1-2_DC or 13-20 | N/A | Colony PCR used to verify secondary recombination and marker excision status in disarmed derivatives of 13-2099-1-2 |
| pTi13-2099-1-2_Right_flanking-R | CGCTCAAGAAACGGGTAGTT | strain 13-2099-1-2_DC or 13-20 | N/A | Colony PCR used to verify secondary recombination and marker excision status in disarmed derivatives of 13-2099-1-2 |
| pTiM56/79_Left_flanking-F | GTACAGACCAATCGAGATCGCC | strain M56/79_DC or M56/79_D | N/A | Colony PCR used to verify secondary recombination and marker excision status in disarmed derivatives of M56/79 |
| pTiM56/79_Right_flanking-R | CGGCAGCCTATCAAACCAATAAAG | strain M56/79_DC or M56/79_D | N/A | Colony PCR used to verify secondary recombination and marker excision status in disarmed derivatives of M56/79 |
| pTiT60/94_Left_flanking-F | CAATGACCAAGCCTATGCCTT | strain T60/94_DC or T60/94_DE | N/A | Colony PCR used to verify secondary recombination and marker excision status in disarmed derivatives of T60/94 |
| pTiT60/94_Right_flanking-R | GCTCTCATGGTTTCTTCCTTTG | strain T60/94_DC or T60/94_DE | N/A | Colony PCR used to verify secondary recombination and marker excision status in disarmed derivatives of T60/94 |
| pTiK224_T1_Left_flanking-F | CCGCGCGAAACGGATATT | strainK224_1DC or K224_1DE ge | N/A | Colony PCR used to verify secondary recombination and marker excision status in T-DNA 1 deletion derivatives of K224 |
| pTiK224_T1_Right_flanking-R | TTCGTCAACTTCAAAGAGCTCT | strainK224_1DC or K224_1DE ge | N/A | Colony PCR used to verify secondary recombination and marker excision status in T-DNA 1 deletion derivatives of K224 |
| pTiK224_T2_Left_flanking-F | GCAATCGTCGCTATACGGA | strain K224_1DE_2DC or K224_1 | N/A | Colony PCR used to verify secondary recombination and marker excision status in T-DNA 2 deletion derivatives of K224 |
| pTiK224_T2_Right_flanking-R | GTCTATCGGCGCGATCTT | strain K224_1DE_2DC or K224_1 | N/A | Colony PCR used to verify secondary recombination and marker excision status in T-DNA 2 deletion derivatives of K224 |
| C58_recF_Left_flanking-F | ACCACCTCGATCTGCTTG | strain AGL1 recF- [MR] or AGL1 r | N/A | Colony PCR used to verify secondary recombination and marker excision status in *recF* knockout derivatives of AGL1 |
| C58_recF_Right_flanking-R | ACATAAGCGTGCGGACAT | strain AGL1 recF- [MR] or AGL1 r | N/A | Colony PCR used to verify secondary recombination and marker excision status in *recF* knockout derivatives of AGL1 |
| C58DE_recA_Left_flanking-F | AATCAGGAAGGCGGATGAA | strain C58_DE recA- [blaTEM] (C | N/A | Colony PCR used to verify secondary recombination status in *recA* knockout (and ampicillin/carbenicillin-resistant) derivatives of C58 |
| C58DE_recA_blaTEM_int-R | CTGCAGCAATGGCAACAA | blaTEM (amplicillin resistance m | N/A | Colony PCR used to verify secondary recombination status in *recA* knockout (and ampicillin/carbenicillin-resistant) derivatives of C58 |
| C58DE_thyA_Left_flanking-F | AGATGGAAATGCAGGCCAG | strain C58_DER thyA- [MR] or C5 | N/A | Colony PCR used to verify secondary recombination and marker excision status in *thyA* knockout derivatives of C58 |
| C58DE_thyA_Right_flanking-R | AGTATACGCAATGTCCCGAAC | strain C58_DER thyA- [MR] or C5 | N/A | Colony PCR used to verify secondary recombination and marker excision status in *thyA* knockout derivatives of C58 |
| PstDC3000_5060_flanking-F | GCCTGTGCACCATCTATGAA | DC3000 [5060_KO] genomic DN | N/A | Colony PCR used to verify secondary recombination and marker excision status in ORF 5060 deletion derivatives of *Pseudomonas syringae* pv. *tomato* DC3000 |
| PstDC3000_5060_flanking-R | GCAGTTCGGCAATTCTGTTG | DC3000 [5060_KO] genomic DN | N/A | Colony PCR used to verify secondary recombination and marker excision status in ORF 5060 deletion derivatives of *Pseudomonas syringae* pv. *tomato* DC3000 |

>pT200-GFP-HygR
tggcaggatatattgtggtgtaaacaaattgacgcttagacaacttaatacacattgcggacgtttta
atgtactgaattaacgccgaattgagtgacttgaagtacactcggtaccgccggcgatcaaacactgagc
tcgatatcgaattcgagctcgctagcgtcgacactagtccatggctgatcttatctttctgagcctgcag
ggatctagtaacatagatgacaccgcgcgcgataatttatcctagtttgcgcgctatattttgttttcta
tcgcgtattaaatgtataattgcgggactctaatcataaaaacccatctcataaataacgtcatgcatta
catgttaattattacatgcttaacgtaattcaacagaaattatatgataatcatcgcaagaccggcaaca
ggattcaatcttaagaaactttattgccaaatgtttgaacgatcggggaaattcgggggatcagcttgtg
atcatgttacatcacctccaagatcccggacgagtgctggggcgtcggtttccactatcggcgagtactt
ctacacagccatcggtccagacggccgcgcttctgcgggcgatttgtgtacgcccgacagtcccggctcc
ggatcggacgattgcgtcgcatcgaccctgcgcccaagctgcatcatcgaaattgccgtcaaccaagctc
tgatagagttggtcaagaccaatgcggagcatatacgcccggagccgcggcgatcctgcaagctccggat
gcctccgctcgaagtagcgcgtctgctgctccatacaagccaaccacggcctccagaagaagatgttggc
gacctcgtattgggaatccccgaacatcgcctcgctccagtcaatgaccgctgttatgcggccattgtcc
gtcaggacattgttggagccgaaatccgcgtgcacgaggtgccggacttcggggcagtcctcggcccaaa
gcatcagctcatcgagagcctgcgcgacggacgcactgacggtgtcgtccatcacagtttgccagtgata
cacatggggatcagcaatcgcgcatatgaaatcacgccatgtagtgtattgaccgattccttgcggtccg
aatgggccgaacccgctcgtctggctaagatcggccgcagcgatcgcatccatggcctccgcgaccggct
gcagaacagcgggcagttcggtttcaggcaggtcttgcaacgtgacaccctgtgcacggcgggagatgca
ataggtcaggctctcgctgaattccccaatgtcaagcacttccggaatcgggagcgcggccgatgcaaag
tgccgataaacataacgatctttgtagaaaccatcggcgcagctatttacccgcaggacatatccacgcc
ctcctacatcgaagctgaaagcacgagattcttcgccctccgagagctgcatcaggtcggagacgctgtc
gaacttttcgatcagaaacttctcgacagacgtcgcggtgagttcaggctttttcatatctcattgcccc
ccgggatcttgcaggtcgagatcctctagcggatccggtgcagattatttggattgagagtgaatatgag
actctaattggataccgaggggaatttatggaacgtcagtggagcatttttgacaagaaatatttgctag
ctgatagtgaccttaggcgacttttgaacgcgcaataatggtttctgacgtatgtgcttagctcattaaa
ctccagaaacccgcggctgagtggctccttcaacgttgcggttctgtcagttccaaacgtaaaacggctt
gtcccgcgtcatcggcgggggtcataacgtgactcccttaattctccgctcatgatcagattgtcgtttc
ccgcctggcgcgccccggtagagctcaattcgcgcgcgatctgcgatcgcattacctgcaggatttggcg
cgccgtccgtcaccagagctccactattttcagaagaagttcccaatagtagtccaaaatttttgtaacg
aagggagcataatagttacatgcaaaggaaaactgccattctttagaggggatgcttgtttaagaacaaa
aaatatatcactttcttttgttccaagtcattgcgtatttttttaaaaatatttgttccttcgtatattt
cgagcttcaatcactttatggttctttgtattctggctttgctgtaaatcgtagctaaccttcttcctag
cagaaattattaatacttgggatatttttttagaatcaagtaaattacatattaccaccacatcgagctg
cttttaaattcatattacagccatataggcttgattcattttgcaaaatttccaggatattgacaacgtt
aacgtcactggattttggttttaggaattagaaattttattgatagaagtattttacaaatacaaataca
tactaagggtttcttatatgctcaacacatgagcgaaaccctataagaaccctaattcccttatctggga
actactcacacattattctggagaaaaatagagagagatagatttgtagagagagactggtgatttttgc
ggacgtcgactcacttgtacagctcgtccatgccgagagtgatcccggcggcggtcacgaactccagcag
gaccatgtgatcgcgcttctcgttggggtctttgctcagggcggactgggtgctcaggtagtggttgtcg
ggcagcagcacggggccgtcgccgatgggggtgttctgctggtagtggtcggcgagctgcacgctgccgt
cctcgatgttgtggcggatcttgaagttcaccttgatgccgttcttctgcttgtcggccatgatatagac
gttgtggctgttgtagttgtactccagcttgtgccccaggatgttgccgtcctccttgaagtcgatgccc
ttcagctcgatgcggttcaccagggtgtcgccctcgaacttcacctcggcgcgggtcttgtagttgccgt
cgtccttgaagaagatggtgcgctcctggacgtagccttcgggcatggcggacttgaagaagtcgtgctg
cttcatgtggtcggggtagcggctgaagcactgcacgccgtaggtcagggtggtcacgagggtgggccag
ggcacgggcagcttgccggtggtgcagatgaacttcagggtcagcttgccgtaggtggcatcgccctcgc
cctcgccggacacgctgaacttgtggccgtttacgtcgccgtccagctcgaccaggatgggcaccacccc
ggtgaacagctcctcgcccttgctcaccatttttcttctacttttagtgttcctgcaggagtcccccgtg
ttctctccaaatgaaatgaacttccttatatagaggaagggtcttgcgaaggatagtgggattgtgcgtc
atcccttacgtcagtggagatatcacatcaatccacttgctttgaagacgtggttggaacgtcttctttt

tccacgatgctcctcgtgggtggggtccatctttggaccactgtcggcagaggcatcttcaacgatgg
cctttcctttatcgcaatgatggcatttgtaggagccaccttccttttccactatcttcacaataaagtg
acagatagctgggcaatggaatccgaggaggtttccggatattaccctttgttgaaaagtctcaattgcc
ctttggtcttctgagactgtatctttgatatttttggagtagacaagtgtgtcgtgctccaccatgttga
cgaagattttcttcttgtcattgagtcgtaagagactctgtatgaactgttcgccagtctttacggcgag
ttctgttaggtcctctatttgaatctttgactccatggcctttgattcagtgggaactaccttttttagag
actccaatctctattacttgccttggtttgtgaagcaagccttgaatcgtccatactggaatagtacttc
tgatcttgagaaatatatctttctctgtgttcttgatgcagttagtcctgaatcttttgactgcatcttt
aaccttcttgggaaggtatttgatttcctggagattattgctcgggtagatcgtcttgatgatacctgct
gcgtaagcctctctaaccatctgtgggttagcattctttctgaaattgaaaaggcaaatctggcgcgccg
atcggtaccgagggtcgtacgaaaaactatcagtgtttgacaggatatattggcgggtaaacctaagaga
aaagagcgtttattagaataatcggatatttaaaagggcgtgaaaaggtttatccgttcgtccatttgta
tgtgcatgccaaccacagggttcccctcgggatcaaagtactttgatccaacccctccgctgctatagtg
cagtcggcttctgacgttcagtgcagccgtcttctgaaaacgacatgtcgcacaagtcctaagttacgcg
acaggctgccgccctgccctttcctggcgttttcttgtcgcgtgttttagtcgcataaagtagaatact
tgcgactagaaccggagacattacgccatgaacaagagcgccgccgctggcctgctgggctatgcccgcg
tcagcaccgacgaccaggacttgaccaaccaacgggccgaactgcacgcggccggctgcaccaagctgtt
ttccgagaagatcaccggcaccaggcgcgaccgcccggagctggccaggatgcttgaccacctacgccct
ggcgacgttgtgacagtgaccaggctagaccgcctggcccgcagcacccgcgacctactggacattgccg
agcgcatccaggaggccggcgcgggcctgcgtagcctggcagagccgtgggccgacaccaccacgccggc
cggccgcatggtgttgaccgtgttcgccggcattgccgagttcgagcgttccctaatcatcgaccgcacc
cggagcgggcgcgaggccgccaaggcccgaggcgtgaagtttggcccccgccctaccctcaccccggcac
agatcgcgcacgcccgcgagctgatcgaccaggaaggccgcaccgtgaaagaggcggctgcactgcttgg
cgtgcatcgctcgaccctgtaccgcgcacttgagcgcagcgaggaagtgacgcccaccgaggccaggcgg
cgcggtgccttccgtgaggacgcattgaccgaggccgacgccctggcggccgccgagaatgaacgccaag
aggaacaagcatgaaaccgcaccaggacggccaggacgaaccgtttttcattaccgaagagatcgaggcg
gagatgatcgcggccgggtacgtgttcgagccgcccgcgcacggctcaaccgtgcggctgcatgaaatcc
tggccggtttgtctgatgccaagctggcggcctggccggccagcttggccgctgaagaaaccgagcgccg
ccgtctaaaaaggtgatgtgtatttgagtaaaacagcttgcgtcatgcggtcgctgcgtatatgatgcga
tgagtaaataaacaaatacgcaaggggaacgcatgaaggttatcgctgtacttaaccagaaaggcgggtc
aggcaagacgaccatcgcaacccatctagcccgcgccctgcaactcgccggggccgatgttctgttagtc
gattccgatccccagggcagtgcccgcgattgggcggccgtgcgggaagatcaaccgctaaccgttgtcg
gcatcgaccgcccgacgattgaccgcgacgtgaaggccatcggccggcgcgacttcgtagtgatcgacgg
agcgccccaggcggcggacttggctgtgtccgcgatcaaggcagccgacttcgtgctgattccggtgcag
ccaagcccttacgacatatgggccaccgccgacctggtggagctggttaagcagcgcattgaggtcacgg
atggaaggctacaagcggcctttgtcgtgtcgcgggcgatcaaaggcacgcgcatcggcggtgaggttgc
cgaggcgctggccgggtacgagctgcccattcttgagtcccgtatcacgcagcgcgtgagctacccaggc
actgccgccgccggcacaaccgttcttgaatcagaacccgagggcgacgctgcccgcgaggtccaggcgc
tggccgctgaaattaaatcaaaactcatttgagttaatgaggtaaagagaaaatgagcaaaagcacaaac
acgctaagtgccggccgtccgagcgcacgcagcagcaaggctgcaacgttggccagcctggcagacacgc
cagccatgaagcgggtcaactttcagttgccggcggaggatcacaccaagctgaagatgtacgcggtacg
ccaaggcaagaccattaccgagctgctatctgaatacatcgcgcagctaccagagtaaatgagcaaatga
ataaatgagtagatgaattttagcggctaaaggaggcggcatggaaaatcaagaacaaccaggcaccgac
gccgtggaatgccccatgtgtggaggaacgggcggttggccaggcgtaagcggctgggttgcctgccggc
cctgcaatggcactggaacccccaagcccgaggaatcggcgtgagcggtcgcaaaccatccggcccggta
caaatcggcgcggcgctgggtgatgacctggtggagaagttgaaggccgcgcaggccgcccagcggcaac
gcatcgaggcagaagcacgccccggtgaatcgtggcaagcggccgctgatcgaatccgcaaagaatcccg
gcaaccgccggcagccggtgcgccgtcgattaggaagccgcccaagggcgacgagcaaccagatttttc
gttccgatgctctatgacgtgggcacccgcgatagtcgcagcatcatggacgtggccgttttccgtctgt
cgaagcgtgaccgacgagctggcgaggtgatccgctacgagcttccagacgggcacgtagaggtttccgc
agggccggccggcatggccagtgtgtgggattacgacctggtactgatggcggtttcccatctaaccgaa
tccatgaaccgataccgggaagggaagggagacaagcccggccgcgtgttccgtccacacgttgcggacg

tactcaagttctgccggcgagccgatggcggaaagcagaaagacgacctggtagaaacctgcattcggtt
aaacaccacgcacgttgccatgcagcgtacgaagaaggccaagaacggccgcctggtgacggtatccgag
ggtgaagccttgattagccgctacaagatcgtaaagagcgaaaccgggcggccggagtacatcgagatcg
agctagctgattggatgtaccgcgagatcacagaaggcaagaacccggacgtgctgacggttcaccccga
ttactttttgatcgatcccggcatcggccgttttctctaccgcctggcacgccgcgccgcaggcaaggca
gaagccagatggttgttcaagacgatctacgaacgcagtggcagcgccggagagttcaagaagttctgtt
tcaccgtgcgcaagctgatcgggtcaaatgacctgccggagtacgatttgaaggaggaggcggggcaggc
tggcccgatcctagtcatgcgctaccgcaacctgatcgagggcgaagcatccgccggttcctaatgtacg
gagcagatgctagggcaaattgccctagcaggggaaaaaggtcgaaaaggcctctttcctgtggatagca
cgtacattgggaacccaaagccgtacattgggaaccggaacccgtacattgggaacccaaagccgtacat
tgggaaccggtcacacatgtaagtgactgatataaaagagaaaaaaggcgatttttccgcctaaaactct
ttaaaacttattaaaactcttaaaacccgcctggcctgtgcataactgtctggccagcgcacagccgaag
agctgcaaaaagcgcctacccttcggtcgctgcgctccctacgccccgccgcttcgcgtcggcctatcgc
ggccgctggccgctcaaaaatggctggcctacggccaggcaatctaccagggcgcggacaagccgcgccg
tcgccactcgaccgccggcgcccacatcaaggcaccctgcctcgcgcgtttcggtgatgacggtgaaaac
ctctgacacatgcagctcccggaaacggtcacagcttgtctgtaagcggatgccgggagcagacaagccc
gtcagggcgcgtcagcgggtgttggcgggtgtcggggcgcagccatgacccagtcacgtagcgatagcgg
agtgtatactggcttaactatgcggcatcagagcagattgtactgagagtgcaccatatgcggtgtgaaa
taccgcacagatgcgtaaggagaaaataccgcatcaggcgctcttccgcttcctcgctcactgactcgct
gcgctcggtcgttcggctgcggcgagcggtatcagctcactcaaaggcggtaatacggttatccacagaa
tcaggggataacgcaggaaagaacatgtgagcaaaaggccagcaaaaggccaggaaccgtaaaaaggccg
cgttgctggcgtttttccataggctccgcccccctgacgagcatcacaaaaatcgacgctcaagtcagag
gtggcgaaacccgacaggactataaagataccaggcgtttccccctggaagctccctcgtgcgctctcct
gttccgaccctgccgcttaccggatacctgtccgcctttctcccttcgggaagcgtggcgctttctcata
gctcacgctgtaggtatctcagttcggtgtaggtcgttcgctccaagctgggctgtgtgcacgaacccc
cgttcagcccgaccgctgcgccttatccggtaactatcgtcttgagtccaacccggtaagacacgactta
tcgccactggcagcagccactggtaacaggattagcagagcgaggtatgtaggcggtgctacagagttct
tgaagtggtggcctaactacggctacactagaaggacagtatttggtatctgcgctctgctgaagccagt
taccttcggaaaaagagttggtagctcttgatccggcaaacaaaccaccgctggtagcggtggtttttt
gtttgcaagcagcagattacgcgcagaaaaaaaggatctcaagaagatcctttgatcttttctacggggt
ctgacgctcagtgctgatgaatcccctaatgattttatcaaaatcattaagttaaggtagatacacatc
ttgtcatatgatcaaatggtttcgccaaaaatcaataatcagacaacaaaatgtgcgaactcgatatttt
acacgactctctttaccaattctgccccgaattacacttaaaacgactcaacagcttaacgttggcttgc
cacgccttacttgactgtaaaactctcactcttaccgaacttggccgtaacctgccaaccaaagcgagaa
caaaacataacatcaaacgaatcgaccgattgttaggtaatcgtcacctccacaaagagcgactcgctgt
ataccgttggcatgctagctttatctgttcgggcaatacgatgcccattgtacttgttgactggtctgat
atccgtgagcaaaaacggcttatggtattgcgagcttcagtcgcactacacggtcgttctgttactcttt
atgagaaagcgttcccgctttcagagcaatgttcaaagaaagctcatgaccaatttctagccgaccttgc
gagcattctaccgagtaacaccacaccgctcattgtcagtgatgctggctttaaagtgccatggtataaa
tccgttgagaagctgggttggtactggttaagtcgagtaagaggaaaagtacaatatgcagacctaggag
cggaaaactggaaacctatcagcaacttacatgatatgtcatctagtcactcaaagactttaggctataa
gaggctgactaaaagcaatccaatctcatgccaaattctattgtataaatctcgctctaaaggccgaaaa
aatcagcgctcgacacggactcattgtcaccacccgtcacctaaaatctactcagcgtcggcaaaggagc
catggattctagcaactaacttacctgttgaaattcgaacacccaaacaacttgttaatatctattcgaa
gcgaatgcagattgaagaaccttccgagacttgaaaagtcctgcctacggactaggcctacgccatagc
cgaacgagcagctcagagcgttttgatatcatgctgctaatcgccctgatgcttcaactaacatgttggc
ttgcgggcgttcatgctcagaaacaaggttgggacaagcacttccaggctaacacagtcagaatcgaaa
cgtactctcaacagttcgcttaggcatggaagttttgcggcattctggctacacaataacaaggaagac
ttactcgtggctgcaaccctactagctcaaaatttattcacacatggttacgctttggggaaattatgag
gggatctctcagcgctcagtggaacgaaaactcacgttaagggattttggtcatgcattctaggtactaa
aacaattcatccagtaaaatataatattttattttctcccaatcaggcttgatccccagtaagtcaaaaa
atagctcgacatactgttcttccccgatatcctccctgatcgaccggacgcagaaggcaatgtcatacca

cttgtccgccctgccgcttctcccaagatcaataaagccacttactttgccatctttcacaaagatgttg
ctgtctcccaggtcgccgtgggaaaagacaagttcctcttcgggcttttccgtctttaaaaaatcataca
gctcgcgcggatctttaaatggagtgtcttcttcccagttttcgcaatccacatcggccagatcgttatt
cagtaagtaatccaattcggctaagcggctgtctaagctattcgtatagggacaatccgatatgtcgatg
gagtgaaagagcctgatgcactccgcatacagctcgataatcttttcagggctttgttcatcttcatact
cttccgagcaaaggacgccatcggcctcactcatgagcagattgctccagccatcatgccgttcaaagtg
caggacctttggaacaggcagctttccttccagccatagcatcatgtccttttcccgttccacatcatag
gtggtccctttataccggctgtccgtcatttttaaatataggttttcattttctcccaccagcttatata
ccttagcaggagacattccttccgtatcttttacgcagcggtatttttcgatcagttttttcaattccgg
tgatattctcatttagccatttattatttccttcctcttttctacagtatttaaagataccccaagaag
ctaattataacaagacgaactccaattcactgttccttgcattctaaaaccttaaataccagaaaacagc
tttttcaaagttgttttcaaagttggcgtataacatagtatcgacggagccgattttgaaaccgcggtga
tcacaggcagcaacgctctgtcatcgttacaatcaacatgctaccctccgcgagatcatccgtgtttcaa
acccggcagcttagttgccgttcttccgaatagcatcggtaacatgagcaaagtctgccgccttacaacg
gctctcccgctgacgccgtcccggactgatgggctgcctgtatcgagtggtgattttgtgccgagctgcc
ggtcggggagctgttggctggctgg

HIGHLIGHTED SEQUENCES:

Left T-DNA border
nos terminator
hygromycin phosphotransferase II coding sequence
nos promoter
CaMV 35S terminator
eGFP coding sequence
CaMV 35S promoter
Right T-DNA border
pVS1 replicon region
neomycin phosphotransferase III (bacterial resistance marker)

| Primer name | Sequence | Template | Enzyme ends | Purpose |
|---|---|---|---|---|
| m2_specR_AscI_amp-F | GGATTTAACTGGCGCGCCTGCTTTTGATCGGTACCA | pVS-rec (S) | AscI | Amplifying the spectinomycin resistance marker from pVS-rec (S) for digestion with AscI and XmaI enzymes followed by ligation into same digested pAGROm2_111, yielding pAGROm2_002 |
| m2_specR_XmaI_amp-R | GGATTTGGATCCCGGGTAGGGCTTATTATGCACGCTT | pVS-rec (S) | XmaI | Amplifying the spectinomycin resistance marker from pVS-rec (S) for digestion with AscI and XmaI enzymes followed by ligation into same digested pAGROm2_111, yielding pAGROm2_002 |
| m4_sacB_only_ol-F | TGGCAATTGCGACCGGCGCGCCGTCGACTTGAGCTCGCGACA( | N/A (used for annealing with par | N/A | Using Gibson assembly to patch an annealed dsOligo pair into a SalI and SacI-digested pAGROm4_210 backbone fragment, yielding pAGROm4_010 |
| m4_sacB_only_ol-R | GTGAACGGCAGGTATATGTGTCGCGAGCTCAAGTCGACGGCGC | N/A (used for annealing with par | N/A | Using Gibson assembly to patch an annealed dsOligo pair into a SalI and SacI-digested pAGROm4_210 backbone fragment, yielding pAGROm4_010 |
| sfGFP_seq-F | GGATGGTGATGTCAACGGT | pDisarm or pDelete constructs a | N/A | Colony PCR for verifying primary OR secondary recombination status |
| sfGFP_seq-R | GCAGAACCATATGATCGCGT | pDisarm or pDelete constructs a | N/A | Colony PCR for verifying primary OR secondary recombination status |
| specR_seq-F | CTGGTTACGGTGACCGTAAG | pDisarm or pDelete constructs a | N/A | Colony PCR for verifying primary OR secondary recombination status |
| specR_seq-R | TCAGCAAGATAGCCAGATCAATG | pDisarm or pDelete constructs a | N/A | Colony PCR for verifying primary OR secondary recombination status |
| sacB_seq-F | GGAAACATACGGCATTTCCC | pDisarm or pDelete constructs a | N/A | Colony PCR for verifying exclusively primary recombination status |
| sacB_seq-R | GCTTGAGGTACAGCGAAG | pDisarm or pDelete constructs a | N/A | Colony PCR for verifying exclusively primary recombination status |
